## Supplementary Materials for "Three Open Questions in Polygenic Score Portability"

### Contents

1 **Sample characteristics and genetic distance** 8

2 **The relationship between $F_{st}$ and genotype matrix PCs** 8

3 **Sensitivity analyses for major portability trends** 9

S5 Trends of portability vary across traits and measures, for other biomarkers. . 19

S6 Full range of individual level prediction error for anthropometric measurements. 20

S7 Full range of individual level prediction error for white blood cell-related traits. 21

S8 Full range of individual level prediction error for red blood cell-related traits. 22

S9 Full range of individual level prediction error for other biomarkers. . . . . . . 23

S10 Divergence of group- and individual-level prediction accuracy for anthropometric measurements. . . . . . . . . . . . . . . . . . . . . . . . . . . . . 24
S11 Divergence of group- and individual-level prediction accuracy for white blood cell-related traits. . . . . . . . . . . . . . . . . . . . . . . . . . . . . . . . . 25
S12 Divergence of group- and individual-level prediction accuracy for red blood cell-related traits. . . . . . . . . . . . . . . . . . . . . . . . . . . . . . . . . 26
S13 Divergence of group- and individual-level prediction accuracy for other biomarkers. . . . . . . . . . . . . . . . . . . . . . . . . . . . . . . . . . . . . . 27
S14 Mean trends in individual-level prediction accuracy by different measures for anthropometric measurements. . . . . . . . . . . . . . . . . . . . . . . . . . 28
S15 Mean trends in individual-level prediction accuracy by different measures for white blood cell-related traits. . . . . . . . . . . . . . . . . . . . . . . . . 29
S16 Mean trends in individual-level prediction accuracy by different measures for red blood cell-related traits. . . . . . . . . . . . . . . . . . . . . . . . . . 30
S17 Mean trends in individual-level prediction accuracy by different measures for other biomarkers. . . . . . . . . . . . . . . . . . . . . . . . . . . . . . . . 31
S18 Individual-level prediction error explained by different measures for anthropometric measurements. . . . . . . . . . . . . . . . . . . . . . . . . . . . . 32
S19 Individual-level prediction error explained by different measures for white blood cell-related traits. . . . . . . . . . . . . . . . . . . . . . . . . . . . . 33
S20 Individual-level prediction error explained by different measures for red blood cell-related traits. . . . . . . . . . . . . . . . . . . . . . . . . . . . . . . . 34
S21 Individual-level prediction error explained by different measures for other biomarkers. . . . . . . . . . . . . . . . . . . . . . . . . . . . . . . . . . . . 35
S22 Mean heterozygosity of SNPs, stratified by effect size. . . . . . . . . . . . . . 36
S23 Squared allelic effect estimate of SNPs. . . . . . . . . . . . . . . . . . . . . . 37

S24 Heritability explained by index SNPs. . . . . . . . . . . . . . . . . . . . . 38
S25 Mean Townsend Deprivation Index ± SE in each bin by genetic distance. . . 39
S26 Average total household income before tax in each bin by genetic distance. . 40
S27 Mean age ± SE in each bin by genetic distance. . . . . . . . . . . . . . . . . 41
S28 Sex composition across bins of genetic distance. . . . . . . . . . . . . . . . 42
S29 Participants' country of residence across bins of genetic distance. . . . . . . . 43
S30 Correlation between $F_{st}$ and PC 1. . . . . . . . . . . . . . . . . . . . . . 44
S31 Correlation between $F_{st}$ and PC 2. . . . . . . . . . . . . . . . . . . . . . 45
S32 Correlation between $F_{st}$ and PC 3. . . . . . . . . . . . . . . . . . . . . . 46
S33 Correlation between $F_{st}$ and PC 40. . . . . . . . . . . . . . . . . . . . . 47
S34 Trends of portability vary across traits and measures, with confidence intervals, for anthropometric measurements. . . . . . . . . . . . . . . . . . . . . 48
S35 Trends of portability vary across traits and measures, with confidence intervals, for white blood-cell related traits. . . . . . . . . . . . . . . . . . . . 49
S36 Trends of portability vary across traits and measures, with confidence intervals, for red blood cell-related traits. . . . . . . . . . . . . . . . . . . . . 50
S37 Trends of portability vary across traits and measures, with confidence intervals, for other biomarkers. . . . . . . . . . . . . . . . . . . . . . . . . . 51
S38 Group level portability, using more bins, for anthropometric measurements. . 52
S39 Group level portability, using more bins, for white blood-cell related traits. . 53
S40 Group level portability, using more bins, for red blood cell-related traits. . . 54
S41 Group level portability, using more bins, for other biomarkers. . . . . . . . . 55
S42 Trends of portability visualized with generalized additive model (GAM) fit, for anthropometric measurements. . . . . . . . . . . . . . . . . . . . . . . . 56
S43 Trends of portability visualized with generalized additive model (GAM) fit, for white blood-cell related traits. . . . . . . . . . . . . . . . . . . . . . . . 57

S44 Trends of portability visualized with generalized additive model (GAM) fit, for red blood cell-related traits. . . . . . . . . . . . . . . . . . . . . . . . . . 58
S45 Trends of portability visualized with generalized additive model (GAM) fit, for other biomarkers. . . . . . . . . . . . . . . . . . . . . . . . . . . . . . . 59
S46 Trends of portability visualized using 12 knots in cubic line fits, for anthropometric measurements. . . . . . . . . . . . . . . . . . . . . . . . . . . . 60
S47 Trends of portability visualized using 12 knots in cubic line fits, for white blood-cell related traits. . . . . . . . . . . . . . . . . . . . . . . . . . . . 61
S48 Trends of portability visualized using 12 knots in cubic line fits, for red blood cell-related traits. . . . . . . . . . . . . . . . . . . . . . . . . . . . . . . 62
S49 Trends of portability visualized using 12 knots in cubic line fits, for other biomarkers. . . . . . . . . . . . . . . . . . . . . . . . . . . . . . . . . . . 63
S50 Trends of portability when including additional GWAS covariates, for anthropometric measurements. . . . . . . . . . . . . . . . . . . . . . . . . . . . 64
S51 Trends of portability when including additional GWAS covariates, for white blood-cell related traits. . . . . . . . . . . . . . . . . . . . . . . . . . . . 65
S52 Trends of portability when including additional GWAS covariates, for red blood cell-related traits. . . . . . . . . . . . . . . . . . . . . . . . . . . . 66
S53 Trends of portability when including additional GWAS covariates, for other biomarkers. . . . . . . . . . . . . . . . . . . . . . . . . . . . . . . . . . . 67
S54 Trends of portability with GWAS run using *regenie*, for anthropometric measurements. . . . . . . . . . . . . . . . . . . . . . . . . . . . . . . . . . . 68
S55 Trends of portability with GWAS run using *regenie*, for white blood-cell related traits. . . . . . . . . . . . . . . . . . . . . . . . . . . . . . . . . . . 69
S56 Trends of portability with GWAS run using *regenie*, for red blood cell-related traits. . . . . . . . . . . . . . . . . . . . . . . . . . . . . . . . . . . . . 70

S57 Trends of portability with GWAS run using *regenie*, for other biomarkers. . . 71
S58 Trends of portability with PGS constructed using *PRS-CS*, for anthropometric measurements. . . . . . . . . . . . . . . . . . . . . . . . . . . . . . . 72
S59 Trends of portability with PGS constructed using *PRS-CS*, for white blood-cell related traits. . . . . . . . . . . . . . . . . . . . . . . . . . . . . . . 73
S60 Trends of portability with PGS constructed using *PRS-CS*, for red blood cell-related traits. . . . . . . . . . . . . . . . . . . . . . . . . . . . . . . 74
S61 Trends of portability with PGS constructed using *PRS-CS*, for other biomarkers. 75
S62 A comparison of portability trends, with genetic distance defined in two different ways, for anthropometric measurements. . . . . . . . . . . . . . . . . 76
S63 A comparison of portability trends, with genetic distance defined in two different ways, for white blood-cell related traits. . . . . . . . . . . . . . . . . 77
S64 A comparison of portability trends, with genetic distance defined in two different ways, for red blood cell-related traits. . . . . . . . . . . . . . . . . 78
S65 A comparison of portability trends, with genetic distance defined in two different ways, for other biomarkers. . . . . . . . . . . . . . . . . . . . . . . 79
S66 Variance in disease PGS as a function of genetic distance. . . . . . . . . . . 80
S67 Portability as a function of both genetic distance and Townsend Deprivation Index, for anthropometric measurements. . . . . . . . . . . . . . . . . . . 81
S68 Portability as a function of both genetic distance and Townsend Deprivation Index, for white blood-cell related traits. . . . . . . . . . . . . . . . . . . 82
S69 Portability as a function of both genetic distance and Townsend Deprivation Index, for red blood cell-related traits. . . . . . . . . . . . . . . . . . . 83
S70 Portability as a function of both genetic distance and Townsend Deprivation Index, for other biomarkers. . . . . . . . . . . . . . . . . . . . . . . . . 84

S71 Measuring "genetic distance" from the GWAS sample, with White British individuals included in the prediction set. . . . . . . . . . . . . . . . . . . . . 85
S72 A comparison of portability trends with different choices of the GWAS and prediction samples, for anthropometric measurements. . . . . . . . . . . . . . 86
S73 A comparison of portability trends with different choices of the GWAS and prediction samples, for white blood-cell related traits. . . . . . . . . . . . . . 87
S74 A comparison of portability trends with different choices of the GWAS and prediction samples, for red blood cell-related traits. . . . . . . . . . . . . . . 88
S75 A comparison of portability trends with different choices of the GWAS and prediction samples, for other biomarkers. . . . . . . . . . . . . . . . . . . . . 89
S76 Zoomed-in portability trends, with WB in the prediction set, for anthropometric measurements. . . . . . . . . . . . . . . . . . . . . . . . . . . . . . 90
S77 Zoomed-in portability trends, with WB in the prediction set, for white blood-cell related traits. . . . . . . . . . . . . . . . . . . . . . . . . . . . . . . 91
S78 Zoomed-in portability trends, with WB in the prediction set, for red blood cell-related traits. . . . . . . . . . . . . . . . . . . . . . . . . . . . . . . . 92
S79 Zoomed-in portability trends, with WB in the prediction set, for other biomarkers. . . . . . . . . . . . . . . . . . . . . . . . . . . . . . . . . . . . . . 93

### 1 Sample characteristics and genetic distance

We examined the distribution of several sample characteristics in the GWAS set and in the prediction set. These characteristics included: Townsend Deprivation Index (UKB data field 189; **Fig. S25**), average total household income before tax (UKB data field 738; **Fig. S26**), sex (UKB data field 22001; **Fig. S28**), age (UKB data field 21022), and country of residence (converted from UKB data field 54, into England, Scotland, or Wales; **Fig. S29**).

For Townsend Deprivation Index (**Fig. S25**) and age (**Fig. S27**), we plotted the mean $\pm$ SE in each bin, along with the values in the GWAS set for reference. To make the SE in the GWAS comparable with the SE in each bin, we calculated it as: $\frac{SD_{GWAS}}{\sqrt{\bar{n}_{pred}}}$, where $\bar{n}_{pred}$ represents the mean sample size across all bins in the prediction set (278). For average total household income before tax, we calculated the frequency of each category in each bin and plotted the cumulative frequency, along with the frequencies in the GWAS set for reference (**Fig. S26**). For sex, we calculated the frequency of females in each bin and also plotted the mean in the GWAS set (**Fig. S28**). For country, we first converted the assessment center to the corresponding country (England, Scotland, or Wales), then calculated the frequency of each category in each bin and plotted the cumulative frequency, along with the frequencies in the GWAS set for reference (**Fig. S29**).

### 2 The relationship between $F_{st}$ and genotype matrix PCs

Using the same 10,000 randomly selected individuals from the prediction sample, we examined the correlation between $F_{st}$ and invidividual coordinates along specific PCs: PCs 1 (**Fig. S30**), 2 (**Fig. S31**), 3 (**Fig. S32**), and 40 (**Fig. S33**). All PCs are taken from UKB field 22009. We color-coded the individuals by self-reported race (UKB field 21000).

### 3 Sensitivity analyses for major portability trends

We performed several sensitivity analyses to test the sensitivity of our conclusions about portability trends to various choices we made in the analyses presented in the main text.

**Binning individuals into 500 bins**. To examine the sensitivity of our analyses to binning parameter choices, we perform the analysis with a different binning. We binned individuals in the prediction set into 500 bins of 139 individuals each, instead of 250 bins of 278 individuals each. In **Figs. S38-S41**, we show relative prediction accuracy trends using the mean prediction accuracy in the 50 bins with median genetic distances that are closest to the mean genetic distance for GWAS individuals as the baseline value. Comparing the two binning choices, we did not notice any change in the portability trends for any of the quantitative traits analyzed.

**Additional GWAS covariates**. We examined whether additional GWAS covariates, not included in the main text analysis, impact our conclusions. We performed polygenic scoring based on new GWAS that included assessment center (UKB data field 54) and genotype array (UKB data field 22000) as additional covariates. We converted these two categorical variables into multiple binary variables using one-hot encoding, and removed one column from either of the variables to prevent multicollinearity). We followed the same procedures as described in **Methods** for clumping and thresholding, PGS construction and analysis of prediction accuracy trends at the group and individual levels for 15 quantitative traits (**Figs. S50-S53**). Adding these covariates did not lead to any prominent change in the portability trends for most traits; for lymphocyte and monocyte counts, the addition of these covariates generally resulted in better prediction at large genetic distances.

**Performing GWAS with *regenie***. To test the sensitivity of our three conclusions to the GWAS methods we used, we re-ran GWAS for all 15 quantitative traits we analyzed using the same covariates as described in **Methods**, but using *regenie*[21]. Then we followed the same procedures for clumping and thresholding with *PLINK*, as described in **Methods**. After that, we computed the PGS prediction accuracy at the group and individual levels (**Figs. S54-S57**). We saw no major change in the portability trends for most traits. For lymphocyte count, however, prediction accuracy at the group level was noticeably higher in some bins.

**Constructing PGS with *PRS-CS***. To examine the sensitivity of our conclusions to the choice of method for PGS construction, we used an approach distinct from clumping and thresholding—*PRS-CS*, Using the `--score` flag in this software[10]. PRS-CS applies Bayesian shrinkage to GWAS-based allelic effect estimates. It also aims to consider LD patterns more rigorously and holistically, rather than via the clumping and thresholding heuristic. To run *PRS-CS*, we used the same GWAS summary statistics as before and the LD reference panel Ge *et al.* constructed using the UKB EUR super-population data. For the other optional parameters $a$, $b$, and $\phi$, we used the default values. We computed the PGS prediction accuracy at the group and individual levels for 15 quantitative traits (**Figs. S58-S61**). With *PRS-CS*, for weight and BMI at the group level, the peak at intermediate genetic distances became less prominent. For lymphocyte count and white blood cell count, the decrease in prediction at the group level became more gradual. However, the general trends of portability of other traits did not change.

**Defining genetic distance using 16 instead of 40 PCs**. Previous research puts into question the optimal number of genotype matrix PCs that should be considered, both generally in genetic analyses and for specific tasks[53,32,42]. In the main text, we use a genetic distance metric of an eigenvalue-weighted Eucledean distance in PCA space. The PCA is the pre-computed PCA of the genotype matrix of the UK Biobank sample, and the genetic distance from the GWAS sample is the distance of a given individual's PC coordinates from the centroid of GWAS sample individuals (**Methods**). Here, we repeated this calculation using the first 16 PCs provided by UKB (UKB data field 22009) rather than the first 40.

Among all samples, the Pearson correlation between genetic distances calculated from 40 PCs and from 16 PCs is 0.9999994 and the Spearman correlation is 0.9984838. We computed PGS prediction accuracy at the group and individual levels for 15 quantitative traits as before. Across all 15 quantitative traits, there were no noticeable changes in portability trends (**Figs. S62-S65**).

**Continuous trend visualization**. To examine whether our conclusions are sensitive to continuous function fits we used for visualizations, We first examined the effect of a change in the cubic spline parameters. Namely, instead of 8 knots, we used 12 (**Figs. S46-S49**). Using 12 knots, the fit appears more flexible, but the general trends of portability discussed in the main text did not change.

We also tested a different functional form for the fit, a generalized additive model (GAM). We used `gam()` from the R package mgcv[51] instead of a cubic spline (**Figs. S42-S45**). For comparison, we placed the 8 knots at the same positions for the GAM model. The GAM fit appears to be smoother at the group level for most traits, but have the same general trends as the cubic spline fit.

**Portability trends stratified by Townsend Deprivation Index**. In the main text analysis, we discuss the observation that, across traits, genetic distance and Townsend Deprivation Index[44] explain comparable fractions of variation in individual-level prediction accuracy (**3B**). Here, we examine the two predictors' joint effect. In particular, we stratify individuals by Townsend Deprivation Index and visualize trends of prediction accuracy as a function of genetic distance in each SES stratum independently.

We performed this analysis at both the group and individual levels. At the group level, we stratified the prediction individuals into 2 equal-size strata of Townsend Deprivation Index. At the individual level, we binned individuals into 5 equal-size strata. In each stratum independently, we then followed the analysis as otherwise described in **Methods** to calculate prediction accuracy at the group and individual levels. The results are shown in

**Figs. S67-S70**.

**Including White British individuals in the prediction set**. In the main text, we opted to include all White British individuals in the GWAS sample and none in the prediction set. Here, we visit the sensitivity of our conclusions to this choice by reducing the GWAS sample size to 300,000 randomly selected White British individuals and adding the remaining 36,923 White British individuals to the prediction sample (now totaling 106,423 individuals). We otherwise followed the same procedures as described in **Methods**, verifying that, across 10,000 individuals from the prediction set, genetic distance (calculated with 40 pre-computed PCs provided by the UKB) remained highly correlated with $F_{st}$ between the new GWAS sample and the individual. The distribution of genetic distance with this new choice of GWAS sample and prediction sample is shown in **Fig. S71**.

We measure the prediction accuracy at the group and individual levels using the same procedures as described in **Methods**. Since the prediction set became larger, we binned the individuals into 382 bins of 279 individuals each at the group level (compared to 250 bins of 278 individuals each in the main text analysis). Across traits, we observed similar qualitative trends as we did in the main text analysis (**Figs. S72-S75**. Notably, in short genetic distances ($< 5$) the group-level prediction accuracy tended to be near-constant (**Figs. S76-S79**).

| Phenotype | SNP $h^2$ | Mean squared prediction error (MSE) | Variance of residualized phenotype ($Var[Z]$) | $1 - \frac{MSE}{Var[Z]}$ | Prediction accuracy (partial $R^2$) |
|---|---|---|---|---|---|
| Height | 0.4852 | 31.8859 | 41.5853 | 0.2332 | 0.2473 |
| Cystatin C level | 0.3214 | 0.0319 | 0.0333 | 0.0426 | 0.0632 |
| Platelet count | 0.3079 | 2950.5930 | 3369.6670 | 0.1244 | 0.1365 |
| Mean corpuscular volume | 0.2667 | 16.5441 | 18.7806 | 0.1191 | 0.1303 |
| Weight | 0.2654 | 205.3049 | 217.1554 | 0.0546 | 0.0620 |
| Mean corpuscular hemoglobin | 0.2530 | 2.6206 | 2.9492 | 0.1114 | 0.1253 |
| BMI | 0.2482 | 23.7059 | 24.6588 | 0.0386 | 0.0442 |
| Red blood cell count | 0.2337 | 0.1061 | 0.1161 | 0.0859 | 0.0912 |
| Body fat percentage | 0.0472 | 41.7102 | 43.2307 | 0.0352 | 0.0419 |
| Monocyte count | 0.2305 | 0.03674 | 0.0380 | 0.0325 | 0.0870 |
| Triglyceride level | 0.2182 | 0.9468 | 1.0010 | 0.0542 | 0.0620 |
| Lymphocyte count | 0.2103 | 1.1740 | 1.1753 | 0.0011 | 0.0342 |
| White blood cell count | 0.1910 | 3.8944 | 4.0020 | 0.0269 | 0.0455 |
| Eosinophil count | 0.1840 | 0.0195 | 0.0204 | 0.0430 | 0.0505 |
| LDL cholesterol level | 0.0825 | 0.6270 | 0.6810 | 0.0792 | 0.0887 |

Table S1: Characteristics of continuous traits and PGS analyzed. SNP heritabilities (SNP $h^2$) are taken from the Neale Lab's UKB analysis[26]. For the 25 bins with a genetic distance most similar to the mean genetic distance of the GWAS group, we calculated the group-level prediction accuracy (partial genetic correlation of the PGS and the trait value), mean individual prediction error (squared prediction error), the variance of residualized phenotype, and the ratio between the two subtracted from one, as another measure of phenotype variance explained by the PGS close to the GWAS sample.

| **Phenotype** | **Number of ref. bins** | **Mean precision** | **Mean recall** |
|---|---|---|---|
| Type 2 diabetes | 25 | 0.0944 | 0.3302 |
| Asthma | 25 | 0.1076 | 0.3481 |

Table S2: Characteristics of binary traits and PGS analyzed. For the reference bins (bins with a genetic distance most similar to the mean genetic distance of the GWAS group), we provide the mean precision and recall values in this table.

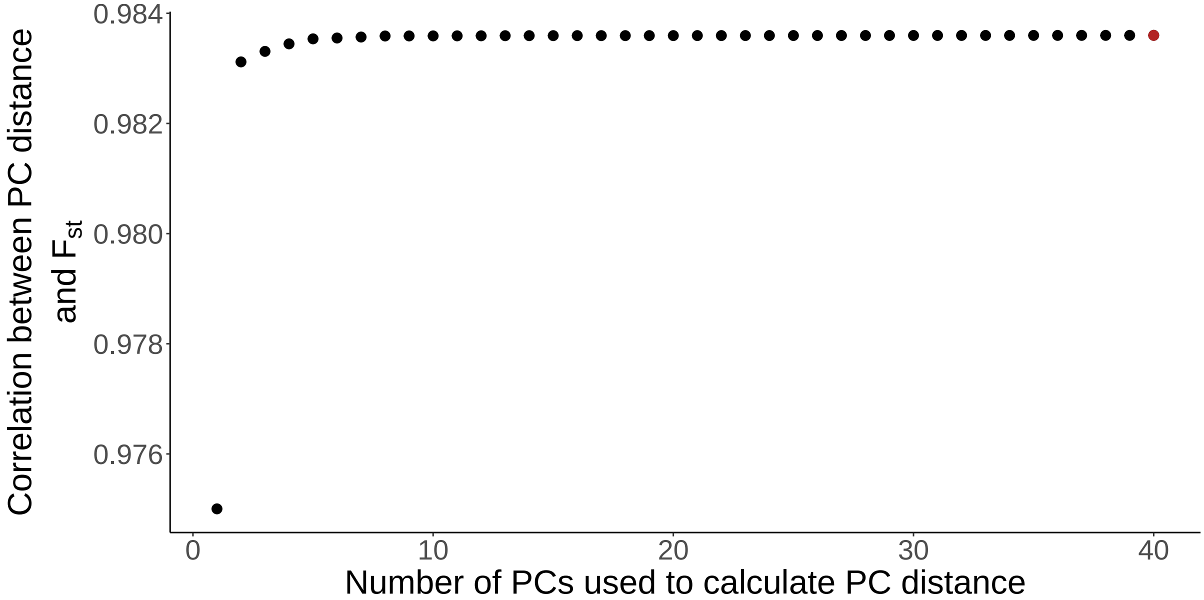

Figure S1: Correlation between PC distance and $F_{st}$. Shown are the Pearson correlations between PC distance using a given number of UKB PCs and $F_{st}$ between the GWAS sample and individuals. The correlations are calculated using a subset of 10,000 prediction sample individuals with a weighted PC distance greater than the weighted PC distance of 95% of the GWAS set (based on weighted PC distance calculated from the $K = 10$), as described in **Methods**. Using 40 PCs produces the highest correlation between PC distance and $F_{st}$ (red dot, $r > 0.98$).

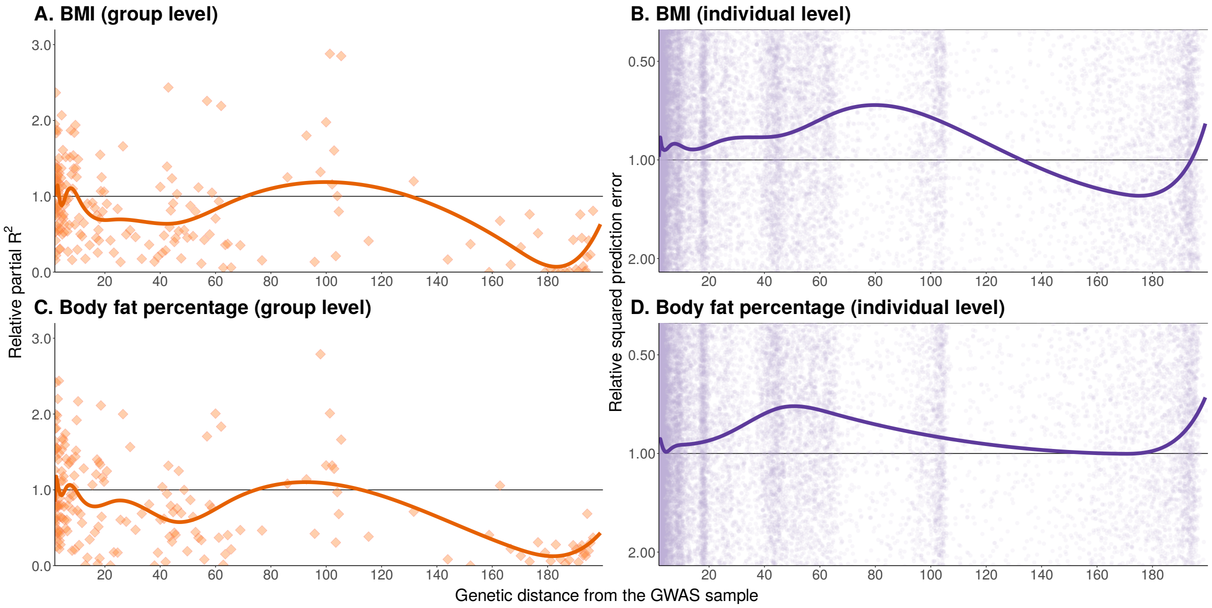

Figure S2: Trends of portability vary across traits and measures, for anthropometric measurements. This figure presents the same analysis as **Fig. 2** in the main text, but shows other anthropometric traits. At the group level (left panels), we measured prediction accuracy with the squared partial correlation between the PGS and the trait value in 250 bins of 278 individuals each. At the individual level (right panels), we measured prediction accuracy as the squared difference between the predicted phenotype and the true phenotype value. At both the group and individual levels, y-axis values show relative prediction accuracy, i.e. prediction accuracy divided by a baseline value. The baseline value is the mean prediction accuracy in 25 bins with median genetic distances that are closest to the mean genetic distance for GWAS individuals. **Table S1** details these baseline values for each trait. Curves show cubic spline fits, with 8 knots placed based on the density of data points. See **Figs. S6-S9** for plots showing the full ranges of individual-level prediction accuracy.

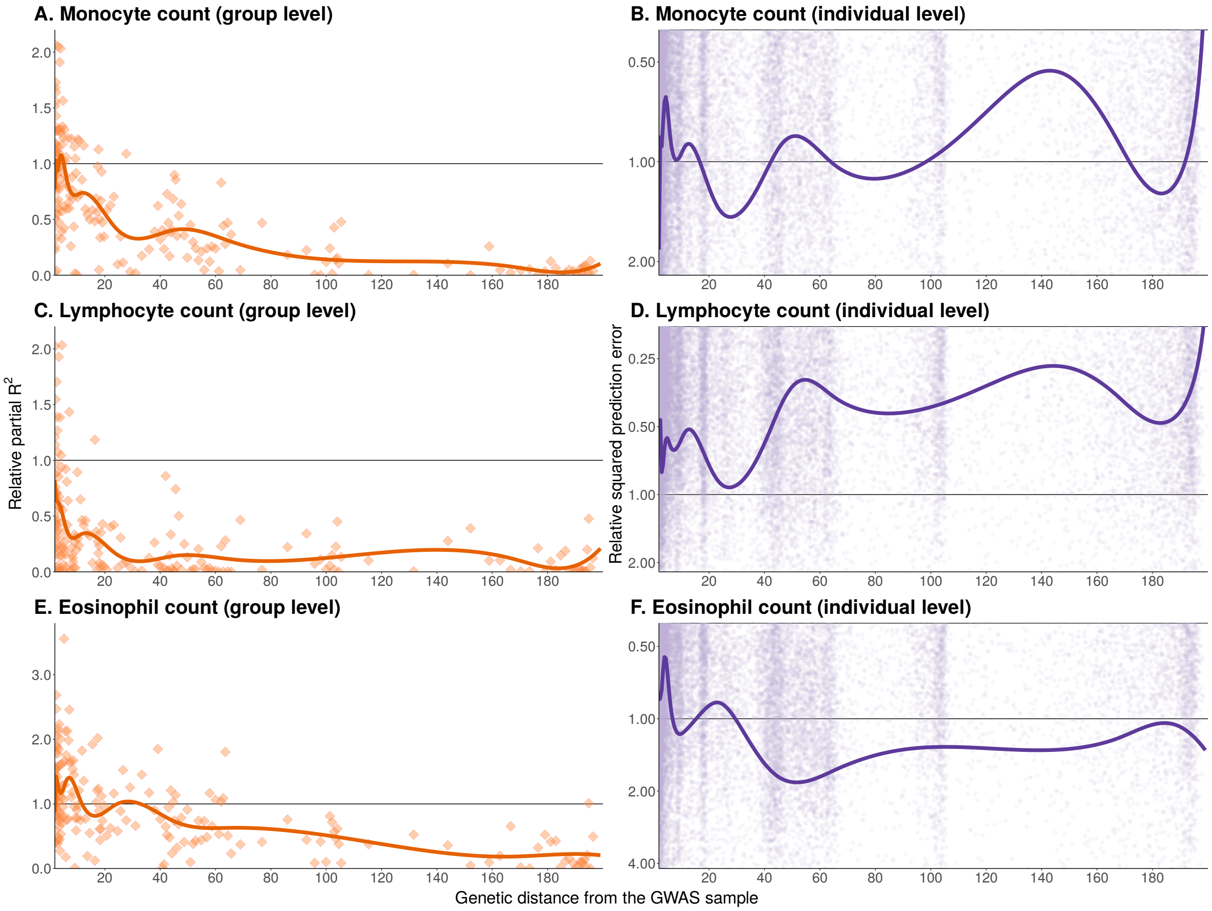

Figure S3: Trends of portability vary across traits and measures, for white blood-cell related traits. This figure presents the same analysis as **Fig. 2** in the main text, but shows other white blood cell-related traits. At the group level (left panels), we measured prediction accuracy with the squared partial correlation between the PGS and the trait value in 250 bins of 278 individuals each. At the individual level (right panels), we measured prediction accuracy as the squared difference between the predicted phenotype and the true phenotype value. At both the group and individual levels, y-axis values show relative prediction accuracy, i.e. prediction accuracy divided by a baseline value. The baseline value is the mean prediction accuracy in 25 bins with median genetic distances that are closest to the mean genetic distance for GWAS individuals. **Table S1** details these baseline values for each trait. Curves show cubic spline fits, with 8 knots placed based on the density of data points. See **Figs. S6-S9** for plots showing the full ranges of individual-level prediction accuracy.

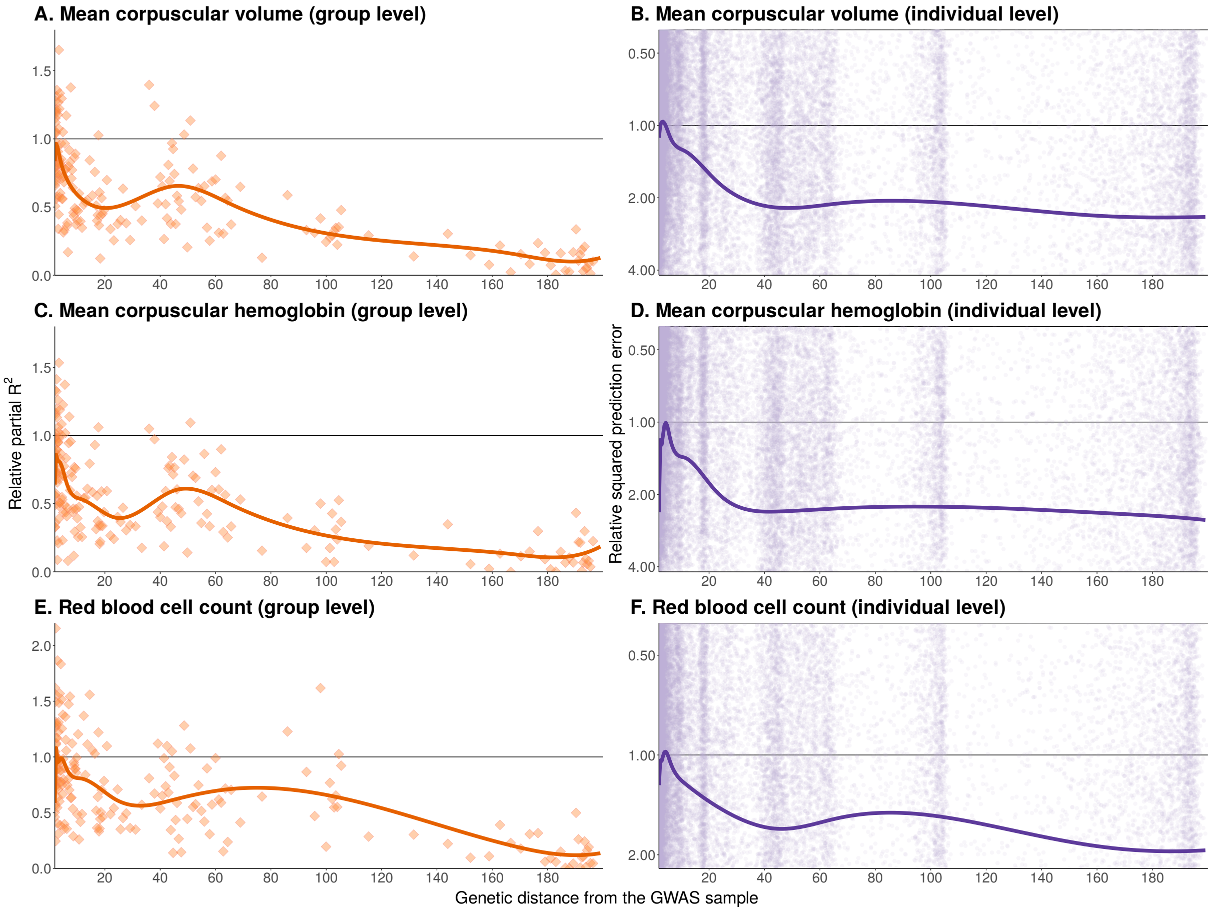

Figure S4: Trends of portability vary across traits and measures, for red blood-cell related traits. This figure presents the same analysis as **Fig. 2** in the main text, but shows other red blood cell-related traits. At the group level (left panels), we measured prediction accuracy with the squared partial correlation between the PGS and the trait value in 250 bins of 278 individuals each. At the individual level (right panels), we measured prediction accuracy as the squared difference between the predicted phenotype and the true phenotype value. At both the group and individual levels, y-axis values show relative prediction accuracy, i.e. prediction accuracy divided by a baseline value. The baseline value is the mean prediction accuracy in 25 bins with median genetic distances that are closest to the mean genetic distance for GWAS individuals. **Table S1** details these baseline values for each trait. Curves show cubic spline fits, with 8 knots placed based on the density of data points. See **Figs. S6-S9** for plots showing the full ranges of individual-level prediction accuracy.

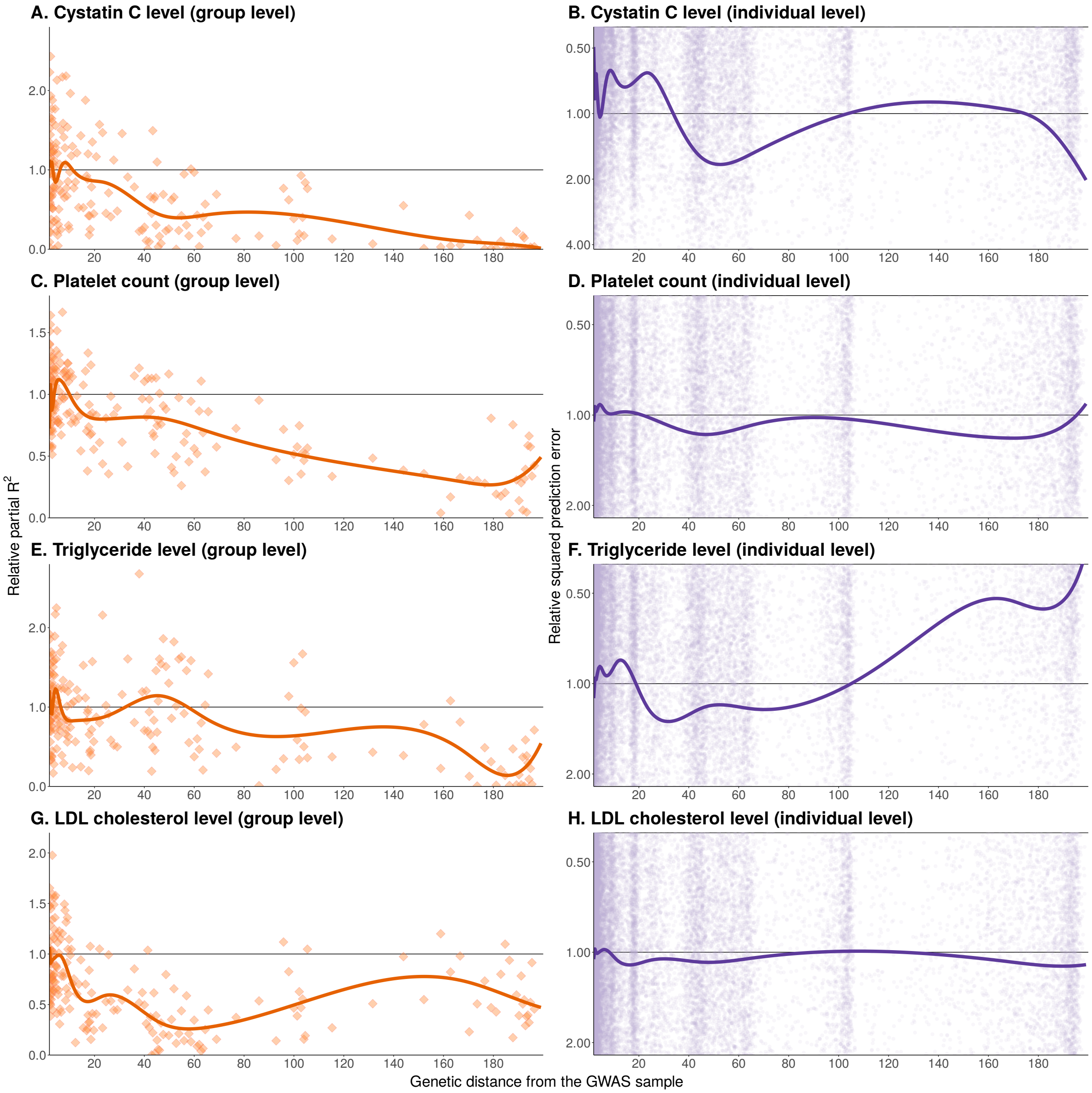

Figure S5: Trends of portability vary across traits and measures, for other biomarkers. This figure presents the same analysis as **Fig. 2** in the main text, but shows other biomarkers. At the group level (left panels), we measured prediction accuracy with the squared partial correlation between the PGS and the trait value in 250 bins of 278 individuals each. At the individual level (right panels), we measured prediction accuracy as the squared difference between the predicted phenotype and the true phenotype value. At both the group and individual levels, y-axis values show relative prediction accuracy, i.e. prediction accuracy divided by a baseline value. The baseline value is the mean prediction accuracy in 25 bins with median genetic distances that are closest to the mean genetic distance for GWAS individuals. **Table S1** details these baseline values for each trait. Curves show cubic spline fits, with 8 knots placed based on the density of data points. See **Figs. S6-S9** for plots showing the full ranges of individual-level prediction accuracy.

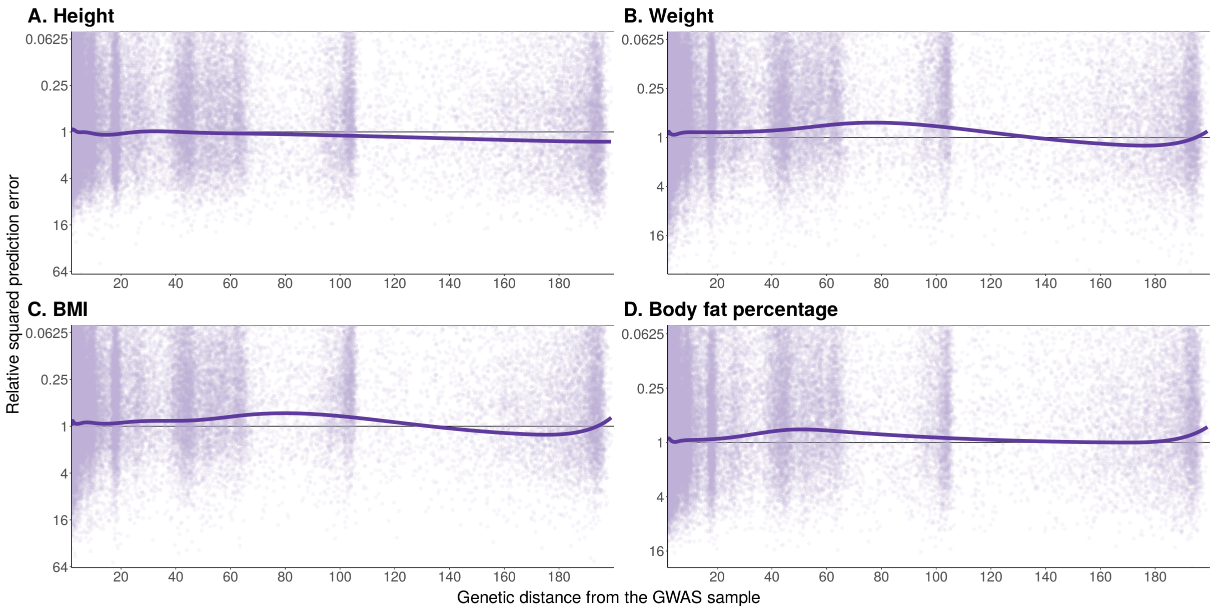

Figure S6: Full range of individual level prediction error for anthropometric measurements. This figure presents the same analysis as **Fig. 2** in the main text, but shows the full range of prediction error for other anthropometric traits. At the individual level, we measured prediction accuracy as the squared difference between the predicted phenotype and the true phenotype value. The y-axis values show relative prediction accuracy, i.e. prediction accuracy divided by a baseline value. The baseline value is the mean prediction accuracy in 25 bins with median genetic distances that are closest to the mean genetic distance for GWAS individuals. **Table S1** details these baseline values for each trait. Curves show cubic spline fits, with 8 knots placed based on the density of data points. See **Figs. S2-S5** for plots showing the zoomed-in version of individual-level prediction accuracy.

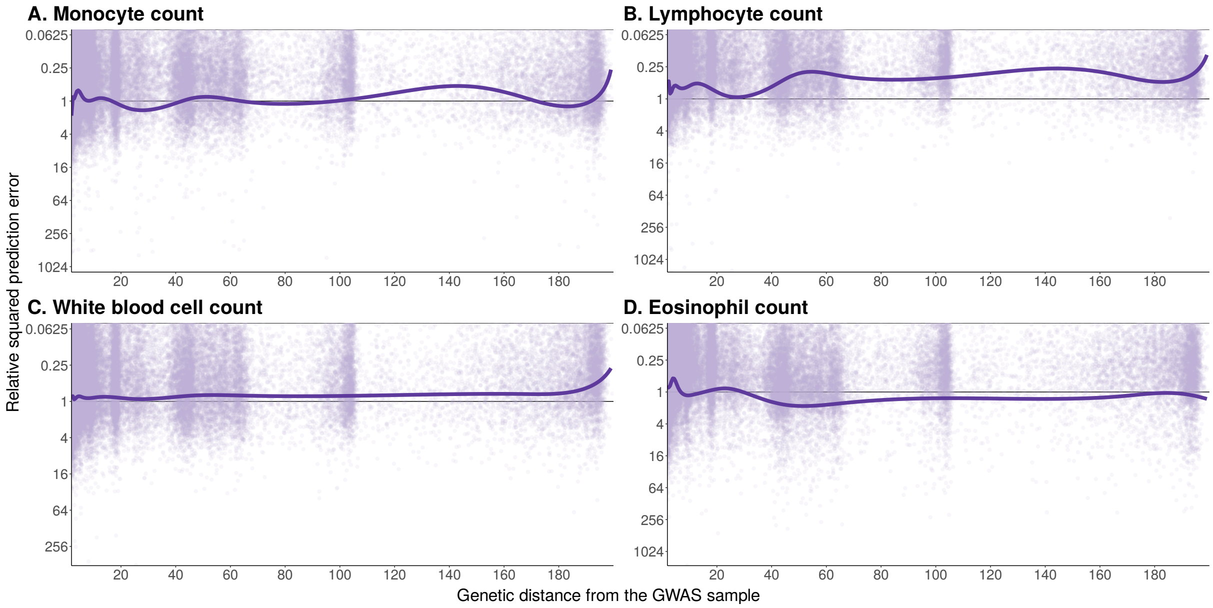

Figure S7: Full range of individual level prediction error for white blood cell-related traits. This figure presents the same analysis as **Fig. 2** in the main text, but shows the full range of prediction error for white blood cell-related traits. At the individual level, we measured prediction accuracy as the squared difference between the predicted phenotype and the true phenotype value. The y-axis values show relative prediction accuracy, i.e. prediction accuracy divided by a baseline value. The baseline value is the mean prediction accuracy in 25 bins with median genetic distances that are closest to the mean genetic distance for GWAS individuals. **Table S1** details these baseline values for each trait. Curves show cubic spline fits, with 8 knots placed based on the density of data points. See **Figs. S2-S5** for plots showing the zoomed-in version of individual-level prediction accuracy.

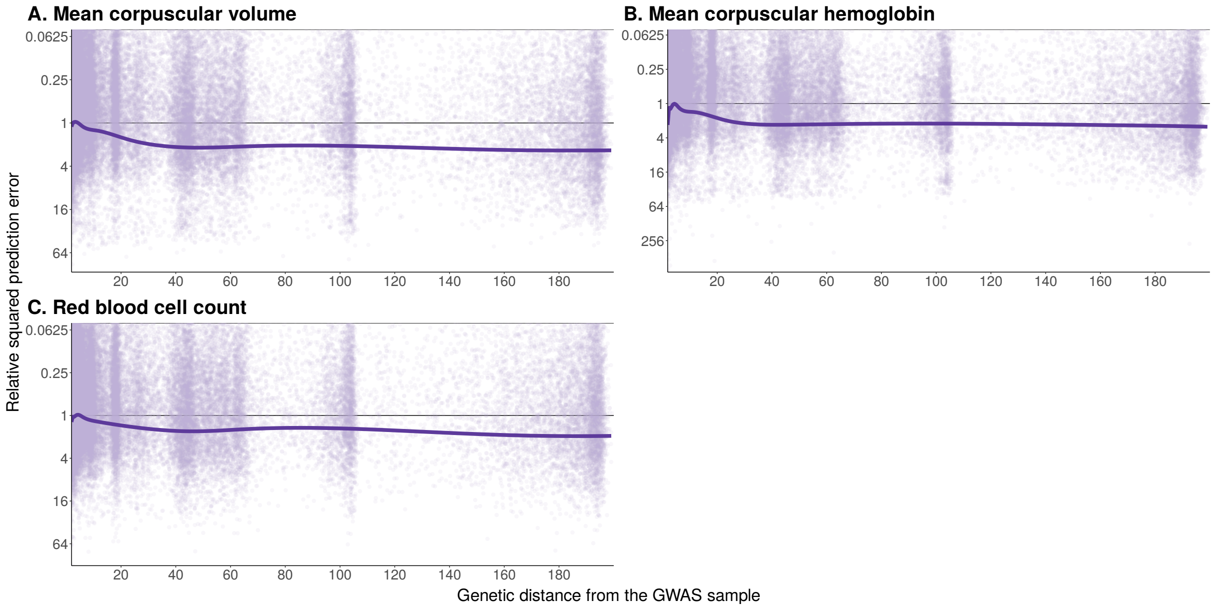

Figure S8: Full range of individual level prediction error for red blood cell-related traits. This figure presents the same analysis as **Fig. 2** in the main text, but shows the full range of prediction error for red blood cell-related traits. At the individual level, we measured prediction accuracy as the squared difference between the predicted phenotype and the true phenotype value. The y-axis values show relative prediction accuracy, i.e. prediction accuracy divided by a baseline value. The baseline value is the mean prediction accuracy in 25 bins with median genetic distances that are closest to the mean genetic distance for GWAS individuals. **Table S1** details these baseline values for each trait. Curves show cubic spline fits, with 8 knots placed based on the density of data points. See **Figs. S2-S5** for plots showing the zoomed-in version of individual-level prediction accuracy.

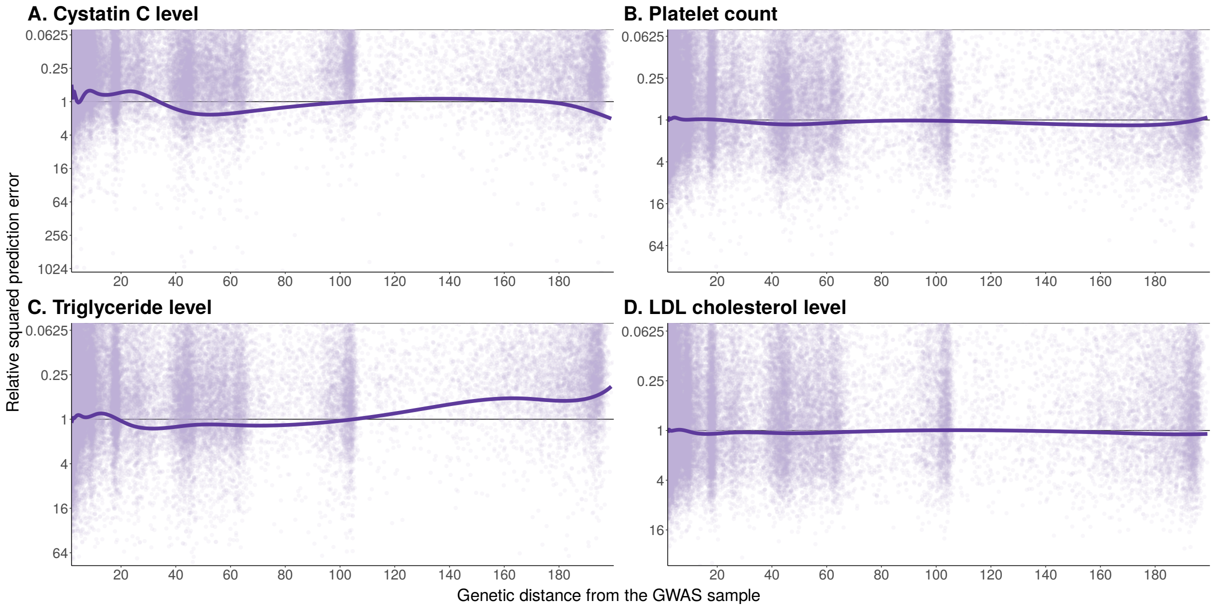

Figure S9: Full range of individual level prediction error for other biomarkers. This figure presents the same analysis as **Fig. 2** in the main text, but shows the full range of prediction error for other biomarkers. At the individual level, we measured prediction accuracy as the squared difference between the predicted phenotype and the true phenotype value. The y-axis values show relative prediction accuracy, i.e. prediction accuracy divided by a baseline value. The baseline value is the mean prediction accuracy in 25 bins with median genetic distances that are closest to the mean genetic distance for GWAS individuals. **Table S1** details these baseline values for each trait. Curves show cubic spline fits, with 8 knots placed based on the density of data points. See **Figs. S2-S5** for plots showing the zoomed-in version of individual-level prediction accuracy.

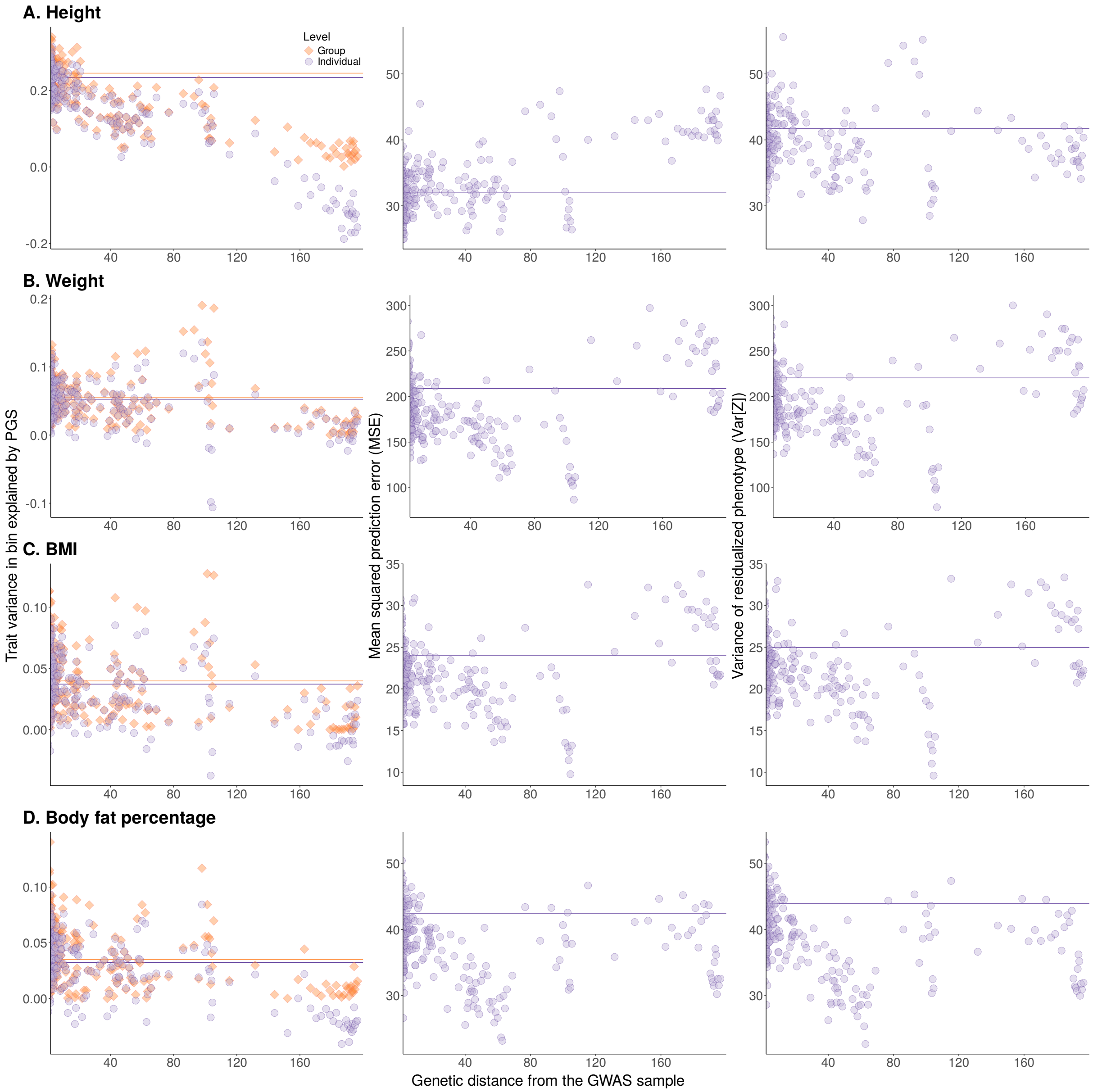

Figure S10: Divergence of group- and individual-level prediction accuracy for anthropometric measurements. The squared prediction error is with respect to the PGS as a predictor of a phenotypic value residualized for covariates ($Z$). Its mean (MSE) and the variation of residualized phenotype ($Var[Z]$) in each bin are shown in the middle and right column, respecitvely. In the left column, we show measures of the variance explained in bin of about 260 individuals, binned by genetic distance. "Group" level refers to the unstandardized partial $R^2$ between PGS and phenotype. "Individual" level refers to $1 - \frac{MSE}{Var[Z]}$. Horizontal lines show mean values across the 25 bins with a genetic distance closest to the mean genetic distance of the GWAS sample individuals (reference bins). The values of the 25 reference bins in all panels can be found in **Table S1**.

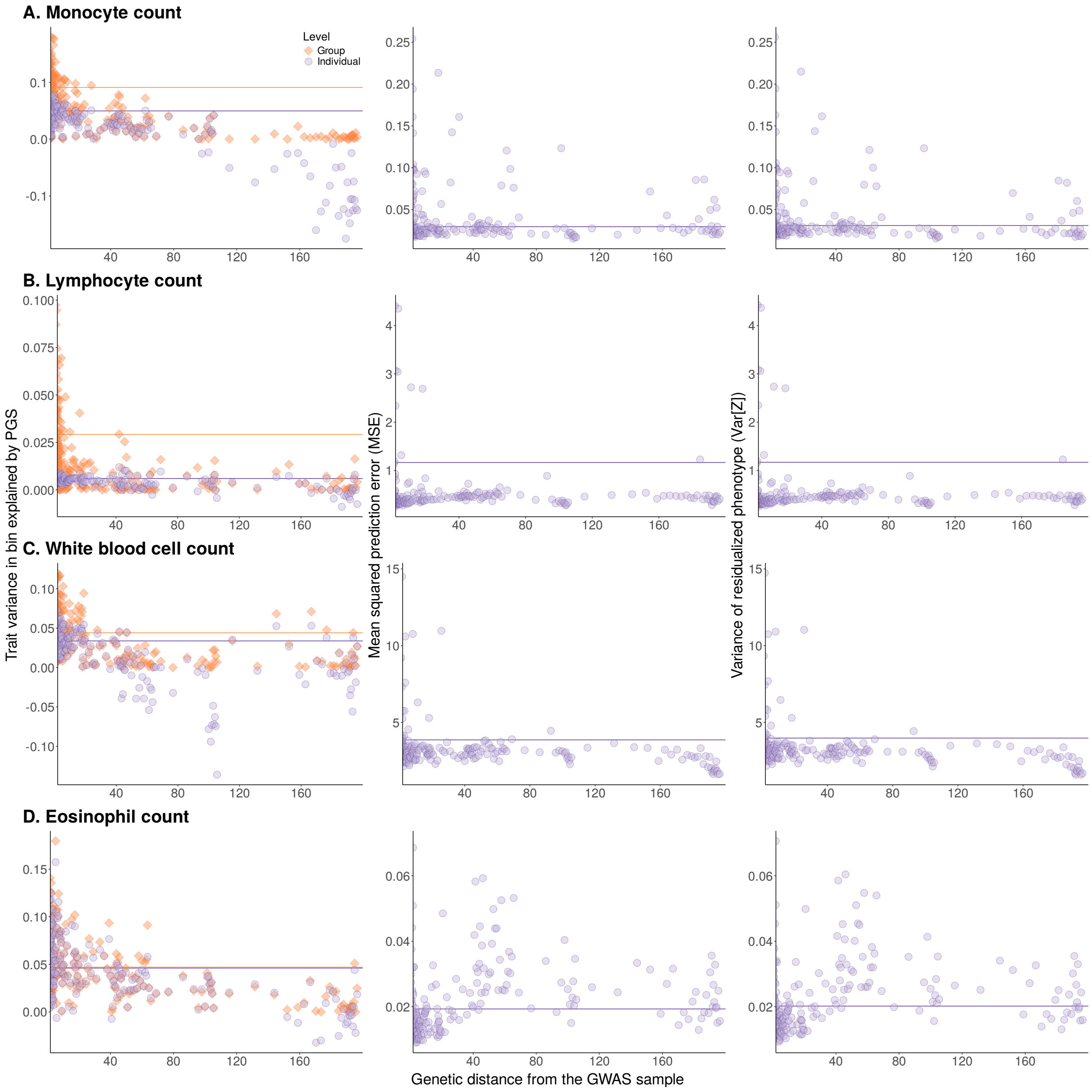

Figure S11: Divergence of group- and individual-level prediction accuracy for white blood cell-related traits. The squared prediction error is with respect to the PGS as a predictor of a phenotypic value residualized for covariates ($Z$). Its mean (MSE) and the variation of residualized phenotype ($Var[Z]$) in each bin are shown in the middle and right column, respecitvely. In the left column, we show measures of the variance explained in bin of about 260 individuals, binned by genetic distance. "Group" level refers to the unstandardized partial $R^2$ between PGS and phenotype. "Individual" level refers to $1 - \frac{MSE}{Var[Z]}$. Horizontal lines show mean values across the 25 bins with a genetic distance closest to the mean genetic distance of the GWAS sample individuals (reference bins). The values of the 25 reference bins in all panels can be found in **Table S1**.

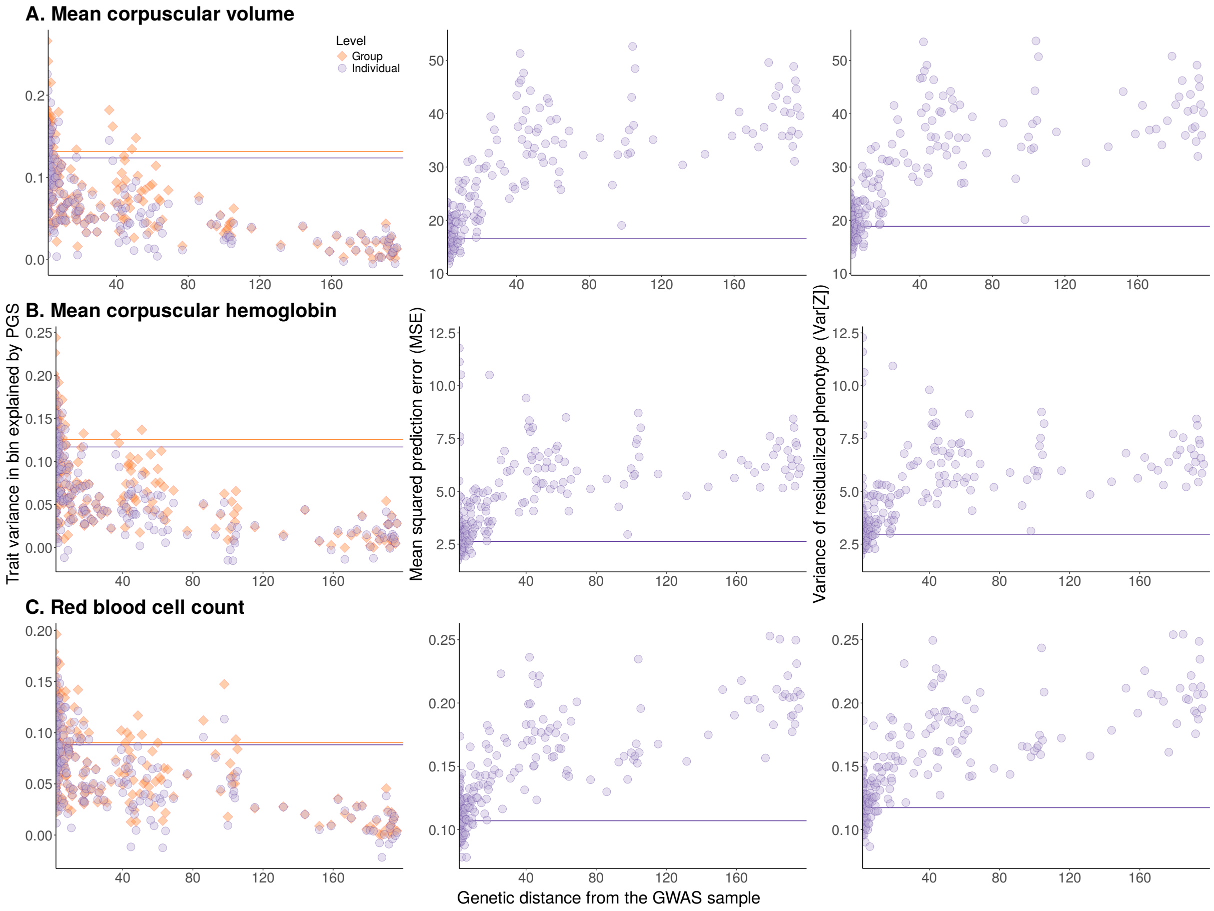

Figure S12: Divergence of group- and individual-level prediction accuracy for red blood cell-related traits. The squared prediction error is with respect to the PGS as a predictor of a phenotypic value residualized for covariates ($Z$). Its mean (MSE) and the variation of residualized phenotype ($Var[Z]$) in each bin are shown in the middle and right column, respecitvely. In the left column, we show measures of the variance explained in bin of about 260 individuals, binned by genetic distance. "Group" level refers to the unstandardized partial $R^2$ between PGS and phenotype. "Individual" level refers to $1 - \frac{MSE}{Var[Z]}$. Horizontal lines show mean values across the 25 bins with a genetic distance closest to the mean genetic distance of the GWAS sample individuals (reference bins). The values of the 25 reference bins in all panels can be found in **Table S1**.

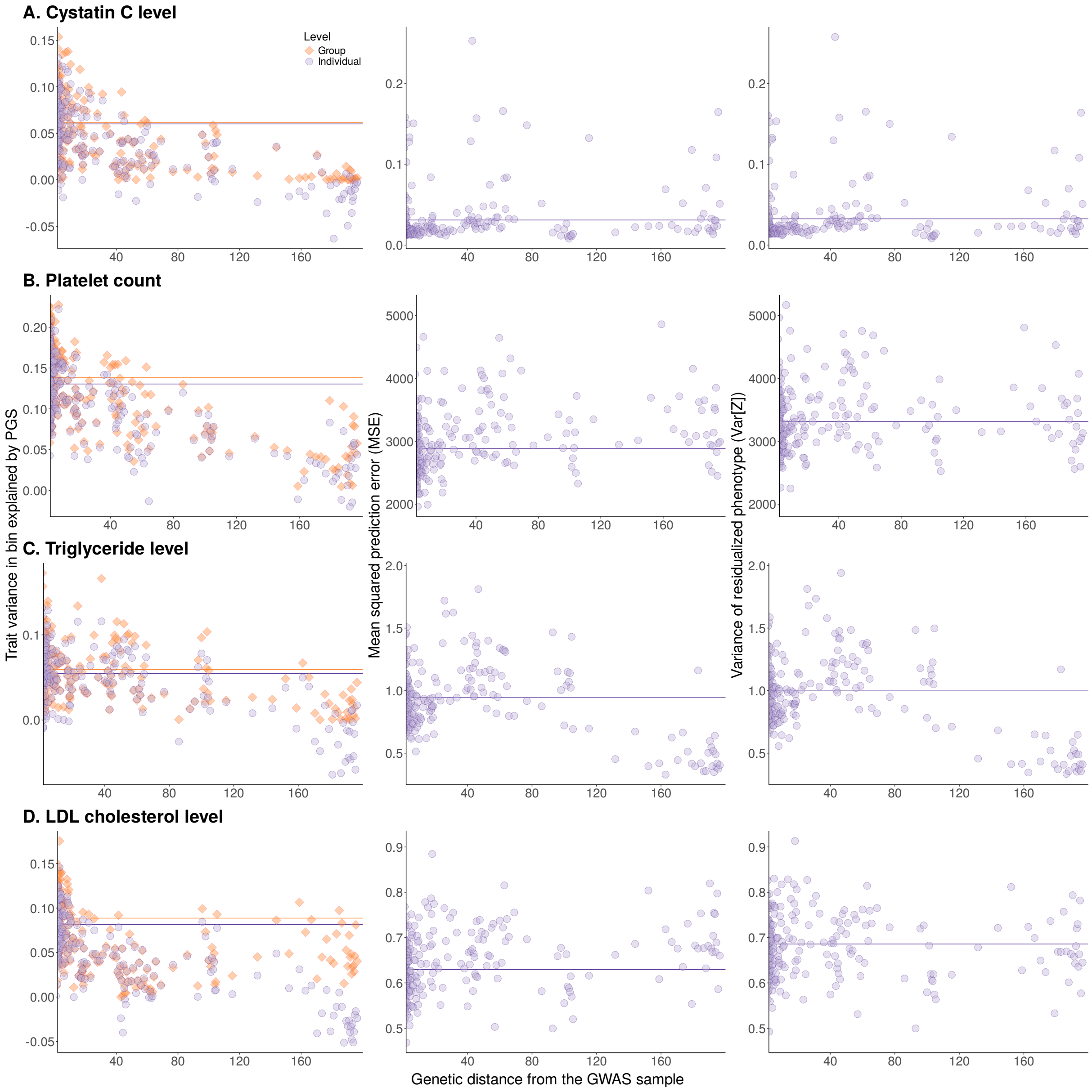

Figure S13: Divergence of group- and individual-level prediction accuracy for other biomarkers. The squared prediction error is with respect to the PGS as a predictor of a phenotypic value residualized for covariates ($Z$). Its mean (MSE) and the variation of residualized phenotype ($Var[Z]$) in each bin are shown in the middle and right column, respecitvely. In the left column, we show measures of the variance explained in bin of about 260 individuals, binned by genetic distance. "Group" level refers to the unstandardized partial $R^2$ between PGS and phenotype. "Individual" level refers to $1 - \frac{MSE}{Var[Z]}$. Horizontal lines show mean values across the 25 bins with a genetic distance closest to the mean genetic distance of the GWAS sample individuals (reference bins). The values of the 25 reference bins in all panels can be found in **Table S1**.

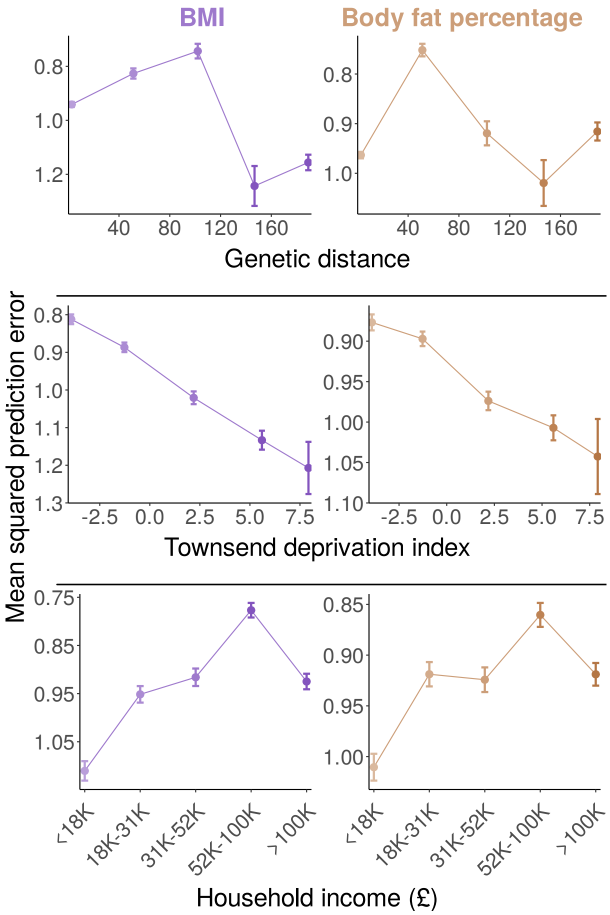

Figure S14: Mean trends in individual-level prediction accuracy by different measures for anthropometric measurements. This figure presents the same analysis as **Fig. 3A** in the main text, but shows other anthropometric traits: body mass index and body fat percentage. Data points confer to mean (±SE) squared prediction errors of individuals in the prediction sample, binned into 5 equidistant strata. The x-axis shows the median measure value for each stratum. "Household income" refers to average yearly total household income before tax.

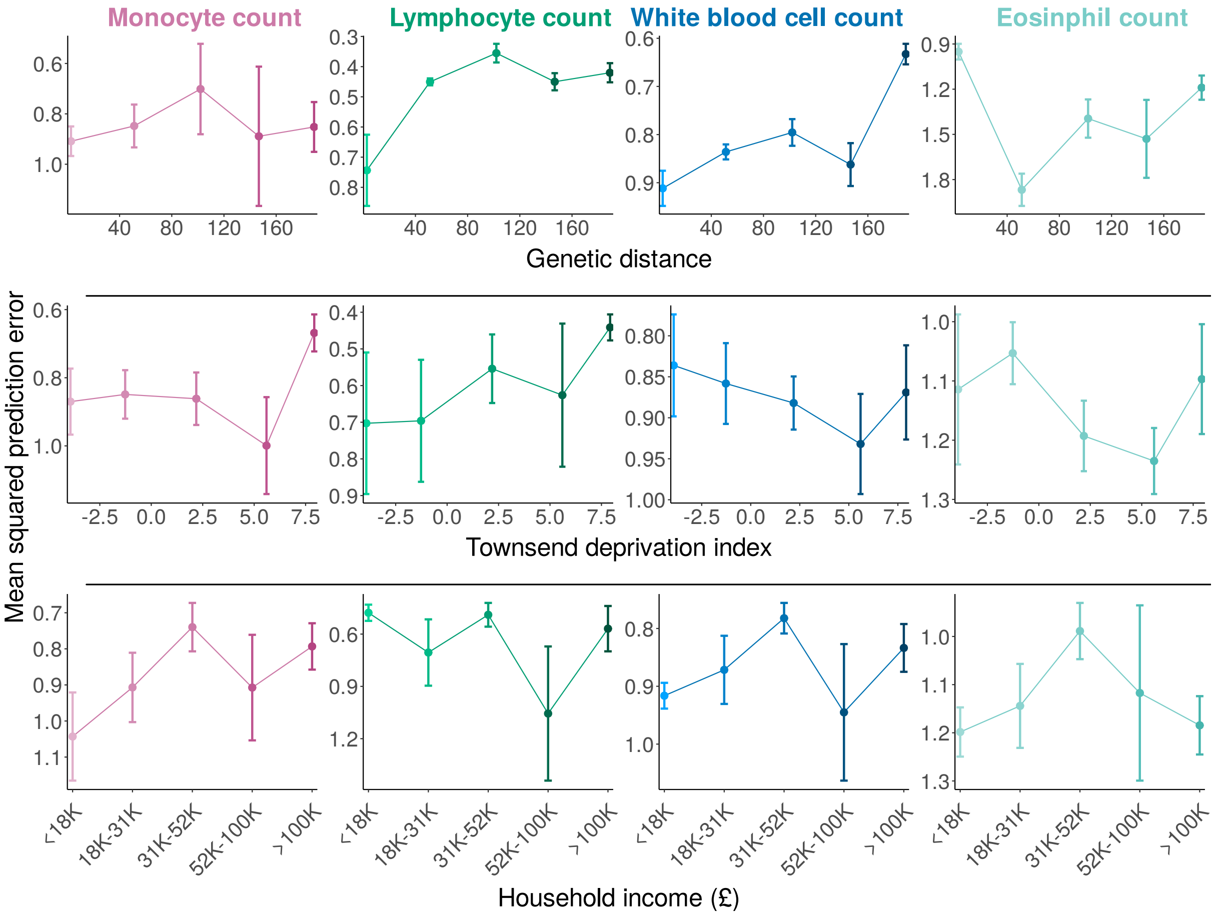

Figure S15: Mean trends in individual-level prediction accuracy by different measures for white blood cell-related traits. This figure presents the same analysis as **Fig. 3A** in the main text, but shows the mean trends for white blood cell-related traits. Data points confer to mean (±SE) squared prediction errors of individuals in the prediction sample, binned into 5 equidistant strata. The x-axis shows the median measure value for each stratum. "Household income" refers to average yearly total household income before tax.

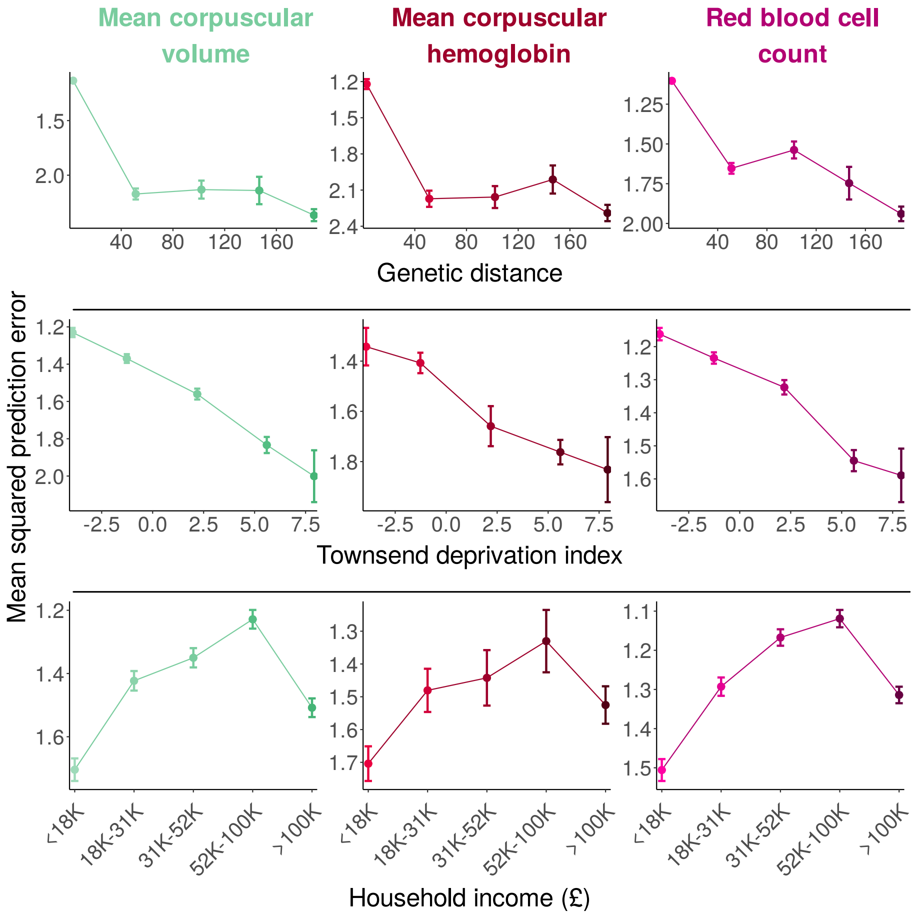

Figure S16: Mean trends in individual-level prediction accuracy by different measures for red blood cell-related traits. This figure presents the same analysis as **Fig. 3A** in the main text, but shows different traits: mean corpuscular volume, mean corpuscular hemoglobin, and red blood cell count. Data points confer to mean (±SE) squared prediction errors of individuals in the prediction sample, binned into 5 equidistant strata. The x-axis shows the median measure value for each stratum. "Household income" refers to average yearly total household income before tax.

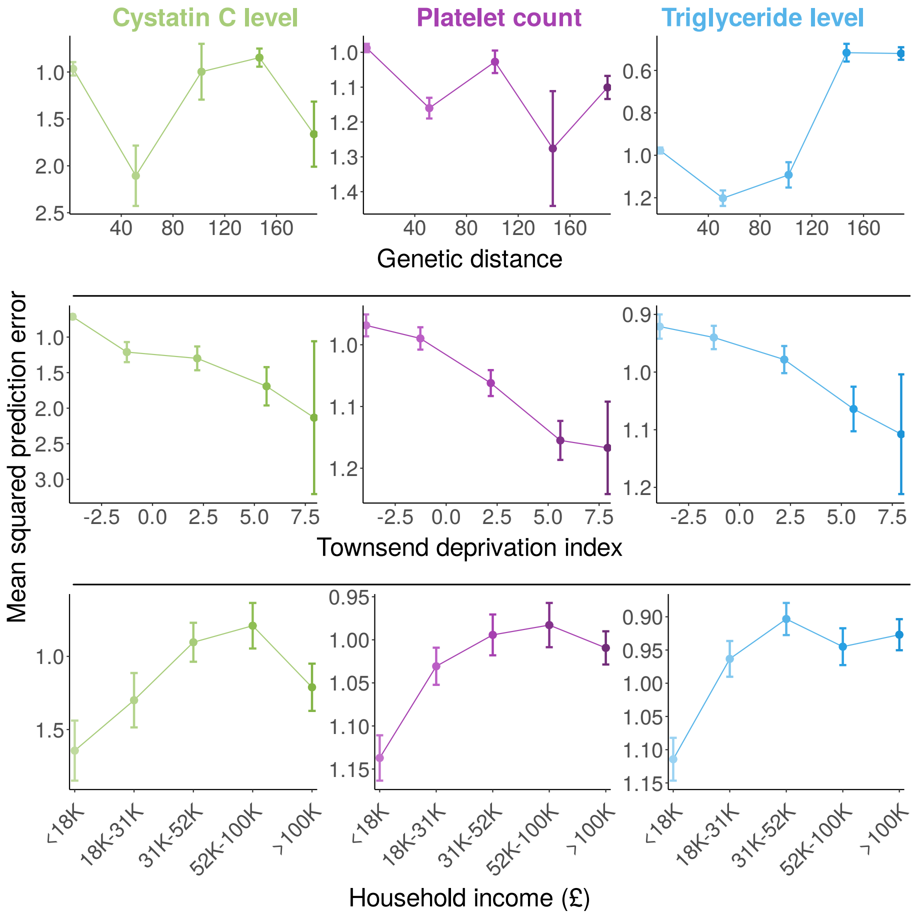

Figure S17: Mean trends in individual-level prediction accuracy by different measures for other biomarkers. This figure presents the same analysis as **Fig. 3A** in the main text, but shows different traits: cystatin C level, platelet count, and triglyceride level. Data points confer to mean (±SE) squared prediction errors of individuals in the prediction sample, binned into 5 equidistant strata. The x-axis shows the median measure value for each stratum. "Household income" refers to average yearly total household income before tax.

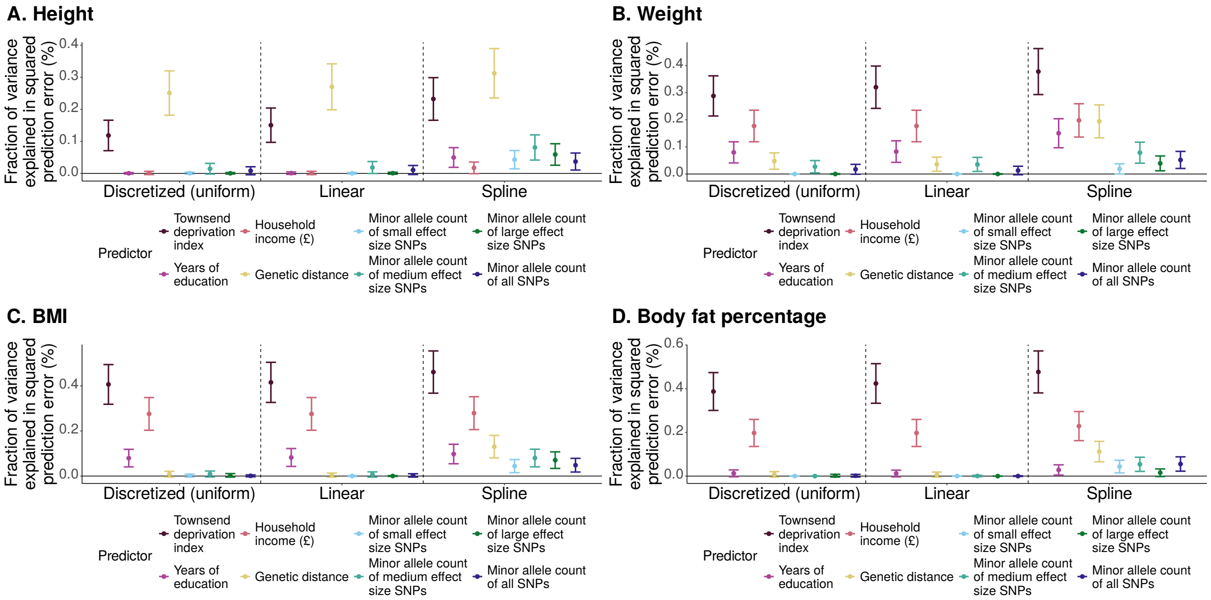

Figure S18: Individual-level prediction error explained by different measures for anthropometric measurements. This figure presents the same analysis as **Fig. 3B** in the main text, but shows more measures and methods for fitting the measures, and focusing on anthropometric traits. “Minor allele” refers to the minor allele with respect to the GWAS sample. For each measure, we independently fit three different models. “Discretized (uniform)” refers to a discretized predictor, using one predicted value per each of the 5 bins where all 5 bins had identical widths. “Linear” refers to a linear predictor, fit using Ordinary Least Squares (OLS). “Spline” refers to a cubic spline, for which 16 knots were placed based on the density of data points, such that there was an equal number of data points between each pair of consecutive knots.

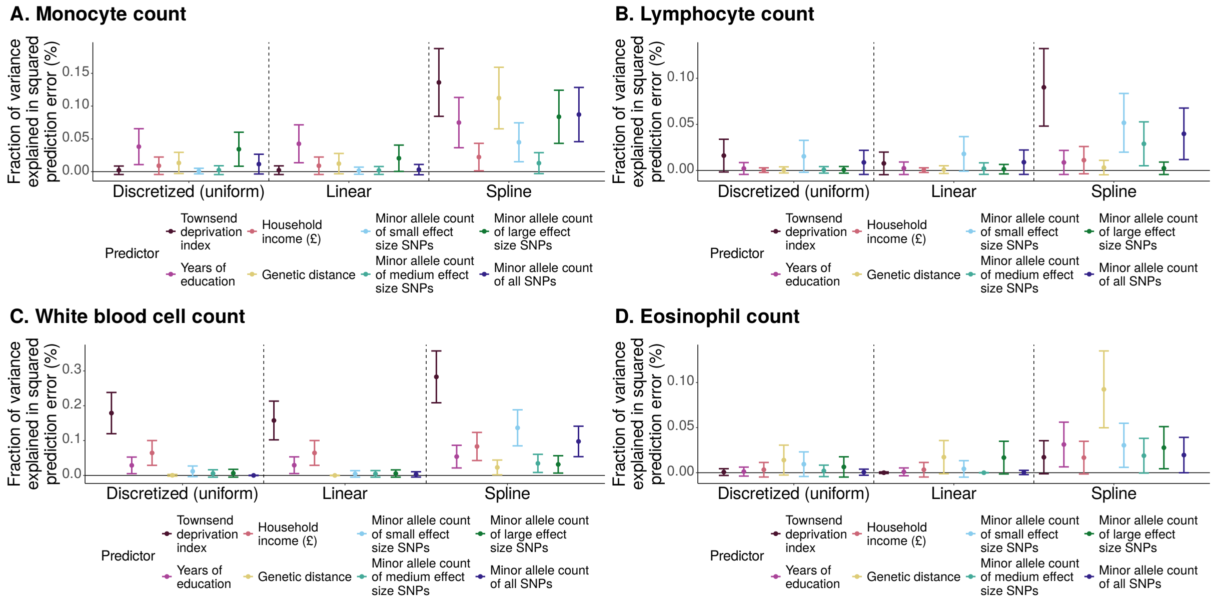

Figure S19: Individual-level prediction error explained by different measures for white blood cell-related traits. This figure presents the same analysis as **Fig. 3B** in the main text, but shows more measures and methods for fitting the measures. "Minor allele" refers to the minor allele with respect to the GWAS sample. For each measure, we independently fit three different models. "Discretized (uniform)" refers to a discretized predictor, using one predicted value per each of the 5 bins where all 5 bins had identical widths. "Linear" refers to a linear predictor, fit using Ordinary Least Squares (OLS). "Spline" refers to a cubic spline, for which 16 knots were placed based on the density of data points, such that there was an equal number of data points between each pair of consecutive knots.

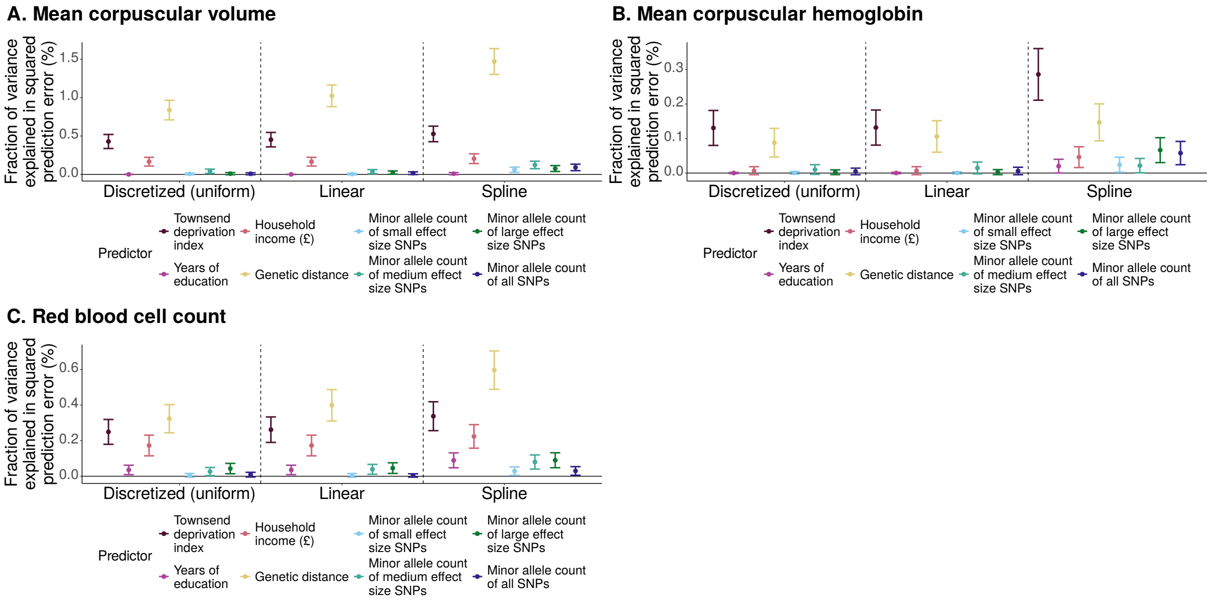

Figure S20: Individual-level prediction error explained by different measures for red blood cell-related traits. This figure presents the same analysis as **Fig. 3B** in the main text, but shows more measures and methods for fitting the measures. "Minor allele" refers to the minor allele with respect to the GWAS sample. For each measure, we independently fit three different models. "Discretized (uniform)" refers to a discretized predictor, using one predicted value per each of the 5 bins where all 5 bins had identical widths. "Linear" refers to a linear predictor, fit using Ordinary Least Squares (OLS). "Spline" refers to a cubic spline, for which 16 knots were placed based on the density of data points, such that there was an equal number of data points between each pair of consecutive knots.

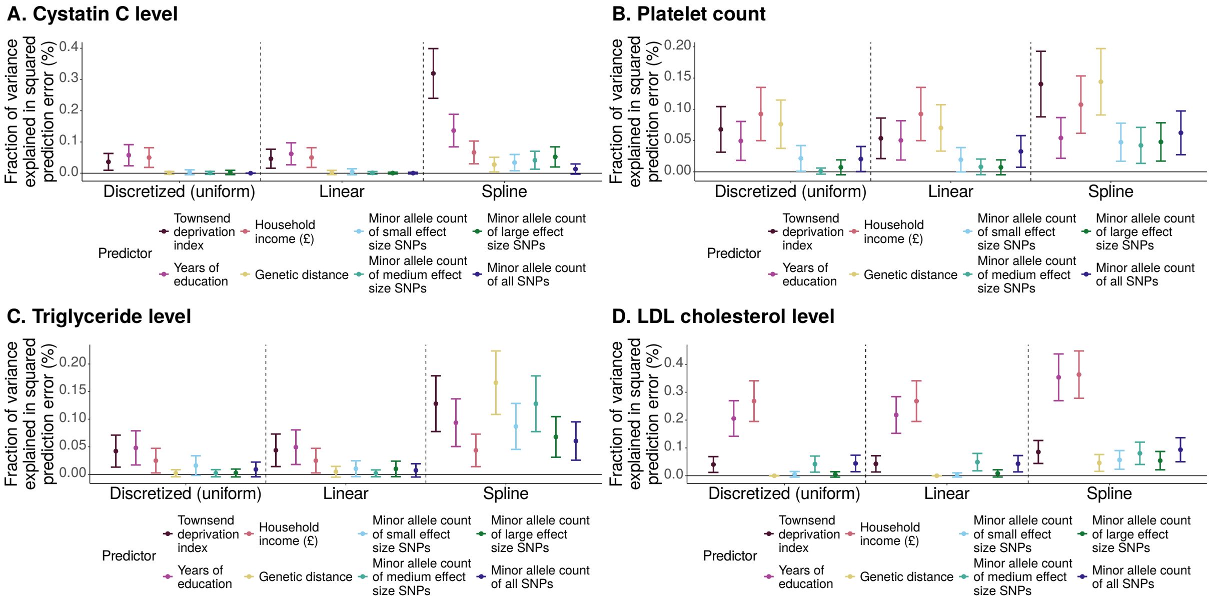

Figure S21: Individual-level prediction error explained by different measures for other biomarkers. This figure presents the same analysis as **Fig. 3B** in the main text, but shows more measures and methods for fitting the measures. "Minor allele" refers to the minor allele with respect to the GWAS sample. For each measure, we independently fit three different models. "Discretized (uniform)" refers to a discretized predictor, using one predicted value per each of the 5 bins where all 5 bins had identical widths. "Linear" refers to a linear predictor, fit using Ordinary Least Squares (OLS). "Spline" refers to a cubic spline, for which 16 knots were placed based on the density of data points, such that there was an equal number of data points between each pair of consecutive knots.

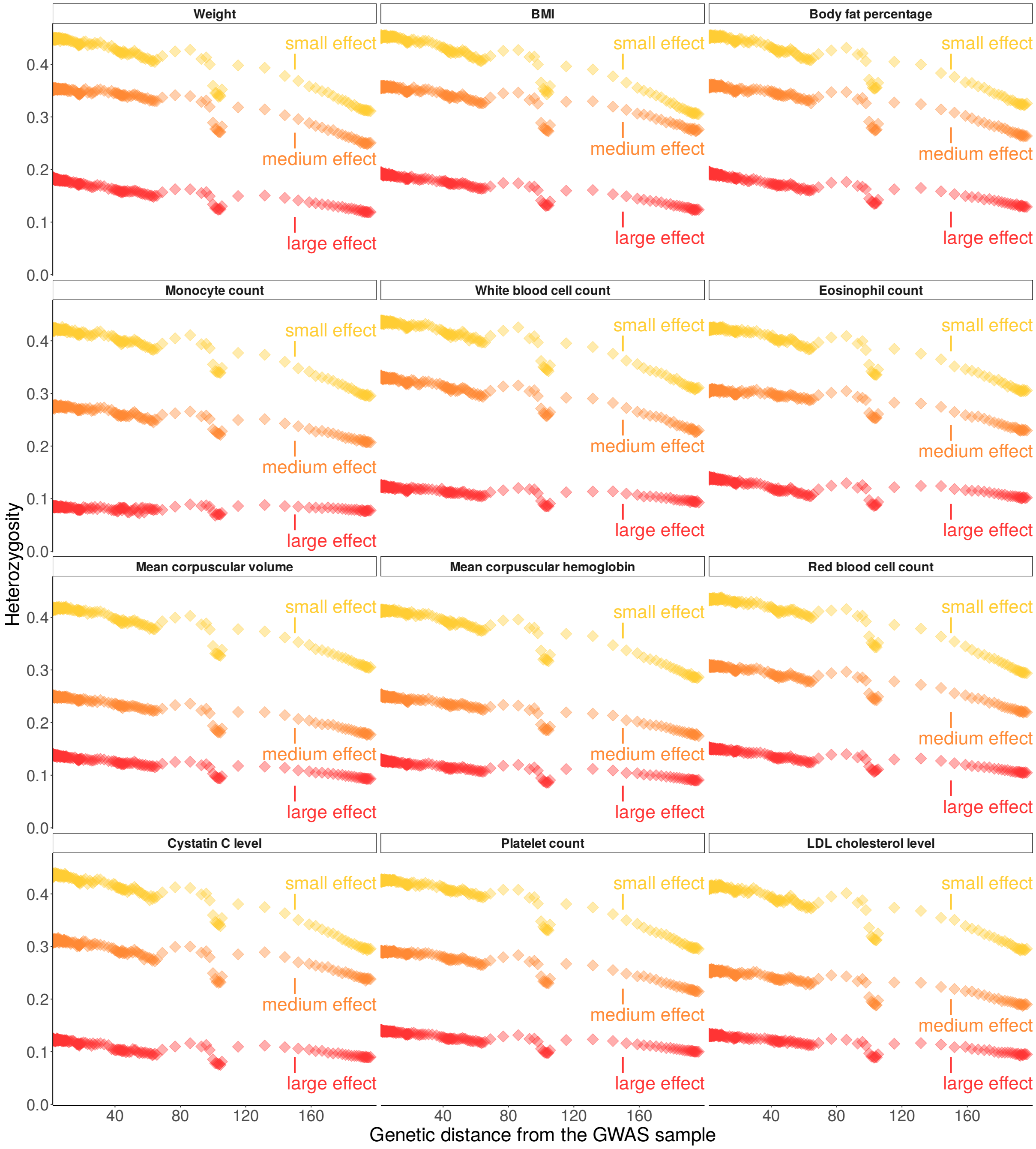

Figure S22: Mean heterozygosity of SNPs, stratified by effect size. This figure presents the same analysis as **Fig. 4B** in the main text, but for showing data for the 12 quantitative traits that we analyzed and are not included there. For each trait, SNPs are stratified into three equal-sized strata (small, medium, and large) based on squared effect sizes (**Fig. S23**). Each data point is the mean heterozygosity of a stratum in a bin of genetic distance.

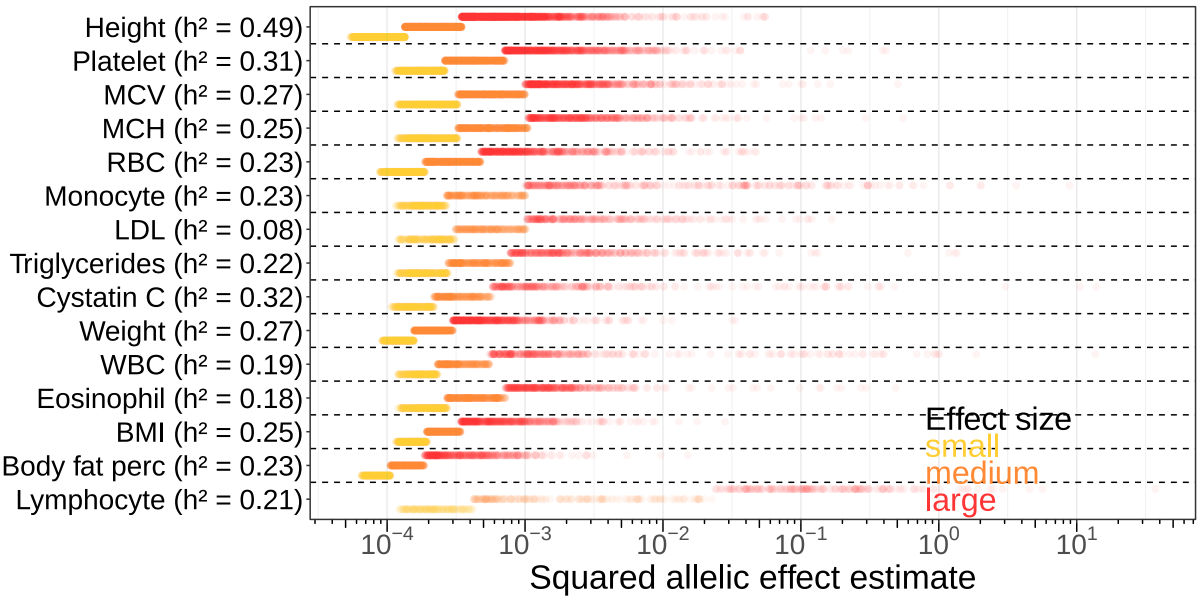

Figure S23: Squared allelic effect estimate. For each trait, the index SNPs of the respective PGS are stratified into three equal-sized strata (small, medium, and large) based on squared effect sizes. The x-axis represents the squared effect sizes in units of trait variance in the GWAS set. SNP heritabilities ($h^2$) are taken from the Neale Lab's UKB analysis[26]. Each data point represents a SNP. MCV: mean corpuscular volume. MCH: mean corpuscular hemoglobin. RBC: red blood cell count. Body fat perc: body fat percentage. WBC: white blood cell count. LDL: LDL cholesterol level.

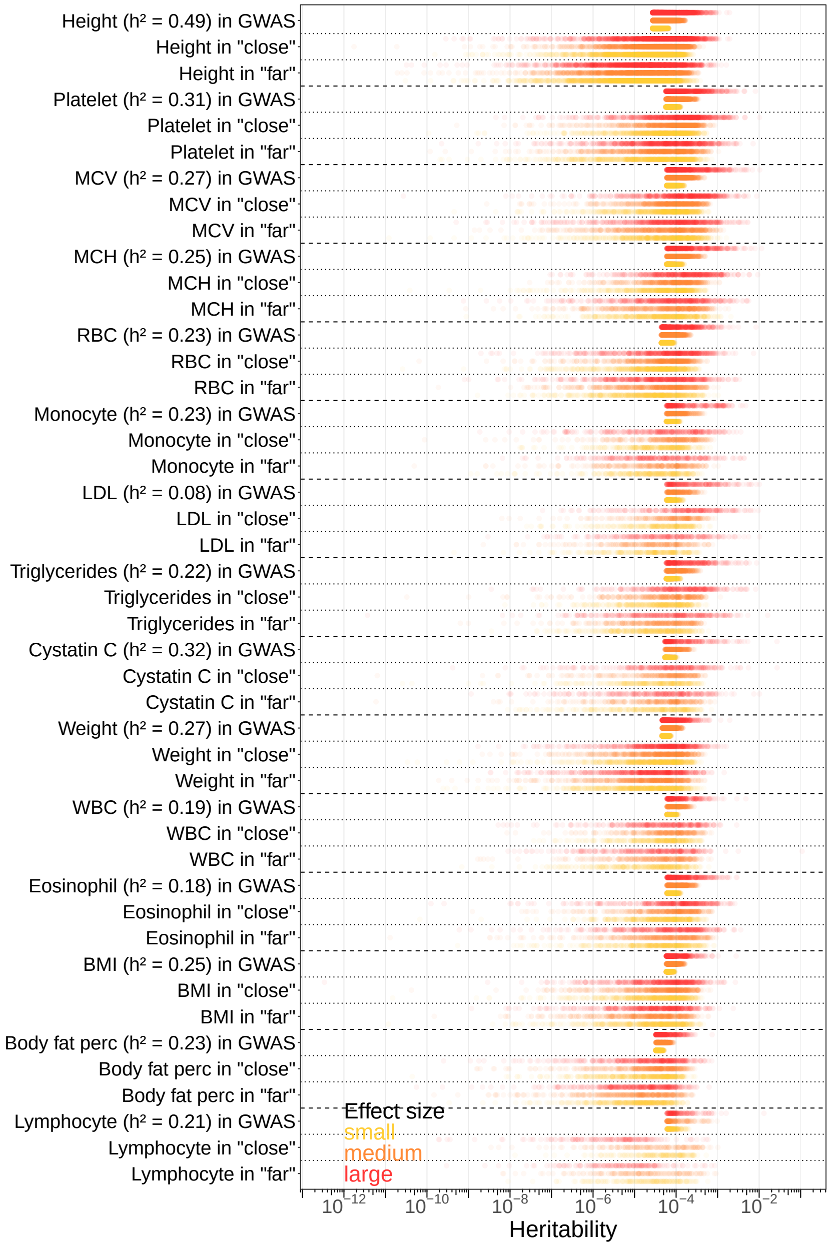

Figure S24: Heritability explained by index SNPs. For each trait, index SNPs of the respective PGS are stratified by their squared effect sizes. Data points confer to index SNPs. In each of the GWAS sample, the "close" subset of the prediction sample (genetic distance $\leq 10$ in the prediction set, with 38,992 individuals) and the "far" subsample of the prediction sample (genetic distance $> 10$, with 30,508 individuals), we estimate the allelic effect of each SNP (in units of trait standard deviations) and its heterozygosity. The product of the two is the estimated heritability. SNP heritability estimates ($h^2$) on the y-axis are taken from the Neale Lab's UKB analysis[26].

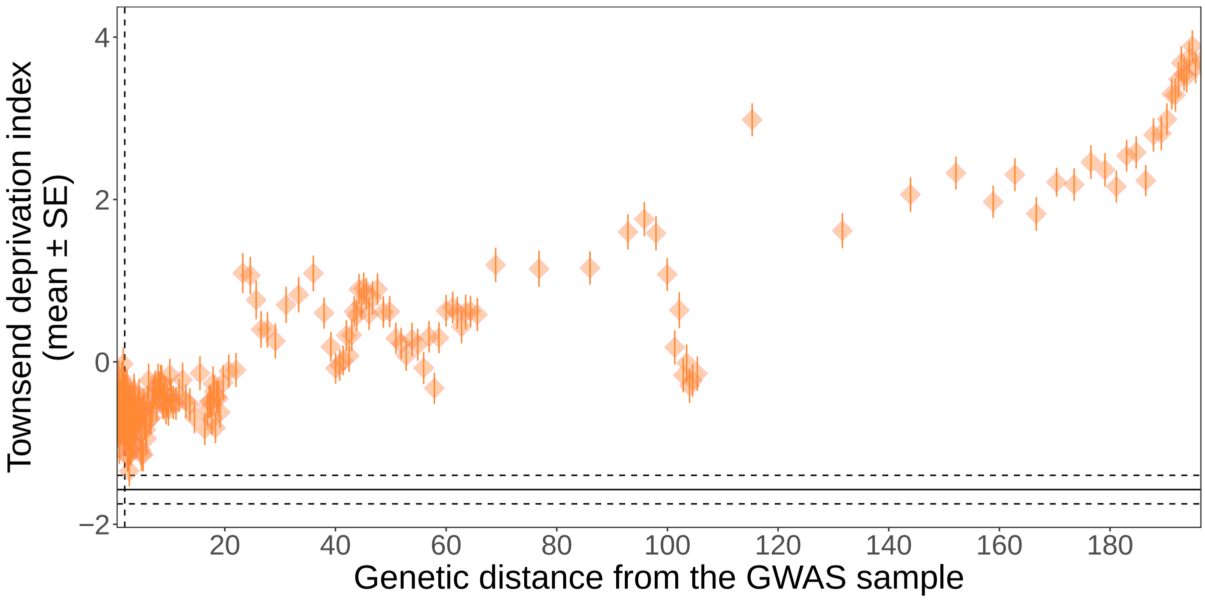

Figure S25: Mean Townsend Deprivation Index ± SE in each bin by genetic distance. In this figure, the orange dot and line represent the mean Townsend Deprivation Index ± SE in each bin in the prediction set. The horizontal solid line represents the mean Townsend Deprivation Index in the GWAS set. The horizontal dashed lines are calculated as mean in the GWAS sample $\pm \frac{SD_{\text{GWAS}}}{\sqrt{\bar{n_{\text{pred}}}}}$, where $\bar{n_{pred}}$ represents the mean sample size across all bins in the prediction set (278). The vertical dashed line represents the 97.5$^{\text{th}}$ percentile of genetic distance among GWAS sample individuals.

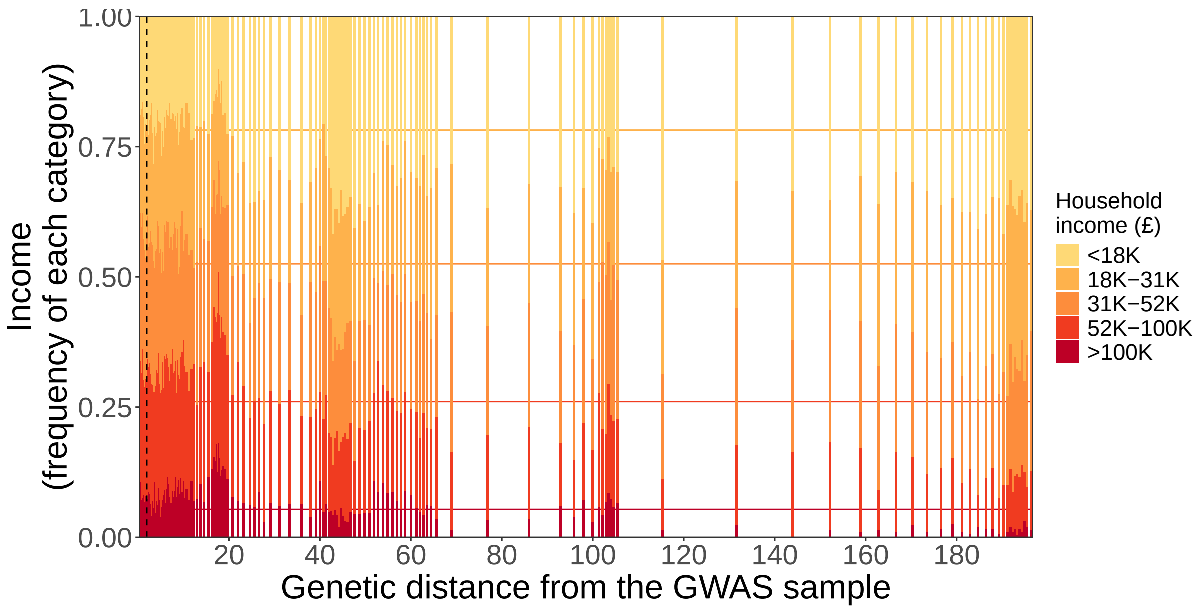

Figure S26: Average total household income before tax in each bin by genetic distance. In this figure, each vertical colored bar corresponds to a bin in the prediction set, with the cumulative frequency of income category color-coded. The horizontal colored lines represent the cumulative frequencies in the GWAS set. The vertical dashed line represents the 97.5$^{\text{th}}$ percentile of genetic distance among GWAS sample individuals.

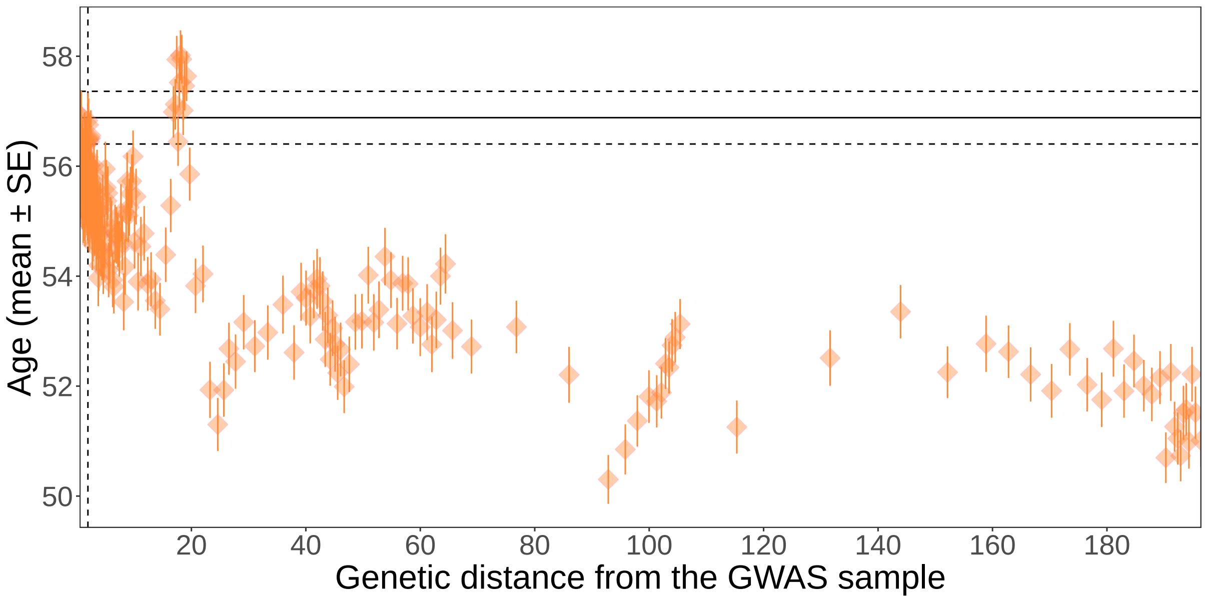

Figure S27: Mean age ± SE in each bin by genetic distance. In this figure, the orange dot and line represent the mean age ± SE in each bin in the prediction set. The horizontal solid line represents the mean age in the GWAS set. The horizontal dashed lines are calculated as mean in the GWAS sample $\pm \frac{SD_{GWAS}}{\sqrt{\bar{n_{pred}}}}$, where $\bar{n_{pred}}$ represents the mean sample size across all bins in the prediction set (278). The vertical dashed line represents the 97.5$^{\text{th}}$ percentile of genetic distance among GWAS sample individuals.

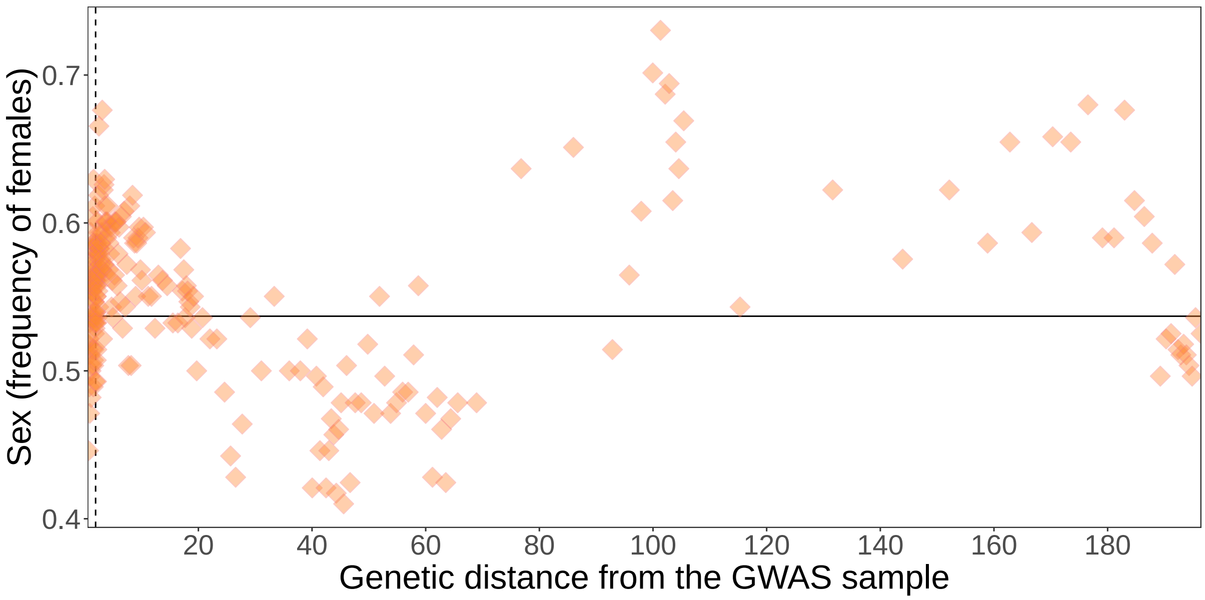

Figure S28: Sex composition across bins of genetic distance. Orange dots represent the mean frequency of females in each genetic distance bin in the prediction sample. The horizontal solid black line represents the mean frequency of females in the GWAS set. The vertical dashed line represents the 97.5$^{th}$ percentile of genetic distance among GWAS sample individuals.

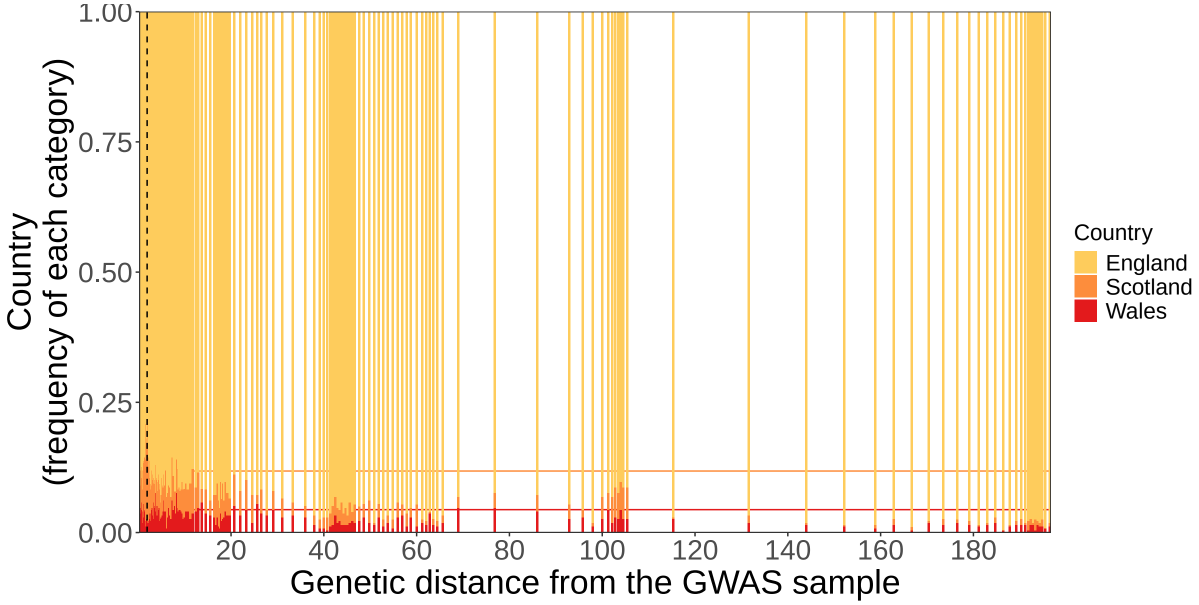

Figure S29: Participants' country of residence across bins of genetic distance.. Each vertical colored bar corresponds to a bin in the prediction set, with the cumulative frequency of country of residence color-coded. The horizontal colored lines represent the cumulative frequencies in the GWAS set. The vertical dashed line represents the $97.5^{\text{th}}$ percentile of genetic distance among GWAS sample individuals.

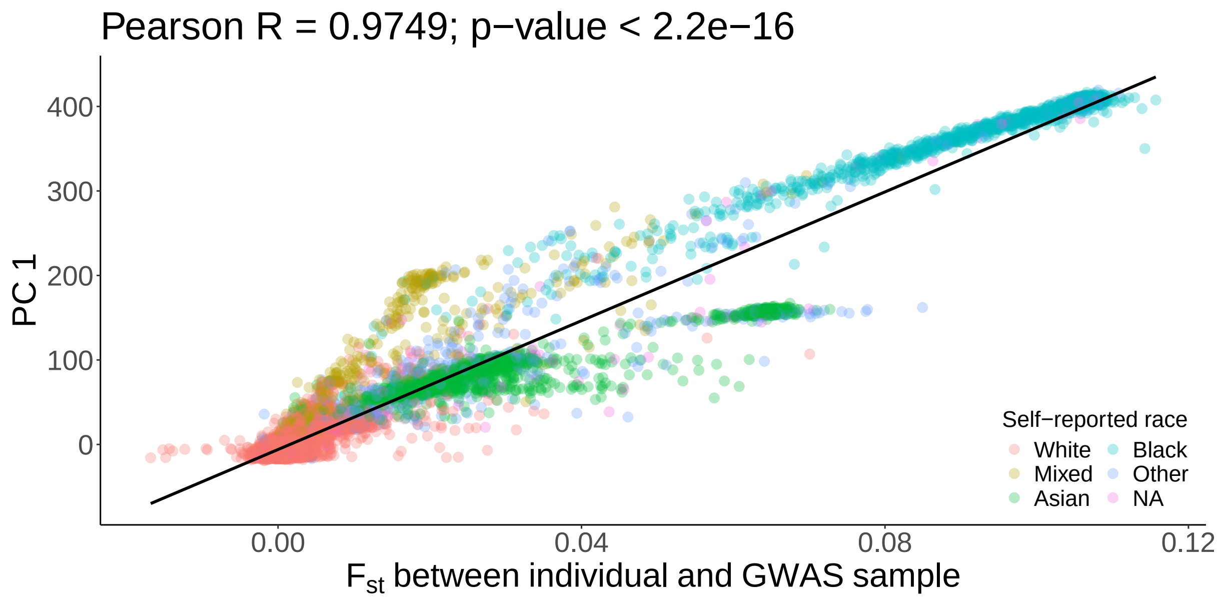

Figure S30: Correlation between $F_{st}$ and PC 1. This figure presents the correlation between $F_{st}$ and PC 1. The data points are colored by self-reported race.

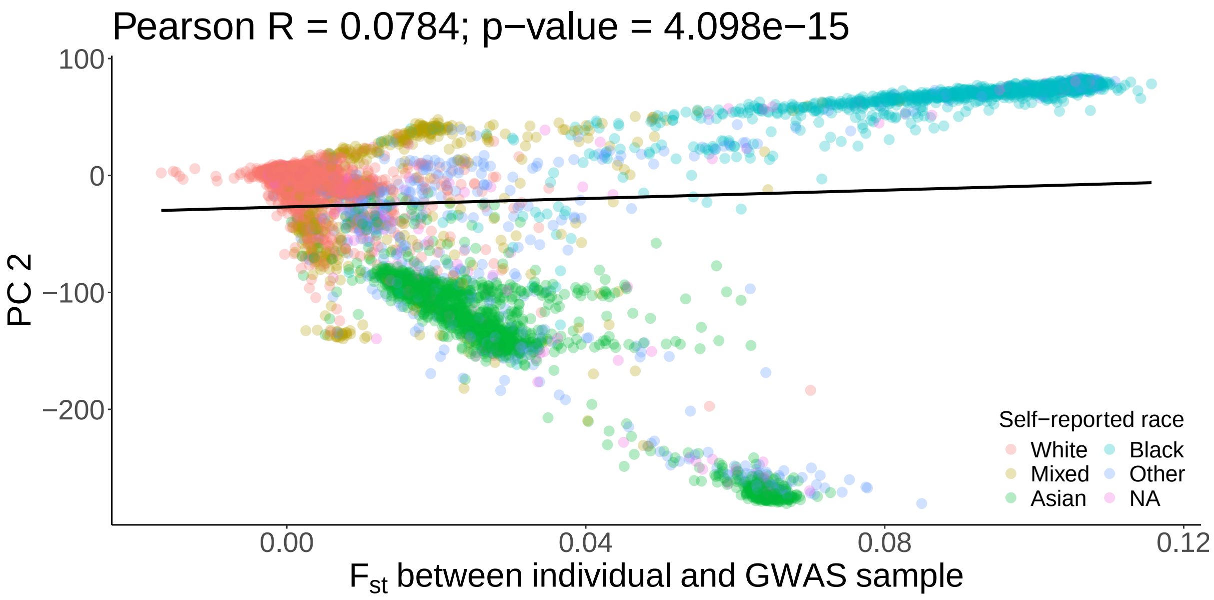

Figure S31: Correlation between $F_{st}$ and PC 2. This figure presents the correlation between $F_{st}$ and PC 2. The data points are colored by self-reported race.

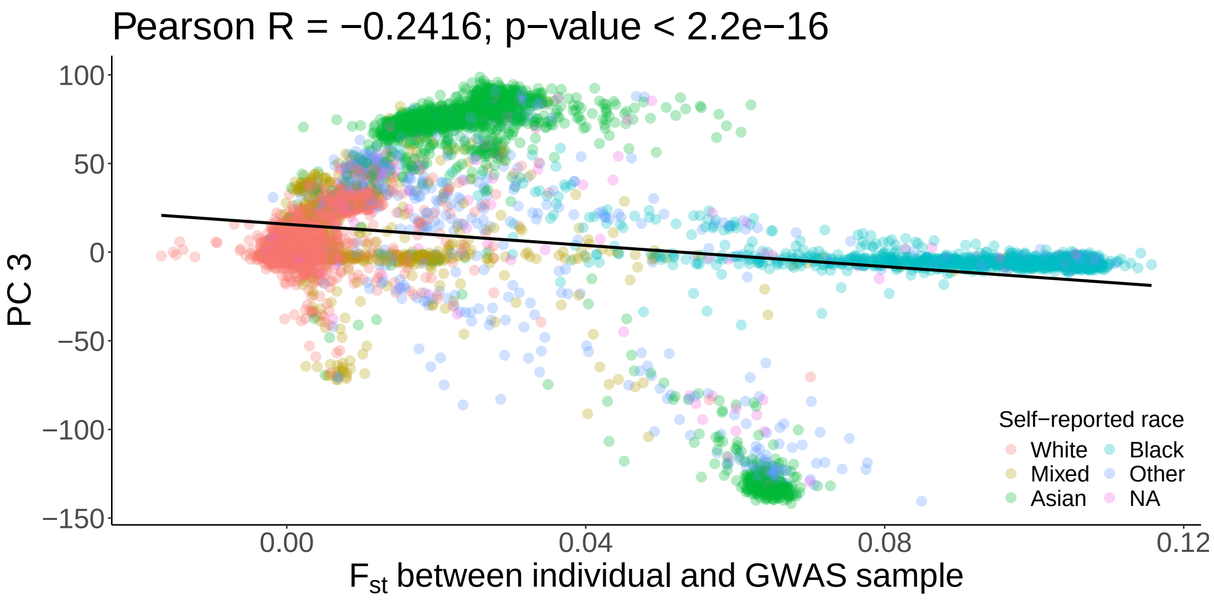

Figure S32: Correlation between $F_{st}$ and PC 3. This figure presents the correlation between $F_{st}$ and PC 3. The data points are colored by self-reported race.

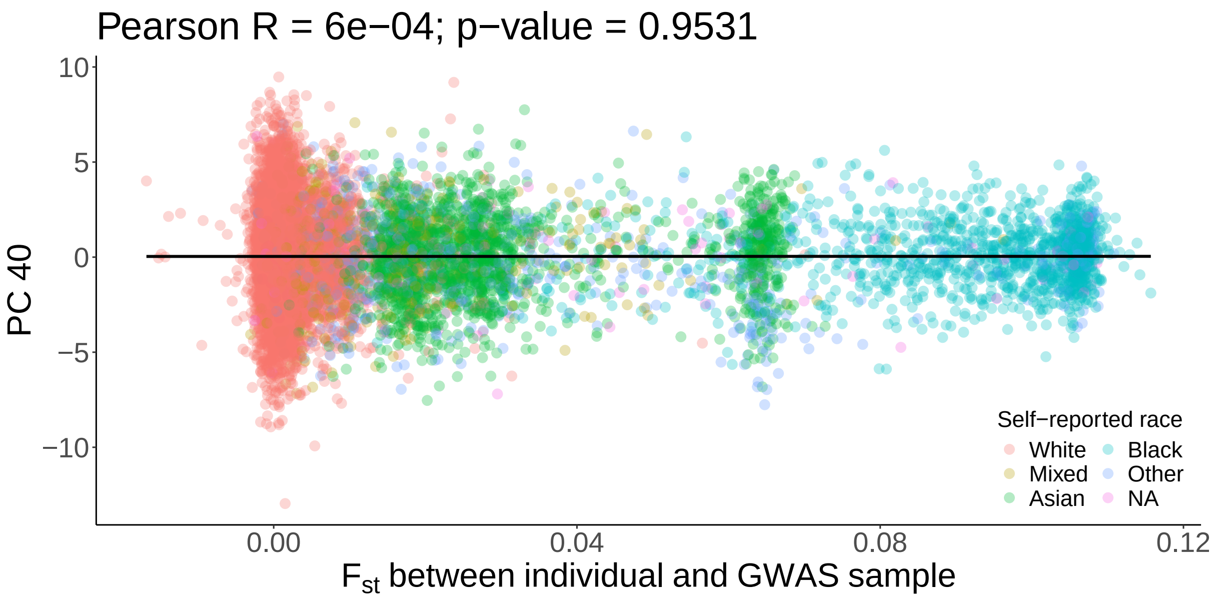

Figure S33: Correlation between $F_{st}$ and PC 40. This figure presents the correlation between $F_{st}$ and PC 40. The data points are colored by self-reported race.

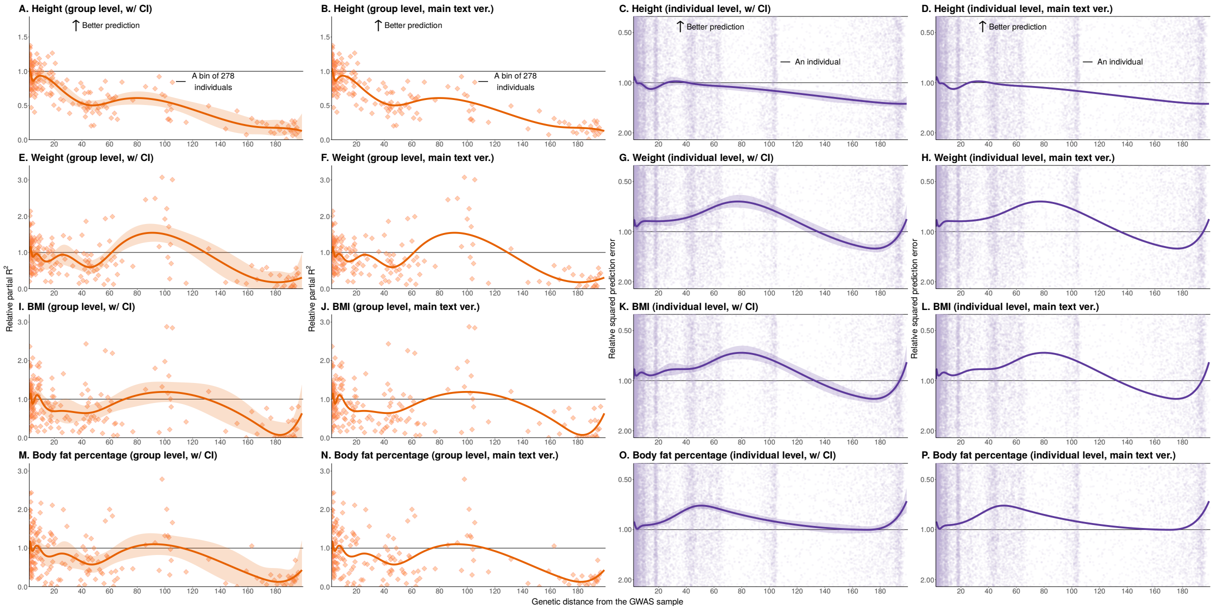

Figure S34: Trends of portability vary across traits and measures, with confidence intervals, for anthropometric measurements. This figure presents the same analysis as **Fig. 2** and **Fig. S2**, but the first and third columns (**A, C, E, G, I, K, M, O**) are plots with confidence intervals. The second and fourth columns (**B, D, F, H, J, L, N, P**) contain plots taken from the main text or supplementary text for comparison. At the group level (left two columns), we measured prediction accuracy with the squared partial correlation between the PGS and the trait value in 250 bins of 278 individuals each. At the individual level (right two columns), we measured prediction accuracy as the squared difference between the predicted phenotype and the true phenotype value. At both the group and individual levels, y-axis values show relative prediction accuracy, i.e. prediction accuracy divided by a baseline value. The baseline value is the mean prediction accuracy in 25 bins with median genetic distances that are closest to the mean genetic distance for GWAS individuals. **Table S1** details these baseline values for each trait. Curves show cubic spline fits, with 8 knots placed based on the density of data points.

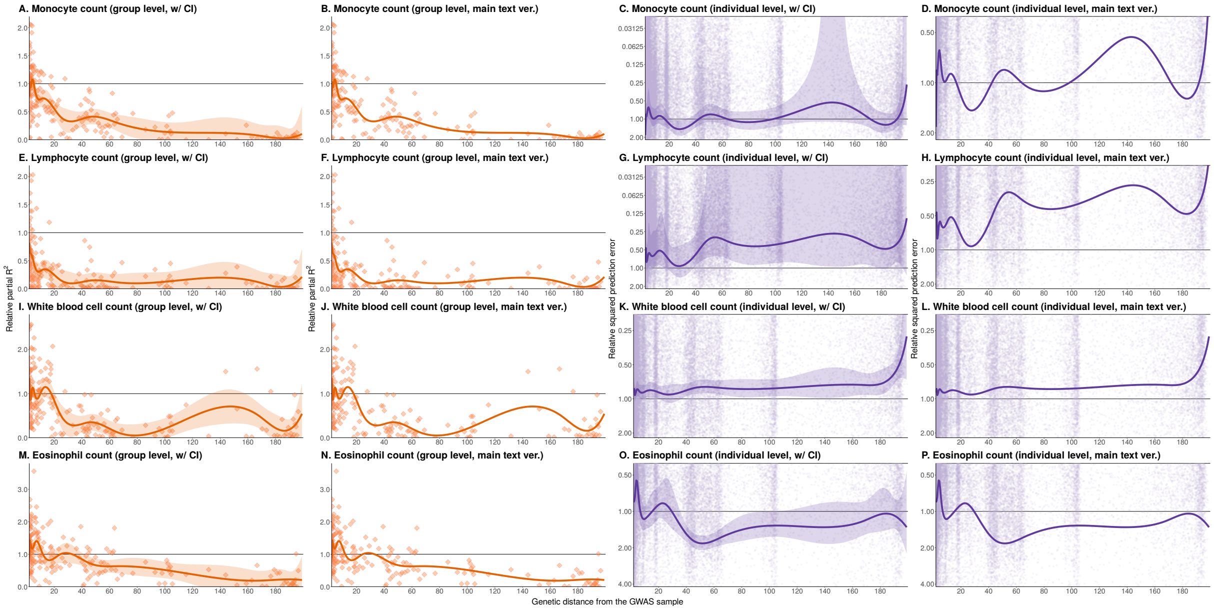

Figure S35: Trends of portability vary across traits and measures, with confidence intervals, for white blood-cell related traits. This figure presents the same analysis as **Fig. 2** and **Fig. S3**, but the first and third columns (**A, C, E, G, I, K, M, O**) are plots with confidence intervals. The second and fourth columns (**B, D, F, H, J, L, N, P**) contain plots taken from the main text or supplementary text for comparison. At the group level (left two columns), we measured prediction accuracy with the squared partial correlation between the PGS and the trait value in 250 bins of 278 individuals each. At the individual level (right two columns), we measured prediction accuracy as the squared difference between the predicted phenotype and the true phenotype value. At both the group and individual levels, y-axis values show relative prediction accuracy, i.e. prediction accuracy divided by a baseline value. The baseline value is the mean prediction accuracy in 25 bins with median genetic distances that are closest to the mean genetic distance for GWAS individuals. **Table S1** details these baseline values for each trait. Curves show cubic spline fits, with 8 knots placed based on the density of data points. For **C, G, K**, when plotting the CI on a logarithmic scale, we replaced the values extremely close to zero with $10^{-6}$.

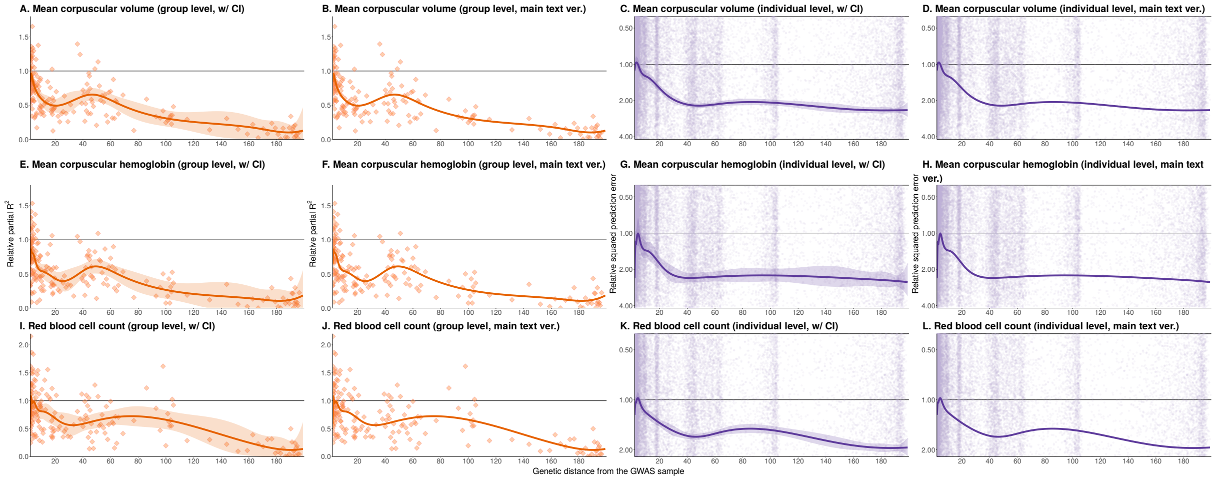

Figure S36: Trends of portability vary across traits and measures, with confidence intervals, for red blood cell-related traits. This figure presents the same analysis as **Fig. 2** and **Fig. S4**, but the first and third columns (**A, C, E, G, I, K**) are plots with confidence intervals. The second and fourth columns (**B, D, F, H, J, L**) contain plots taken from the main text or supplementary text for comparison. At the group level (left two columns), we measured prediction accuracy with the squared partial correlation between the PGS and the trait value in 250 bins of 278 individuals each. At the individual level (right two columns), we measured prediction accuracy as the squared difference between the predicted phenotype and the true phenotype value. At both the group and individual levels, y-axis values show relative prediction accuracy, i.e. prediction accuracy divided by a baseline value. The baseline value is the mean prediction accuracy in 25 bins with median genetic distances that are closest to the mean genetic distance for GWAS individuals. **Table S1** details these baseline values for each trait. Curves show cubic spline fits, with 8 knots placed based on the density of data points.

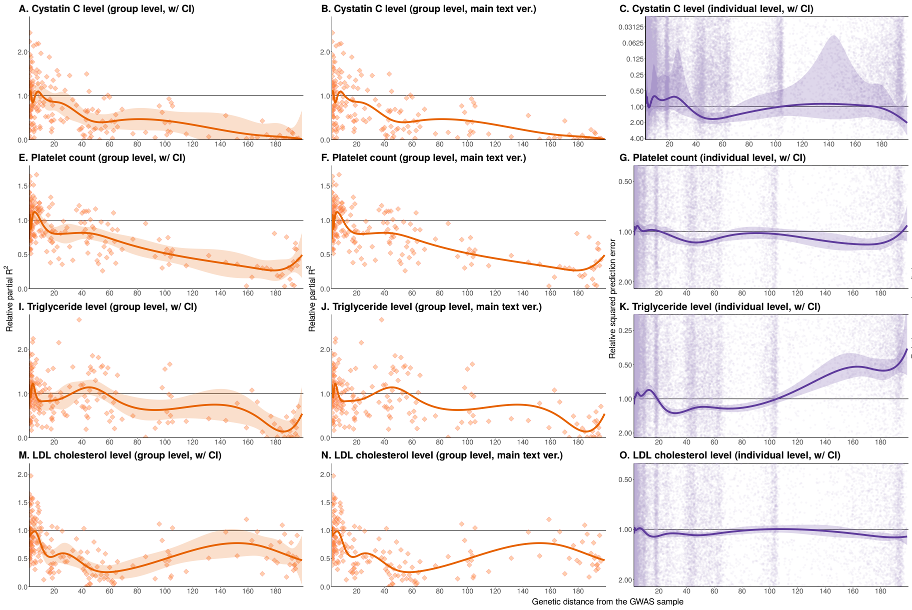

Figure S37: Trends of portability vary across traits and measures, with confidence intervals, for other biomarkers. This figure presents the same analysis as **Fig. 2** and **Fig. S5**, but the first and third columns (**A, C, E, G, I, K, M, O**) are plots with 95% confidence intervals. The second and fourth columns (**B, D, F, H, J, L, N, P**) contain plots taken from the main text or supplementary text for comparison. At the group level (left two columns), we measured prediction accuracy with the squared partial correlation between the PGS and the trait value in 250 bins of 278 individuals each. At the individual level (right two columns), we measured prediction accuracy as the squared difference between the predicted phenotype and the true phenotype value. At both the group and individual levels, y-axis values show relative prediction accuracy, i.e. prediction accuracy divided by a baseline value. The baseline value is the mean prediction accuracy in 25 bins with median genetic distances that are closest to the mean genetic distance for GWAS individuals. **Table S1** details these baseline values for each trait. Curves show cubic spline fits, with 8 knots placed based on the density of data points.

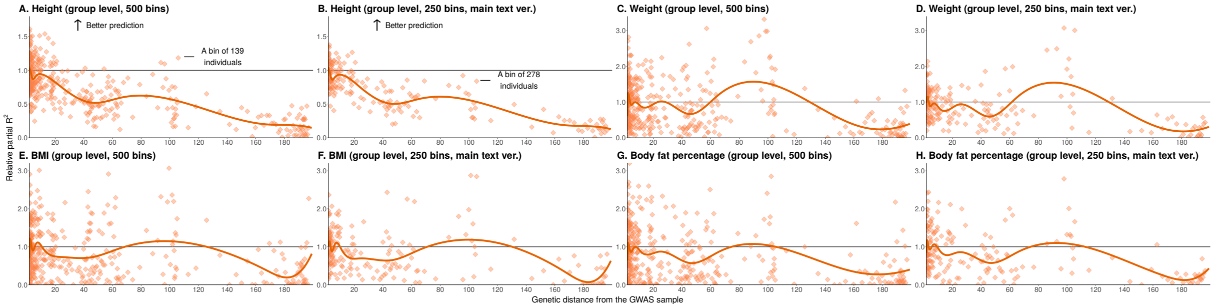

Figure S38: Group level portability, using more bins, for anthropometric measurements. This figure presents the same analysis as **Fig. 2** and **Fig. S2** at the group level, but the first and third columns (**A, C, E, G**) are plots with 500 bins of 139 each. The second and fourth columns (**B, D, F, H**) contain plots with 250 bins of 278 each, taken from the main text or supplementary text for comparison. At the group level, we measured prediction accuracy with the squared partial correlation between the PGS and the trait value. The y-axis values show relative prediction accuracy, i.e. prediction accuracy divided by a baseline value. The baseline value for the panels with 500 bins is the mean prediction accuracy in 50 bins with median genetic distances that are closest to the mean genetic distance for GWAS individuals, and is based on 25 bins with median genetic distances that are closest to the mean genetic distance for GWAS individuals for the original plots with 250 bins. **Table S1** details these baseline values for each trait. Curves show cubic spline fits, with 8 knots placed based on the density of data points.

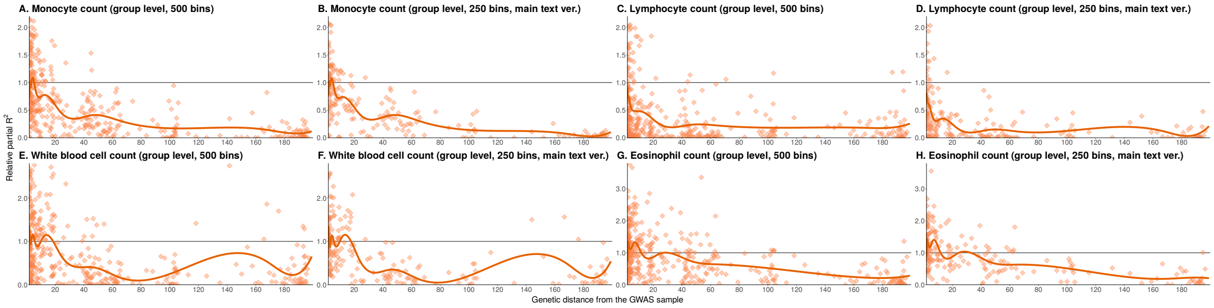

Figure S39: Group level portability, using more bins, for white blood-cell related traits. This figure presents the same analysis as **Fig. 2** and **Fig. S3** at the group level, but the first and third columns (**A, C, E, G**) are plots with 500 bins of 139 each. The second and fourth columns (**B, D, F, H**) contain plots with 250 bins of 278 each, taken from the main text or supplementary text for comparison. At the group level, we measured prediction accuracy with the squared partial correlation between the PGS and the trait value. The y-axis values show relative prediction accuracy, i.e. prediction accuracy divided by a baseline value. The baseline value for the panels with 500 bins is the mean prediction accuracy in 50 bins with median genetic distances that are closest to the mean genetic distance for GWAS individuals, and is based on 25 bins with median genetic distances that are closest to the mean genetic distance for GWAS individuals for the original plots with 250 bins. **Table S1** details these baseline values for each trait. Curves show cubic spline fits, with 8 knots placed based on the density of data points.

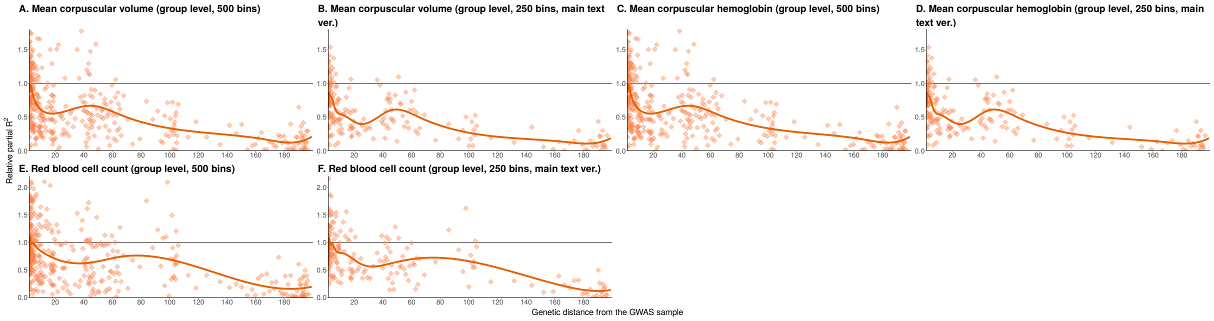

Figure S40: Group level portability, using more bins, for red blood cell-related traits. This figure presents the same analysis as **Fig. 2** and **Fig. S4** at the group level, but the first and third columns (**A, C, E**) are plots with 500 bins of 139 each. The second and fourth columns (**B, D, F**) contain plots with 250 bins of 278 each, taken from the main text or supplementary text for comparison. At the group level, we measured prediction accuracy with the squared partial correlation between the PGS and the trait value. The y-axis values show relative prediction accuracy, i.e. prediction accuracy divided by a baseline value. The baseline value for the panels with 500 bins is the mean prediction accuracy in 50 bins with median genetic distances that are closest to the mean genetic distance for GWAS individuals, and is based on 25 bins with median genetic distances that are closest to the mean genetic distance for GWAS individuals for the original plots with 250 bins. **Table S1** details these baseline values for each trait. Curves show cubic spline fits, with 8 knots placed based on the density of data points.

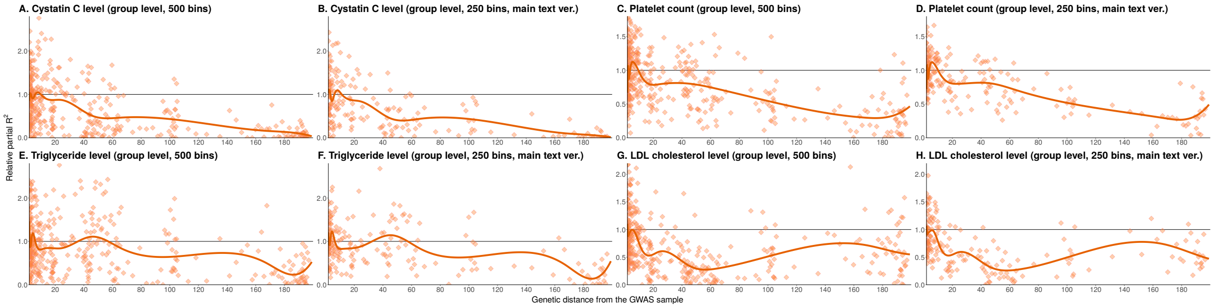

Figure S41: Group level portability, using more bins, for other biomarkers. This figure presents the same analysis as **Fig. 2** and **Fig. S5** at the group level, but the first and third columns (**A, C, E, G**) are plots with 500 bins of 139 each. The second and fourth columns (**B, D, F, H**) contain plots with 250 bins of 278 each, taken from the main text or supplementary text for comparison. At the group level, we measured prediction accuracy with the squared partial correlation between the PGS and the trait value. The y-axis values show relative prediction accuracy, i.e. prediction accuracy divided by a baseline value. The baseline value for the panels with 500 bins is the mean prediction accuracy in 50 bins with median genetic distances that are closest to the mean genetic distance for GWAS individuals, and is based on 25 bins with median genetic distances that are closest to the mean genetic distance for GWAS individuals for the original plots with 250 bins. **Table S1** details these baseline values for each trait. Curves show cubic spline fits, with 8 knots placed based on the density of data points.

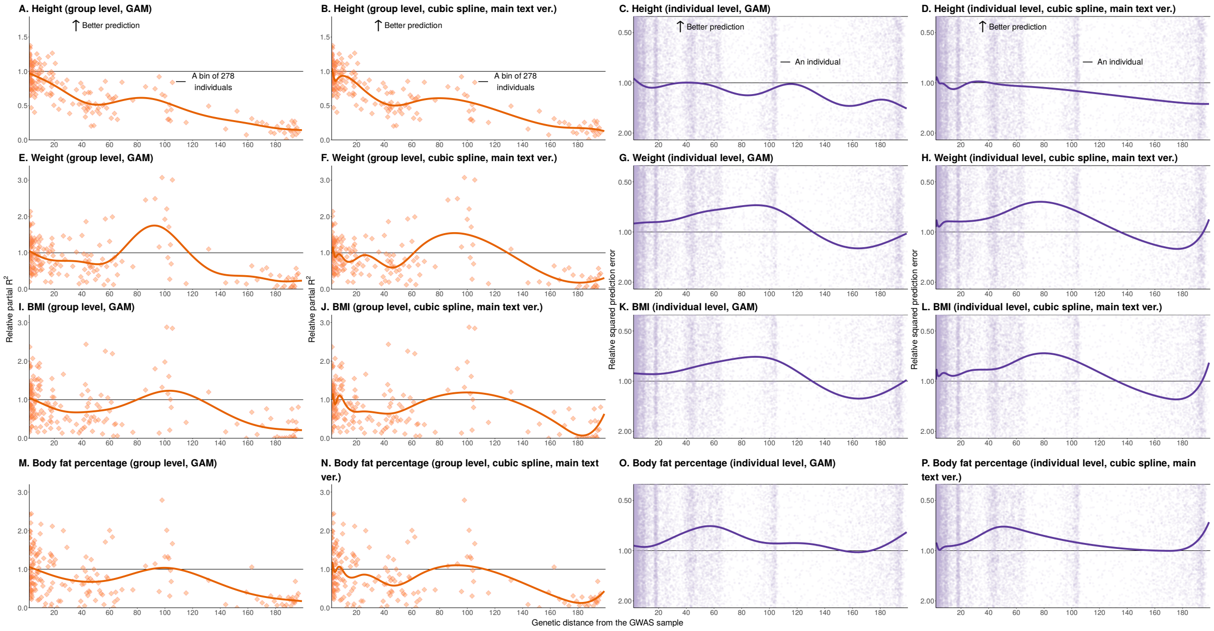

Figure S42: Trends of portability visualized with generalized additive model (GAM) fit, for anthropometric measurements. This figure presents the same analysis as **Fig. 2** and **Fig. S2**, but the first and third columns (**A, C, E, G, I, K, M, O**) are plots with GAM fits, as shown by the curves. The second and fourth columns (**B, D, F, H, J, L, N, P**) contain plots with cubic spline fits, taken from the main text or supplementary text for comparison. At the group level (left two columns), we measured prediction accuracy with the squared partial correlation between the PGS and the trait value in 250 bins of 278 individuals each. At the individual level (right two columns), we measured prediction accuracy as the squared difference between the predicted phenotype and the true phenotype value. At both the group and individual levels, y-axis values show relative prediction accuracy, i.e. prediction accuracy divided by a baseline value. The baseline value is the mean prediction accuracy in 25 bins with median genetic distances that are closest to the mean genetic distance for GWAS individuals. **Table S1** details these baseline values for each trait. For both GAM and cubic spline fits, curves show cubic spline fits, with 8 knots placed based on the density of data points.

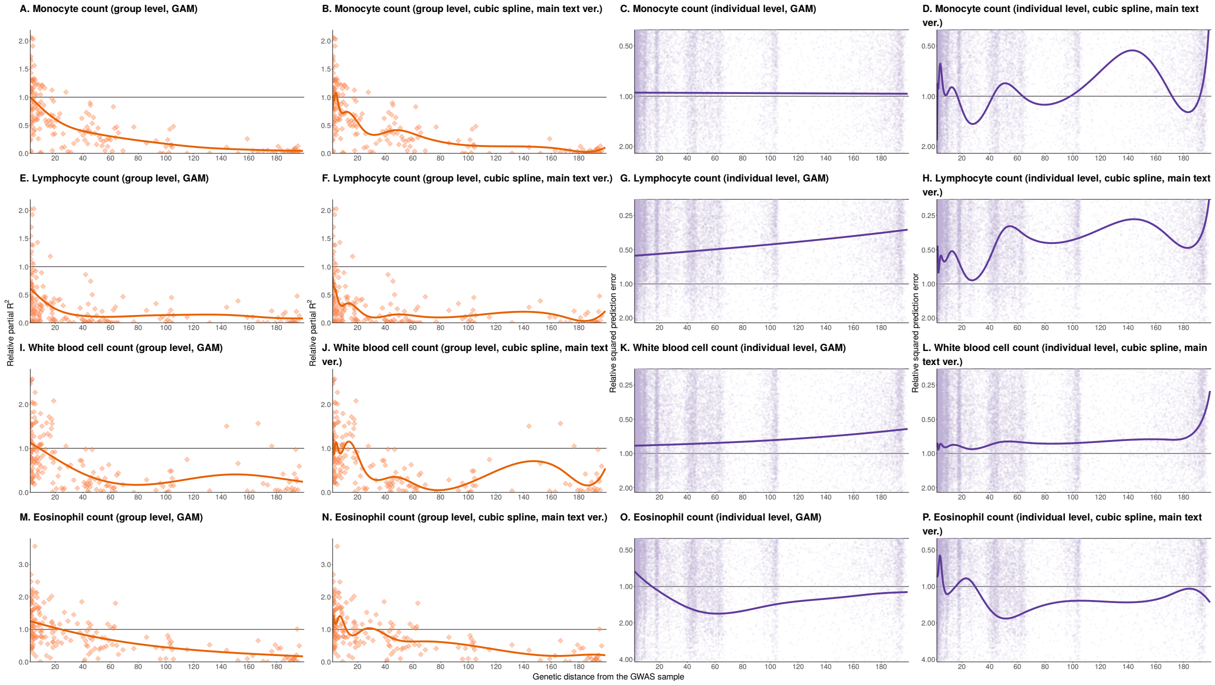

Figure S43: Trends of portability visualized with generalized additive model (GAM) fit, for white blood-cell related traits. This figure presents the same analysis as **Fig. 2** and **Fig. S3**, but the first and third columns (**A, C, E, G, I, K, M, O**) are plots with GAM fits, as shown by the curves. The second and fourth columns (**B, D, F, H, J, L, N, P**) contain plots with cubic spline fits, taken from the main text or supplementary text for comparison. At the group level (left two columns), we measured prediction accuracy with the squared partial correlation between the PGS and the trait value in 250 bins of 278 individuals each. At the individual level (right two columns), we measured prediction accuracy as the squared difference between the predicted phenotype and the true phenotype value. At both the group and individual levels, y-axis values show relative prediction accuracy, i.e. prediction accuracy divided by a baseline value. The baseline value is the mean prediction accuracy in 25 bins with median genetic distances that are closest to the mean genetic distance for GWAS individuals. **Table S1** details these baseline values for each trait. For both GAM and cubic spline fits, curves show cubic spline fits, with 8 knots placed based on the density of data points.

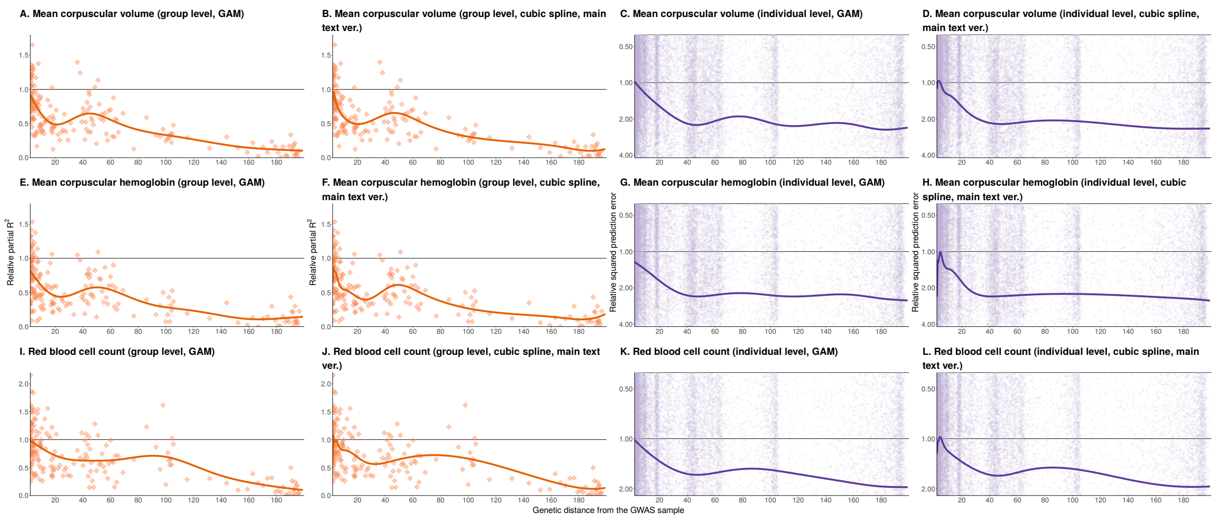

Figure S44: Trends of portability visualized with generalized additive model (GAM) fit, for red blood cell-related traits. This figure presents the same analysis as **Fig. 2** and **Fig. S4**, but the first and third columns (**A, C, E, G, I, K**) are plots with GAM fits, as shown by the curves. The second and fourth columns (**B, D, F, H, J, L**) contain plots with cubic spline fits, taken from the main text or supplementary text for comparison. At the group level (left two columns), we measured prediction accuracy with the squared partial correlation between the PGS and the trait value in 250 bins of 278 individuals each. At the individual level (right two columns), we measured prediction accuracy as the squared difference between the predicted phenotype and the true phenotype value. At both the group and individual levels, y-axis values show relative prediction accuracy, i.e. prediction accuracy divided by a baseline value. The baseline value is the mean prediction accuracy in 25 bins with median genetic distances that are closest to the mean genetic distance for GWAS individuals. **Table S1** details these baseline values for each trait. For both GAM and cubic spline fits, curves show cubic spline fits, with 8 knots placed based on the density of data points.

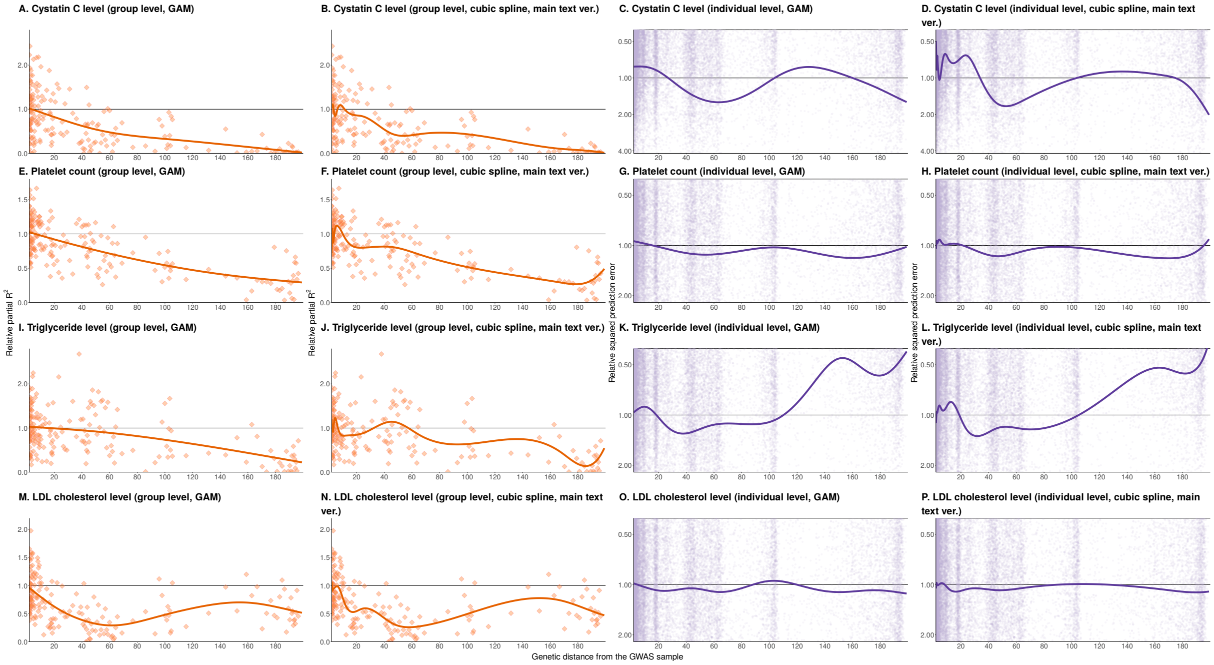

Figure S45: Trends of portability visualized with generalized additive model (GAM) fit, for other biomarkers. This figure presents the same analysis as **Fig. 2** and **Fig. S5**, but the first and third columns (**A, C, E, G, I, K, M, O**) are plots with GAM fits, as shown by the curves. The second and fourth columns (**B, D, F, H, J, L, N, P**) contain plots with cubic spline fits, taken from the main text or supplementary text for comparison. At the group level (left two columns), we measured prediction accuracy with the squared partial correlation between the PGS and the trait value in 250 bins of 278 individuals each. At the individual level (right two columns), we measured prediction accuracy as the squared difference between the predicted phenotype and the true phenotype value. At both the group and individual levels, y-axis values show relative prediction accuracy, i.e. prediction accuracy divided by a baseline value. The baseline value is the mean prediction accuracy in 25 bins with median genetic distances that are closest to the mean genetic distance for GWAS individuals. **Table S1** details these baseline values for each trait. For both GAM and cubic spline fits, curves show cubic spline fits, with 8 knots placed based on the density of data points.

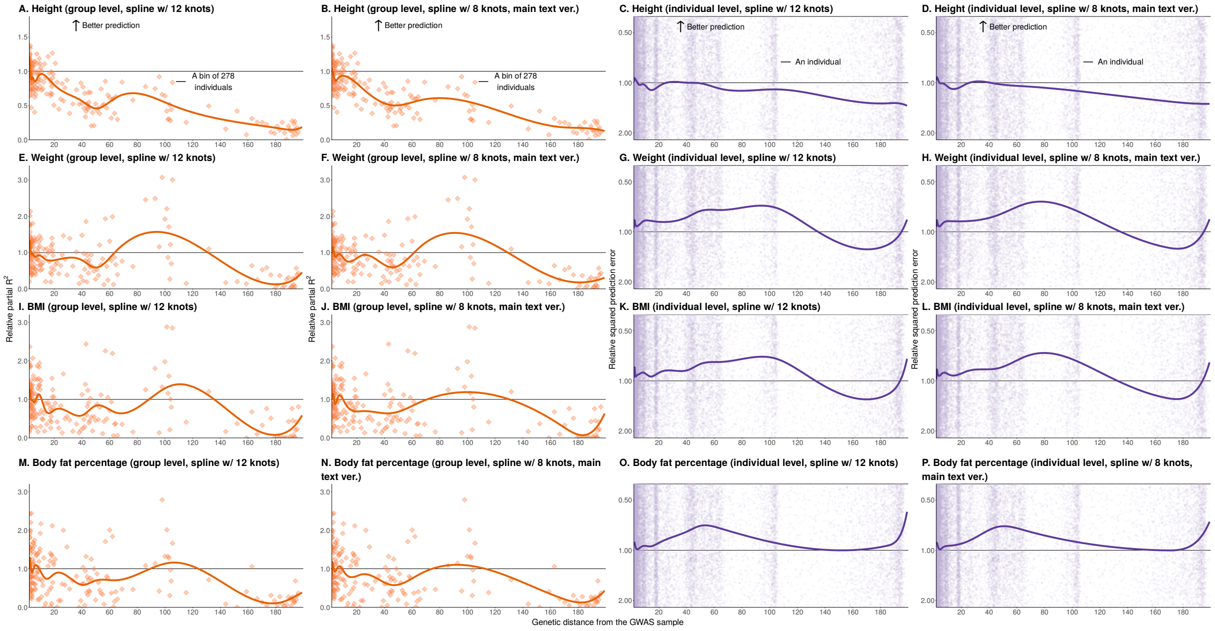

Figure S46: Trends of portability visualized using 12 knots in cubic line fits, for anthropometric measurements. This figure presents the same analysis as **Fig. 2** and **Fig. S2**, but the first and third columns (**A, C, E, G, I, K, M, O**) are plots with cubic splines of 12 knots. The second and fourth columns (**B, D, F, H, J, L, N, P**) contain plots with cubic splines of 8 knots, taken from the main text or supplementary text for comparison. For both knot numbers, the knots were placed based on the density of data points. At the group level (left two columns), we measured prediction accuracy with the squared partial correlation between the PGS and the trait value in 250 bins of 278 individuals each. At the individual level (right two columns), we measured prediction accuracy as the squared difference between the predicted phenotype and the true phenotype value. At both the group and individual levels, y-axis values show relative prediction accuracy, i.e. prediction accuracy divided by a baseline value. The baseline value is the mean prediction accuracy in 25 bins with median genetic distances that are closest to the mean genetic distance for GWAS individuals. **Table S1** details these baseline values for each trait.

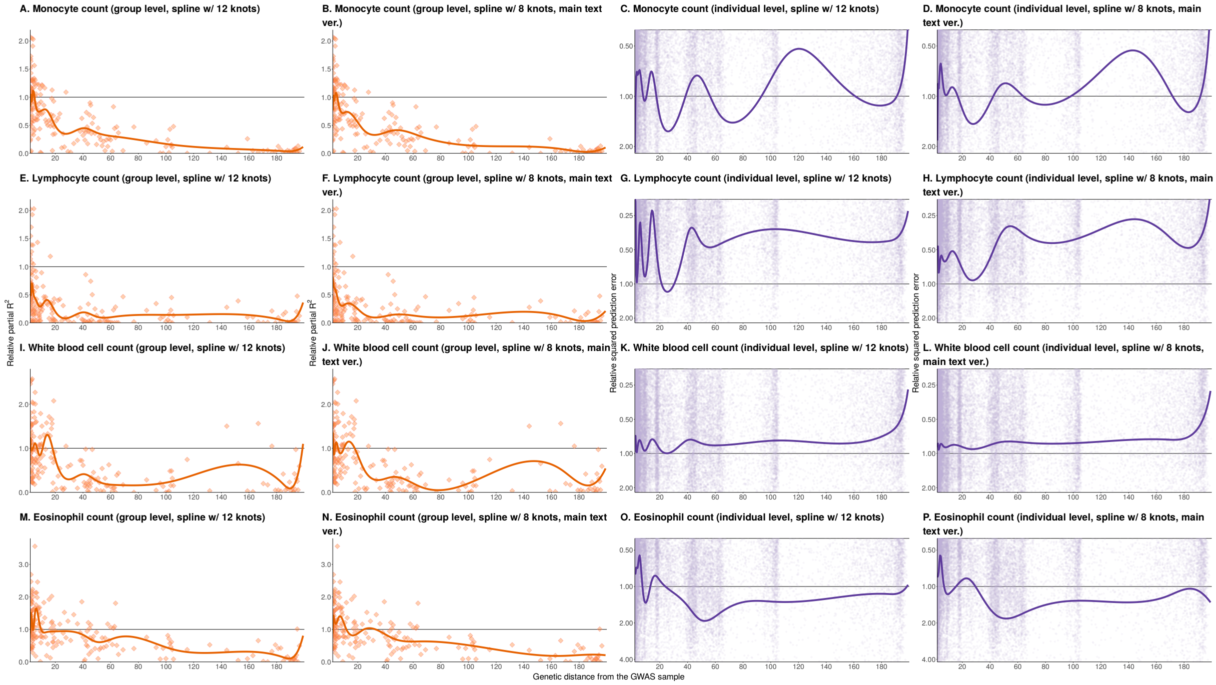

Figure S47: Trends of portability visualized using 12 knots in cubic line fits, for white blood-cell related traits. This figure presents the same analysis as **Fig. 2** and **Fig. S3**, but the first and third columns (**A, C, E, G, I, K, M, O**) are plots with cubic splines of 12 knots. The second and fourth columns (**B, D, F, H, J, L, N, P**) contain plots with cubic splines of 8 knots, taken from the main text or supplementary text for comparison. For both knot numbers, the knots were placed based on the density of data points. At the group level (left two columns), we measured prediction accuracy with the squared partial correlation between the PGS and the trait value in 250 bins of 278 individuals each. At the individual level (right two columns), we measured prediction accuracy as the squared difference between the predicted phenotype and the true phenotype value. At both the group and individual levels, y-axis values show relative prediction accuracy, i.e. prediction accuracy divided by a baseline value. The baseline value is the mean prediction accuracy in 25 bins with median genetic distances that are closest to the mean genetic distance for GWAS individuals. **Table S1** details these baseline values for each trait.

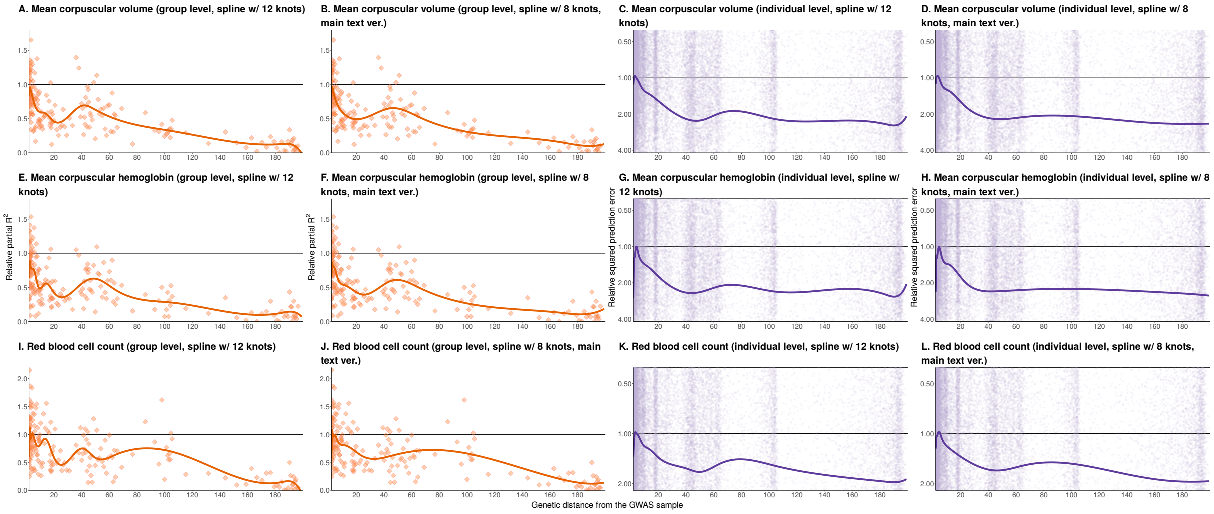

Figure S48: Trends of portability visualized using 12 knots in cubic line fits, for red blood cell-related traits. This figure presents the same analysis as **Fig. 2** and **Fig. S4**, but the first and third columns (**A, C, E, G, I, K**) are plots with cubic splines of 12 knots. The second and fourth columns (**B, D, F, H, J, L**) contain plots with cubic splines of 8 knots, taken from the main text or supplementary text for comparison. For both knot numbers, the knots were placed based on the density of data points. At the group level (left two columns), we measured prediction accuracy with the squared partial correlation between the PGS and the trait value in 250 bins of 278 individuals each. At the individual level (right two columns), we measured prediction accuracy as the squared difference between the predicted phenotype and the true phenotype value. At both the group and individual levels, y-axis values show relative prediction accuracy, i.e. prediction accuracy divided by a baseline value. The baseline value is the mean prediction accuracy in 25 bins with median genetic distances that are closest to the mean genetic distance for GWAS individuals. **Table S1** details these baseline values for each trait.

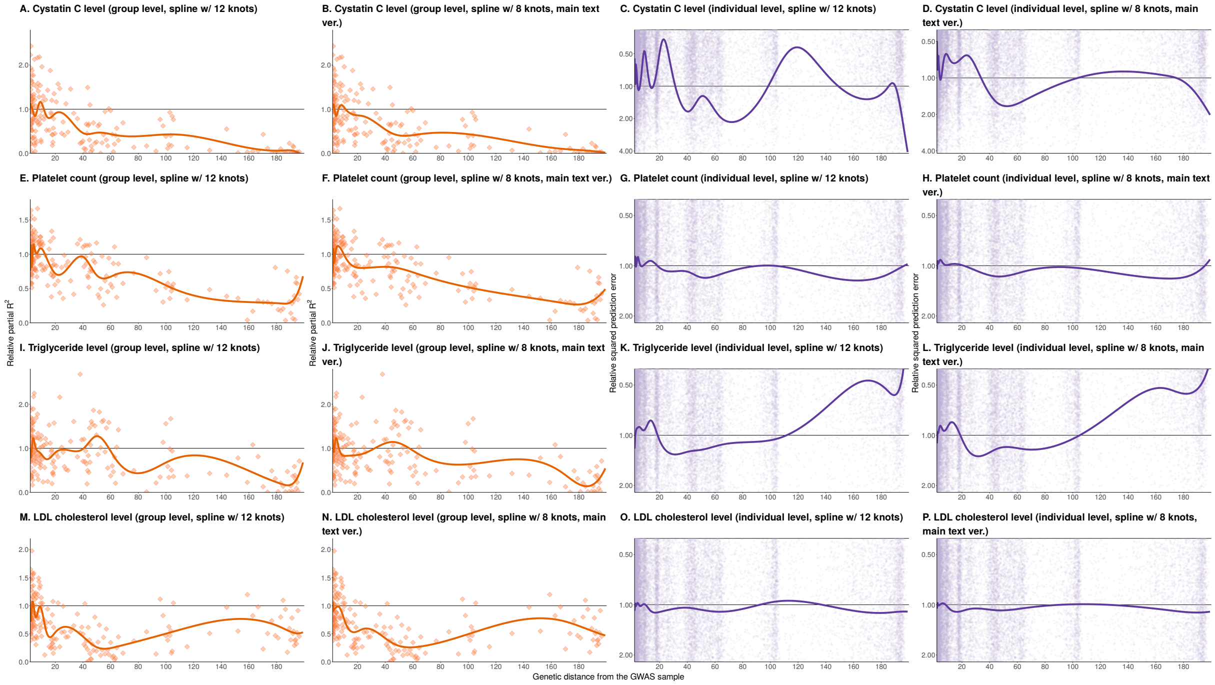

Figure S49: Trends of portability visualized using 12 knots in cubic line fits, for other biomarkers. This figure presents the same analysis as **Fig. 2** and **Fig. S5**, but the first and third columns (**A, C, E, G, I, K, M, O**) are plots with cubic splines of 12 knots. The second and fourth columns (**B, D, F, H, J, L, N, P**) contain plots with cubic splines of 8 knots, taken from the main text or supplementary text for comparison. For both knot numbers, the knots were placed based on the density of data points. At the group level (left two columns), we measured prediction accuracy with the squared partial correlation between the PGS and the trait value in 250 bins of 278 individuals each. At the individual level (right two columns), we measured prediction accuracy as the squared difference between the predicted phenotype and the true phenotype value. At both the group and individual levels, y-axis values show relative prediction accuracy, i.e. prediction accuracy divided by a baseline value. The baseline value is the mean prediction accuracy in 25 bins with median genetic distances that are closest to the mean genetic distance for GWAS individuals. **Table S1** details these baseline values for each trait.

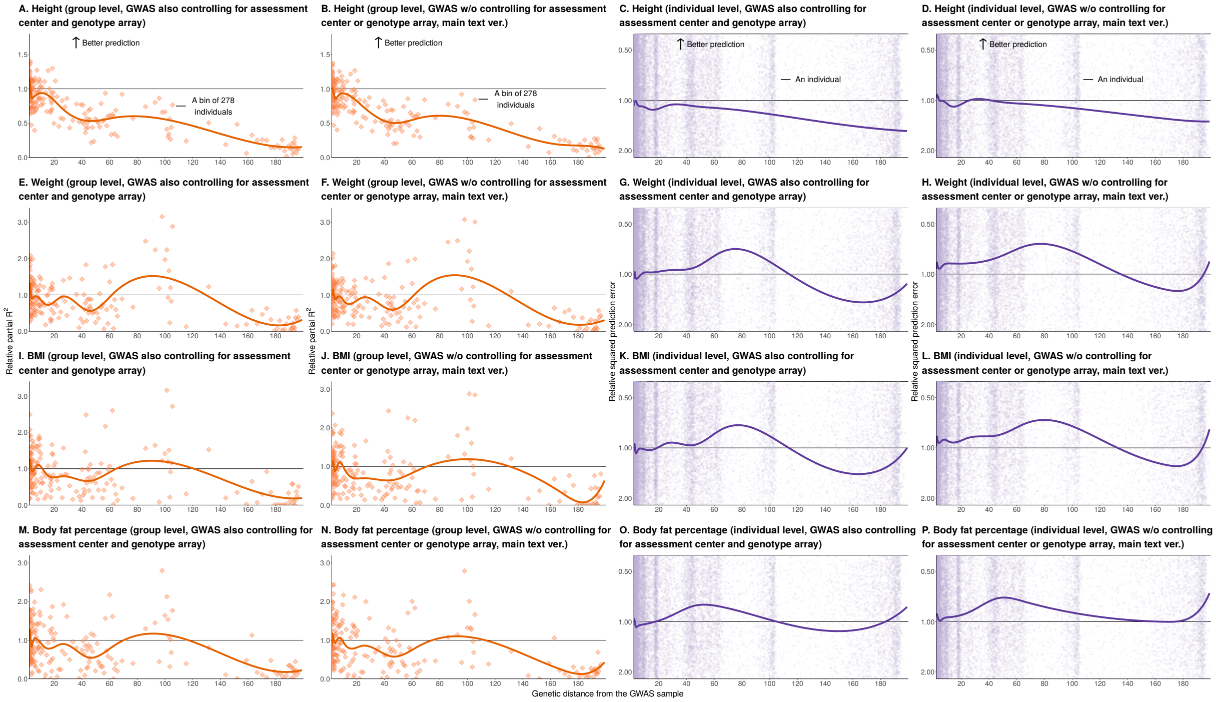

Figure S50: Trends of portability when including additional GWAS covariates, for anthropometric measurements. This figure presents the same analysis as **Fig. 2** and **Fig. S2**, but the first and third columns (**A, C, E, G, I, K, M, O**) are plots controlling for assessment center and genotype array in the GWAS. The second and fourth columns (**B, D, F, H, J, L, N, P**) contain plots without controlling for assessment center and genotype array in the GWAS, taken from the main text or supplementary text for comparison. At the group level (left two columns), we measured prediction accuracy with the squared partial correlation between the PGS and the trait value in 250 bins of 278 individuals each. At the individual level (right two columns), we measured prediction accuracy as the squared difference between the predicted phenotype and the true phenotype value. At both the group and individual levels, y-axis values show relative prediction accuracy, i.e. prediction accuracy divided by a baseline value. The baseline value is the mean prediction accuracy in 25 bins with median genetic distances that are closest to the mean genetic distance for GWAS individuals. **Table S1** details these baseline values for each trait. Curves show cubic spline fits, with 8 knots placed based on the density of data points.

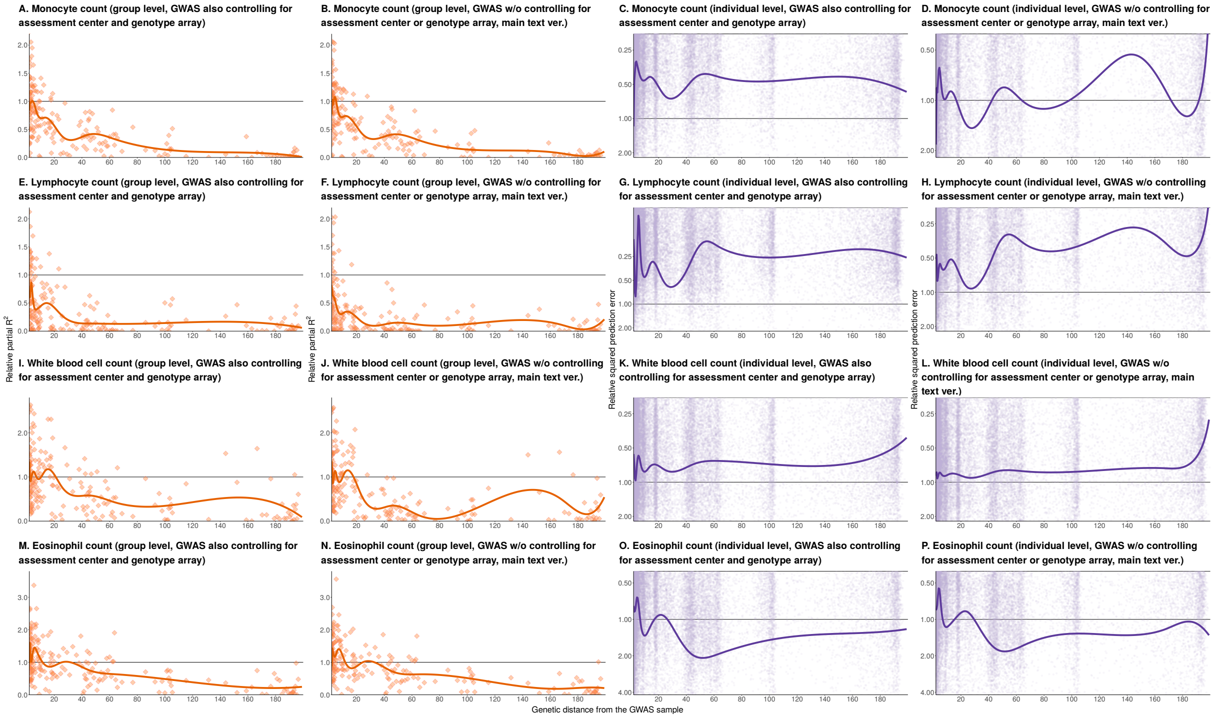

Figure S51: Trends of portability when including additional GWAS covariates, for white blood-cell related traits. This figure presents the same analysis as **Fig. 2** and **Fig. S2**, but the first and third columns (**A, C, E, G, I, K, M, O**) are plots controlling for assessment center and genotype array in the GWAS. The second and fourth columns (**B, D, F, H, J, L, N, P**) contain plots without controlling for assessment center and genotype array in the GWAS, taken from the main text or supplementary text for comparison. At the group level (left two columns), we measured prediction accuracy with the squared partial correlation between the PGS and the trait value in 250 bins of 278 individuals each. At the individual level (right two columns), we measured prediction accuracy as the squared difference between the predicted phenotype and the true phenotype value. At both the group and individual levels, y-axis values show relative prediction accuracy, i.e. prediction accuracy divided by a baseline value. The baseline value is the mean prediction accuracy in 25 bins with median genetic distances that are closest to the mean genetic distance for GWAS individuals. **Table S1** details these baseline values for each trait. Curves show cubic spline fits, with 8 knots placed based on the density of data points.

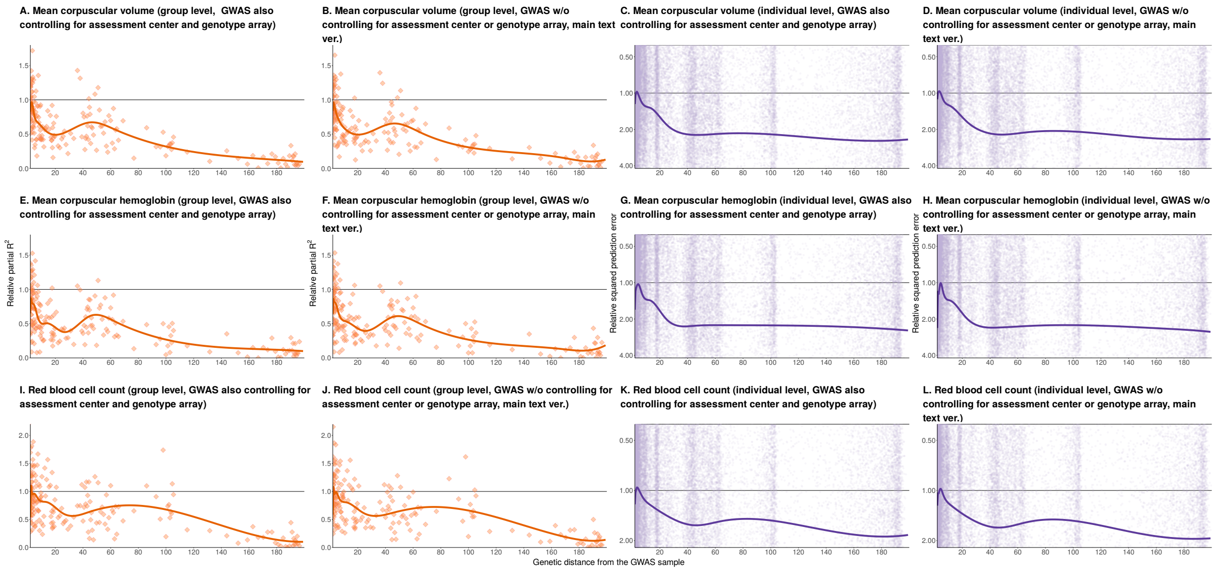

Figure S52: Trends of portability when including additional GWAS covariates, for red blood cell-related traits. This figure presents the same analysis as **Fig. 2** and **Fig. S2**, but the first and third columns (**A, C, E, G, I, K, M, O**) are plots controlling for assessment center and genotype array in the GWAS. The second and fourth columns (**B, D, F, H, J, L, N, P**) contain plots without controlling for assessment center and genotype array in the GWAS, taken from the main text or supplementary text for comparison. At the group level (left two columns), we measured prediction accuracy with the squared partial correlation between the PGS and the trait value in 250 bins of 278 individuals each. At the individual level (right two columns), we measured prediction accuracy as the squared difference between the predicted phenotype and the true phenotype value. At both the group and individual levels, y-axis values show relative prediction accuracy, i.e. prediction accuracy divided by a baseline value. The baseline value is the mean prediction accuracy in 25 bins with median genetic distances that are closest to the mean genetic distance for GWAS individuals. **Table S1** details these baseline values for each trait. Curves show cubic spline fits, with 8 knots placed based on the density of data points.

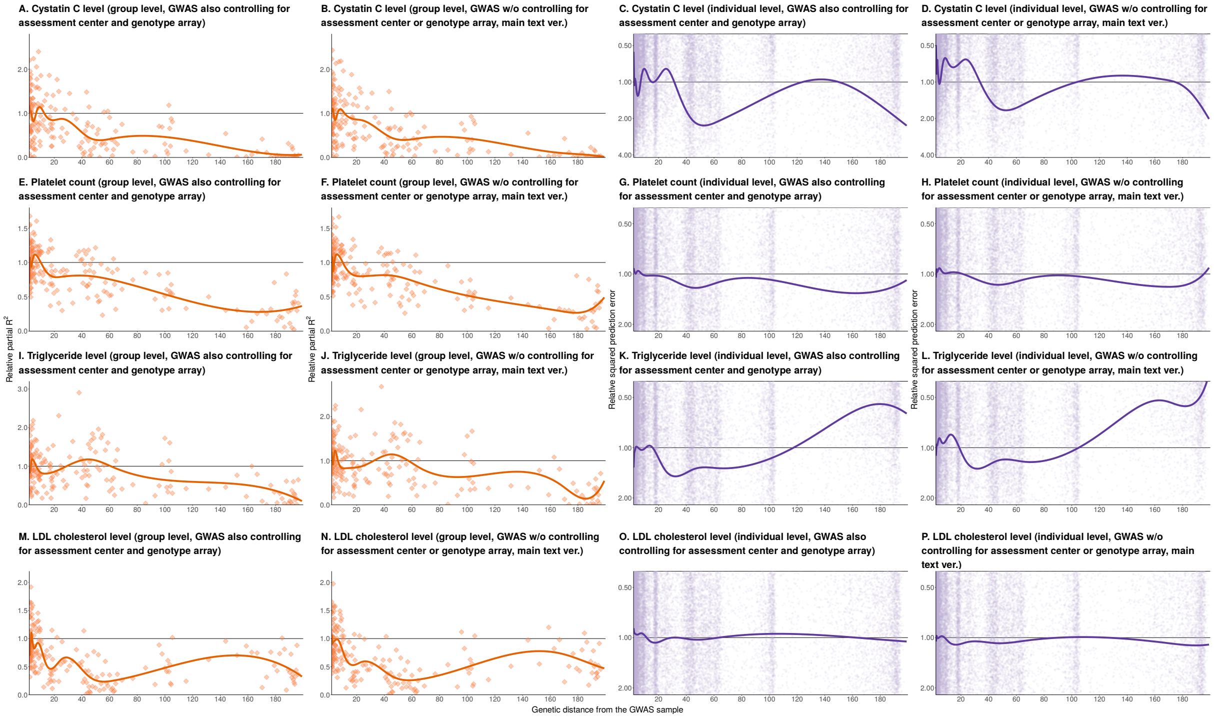

Figure S53: Trends of portability when including additional GWAS covariates, for other biomarkers. This figure presents the same analysis as **Fig. 2** and **Fig. S2**, but the first and third columns (**A, C, E, G, I, K, M, O**) are plots controlling for assessment center and genotype array in the GWAS. The second and fourth columns (**B, D, F, H, J, L, N, P**) contain plots without controlling for assessment center and genotype array in the GWAS, taken from the main text or supplementary text for comparison. At the group level (left two columns), we measured prediction accuracy with the squared partial correlation between the PGS and the trait value in 250 bins of 278 individuals each. At the individual level (right two columns), we measured prediction accuracy as the squared difference between the predicted phenotype and the true phenotype value. At both the group and individual levels, y-axis values show relative prediction accuracy, i.e. prediction accuracy divided by a baseline value. The baseline value is the mean prediction accuracy in 25 bins with median genetic distances that are closest to the mean genetic distance for GWAS individuals. **Table S1** details these baseline values for each trait. Curves show cubic spline fits, with 8 knots placed based on the density of data points.

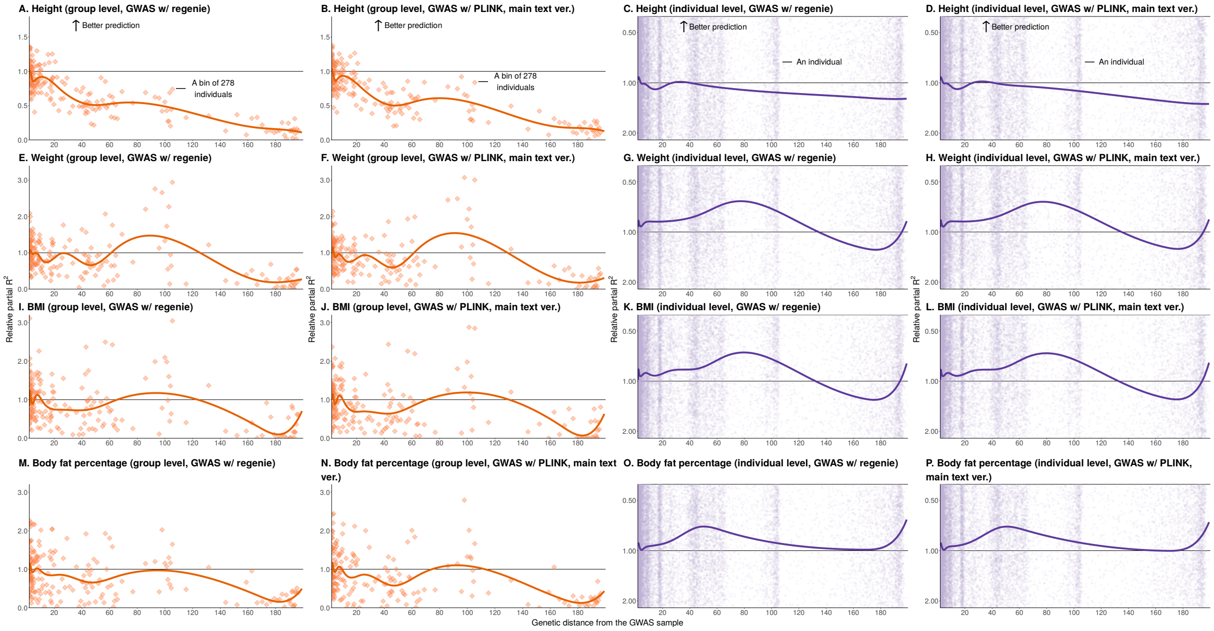

Figure S54: Trends of portability with GWAS run using *regenie*, for anthropometric measurements. This figure presents the same analysis as **Fig. 2** and **Fig. S2**, but the first and third columns (**A, C, E, G, I, K, M, O**) are plots with GWAS run with *regenie*. The second and fourth columns (**B, D, F, H, J, L, N, P**) contain plots with GWAS run with *PLINK*, taken from the main text or supplementary text for comparison. At the group level (left two columns), we measured prediction accuracy with the squared partial correlation between the PGS and the trait value in 250 bins of 278 individuals each. At the individual level (right two columns), we measured prediction accuracy as the squared difference between the predicted phenotype and the true phenotype value. At both the group and individual levels, y-axis values show relative prediction accuracy, i.e. prediction accuracy divided by a baseline value. The baseline value is the mean prediction accuracy in 25 bins with median genetic distances that are closest to the mean genetic distance for GWAS individuals. **Table S1** details these baseline values for each trait. Curves show cubic spline fits, with 8 knots placed based on the density of data points.

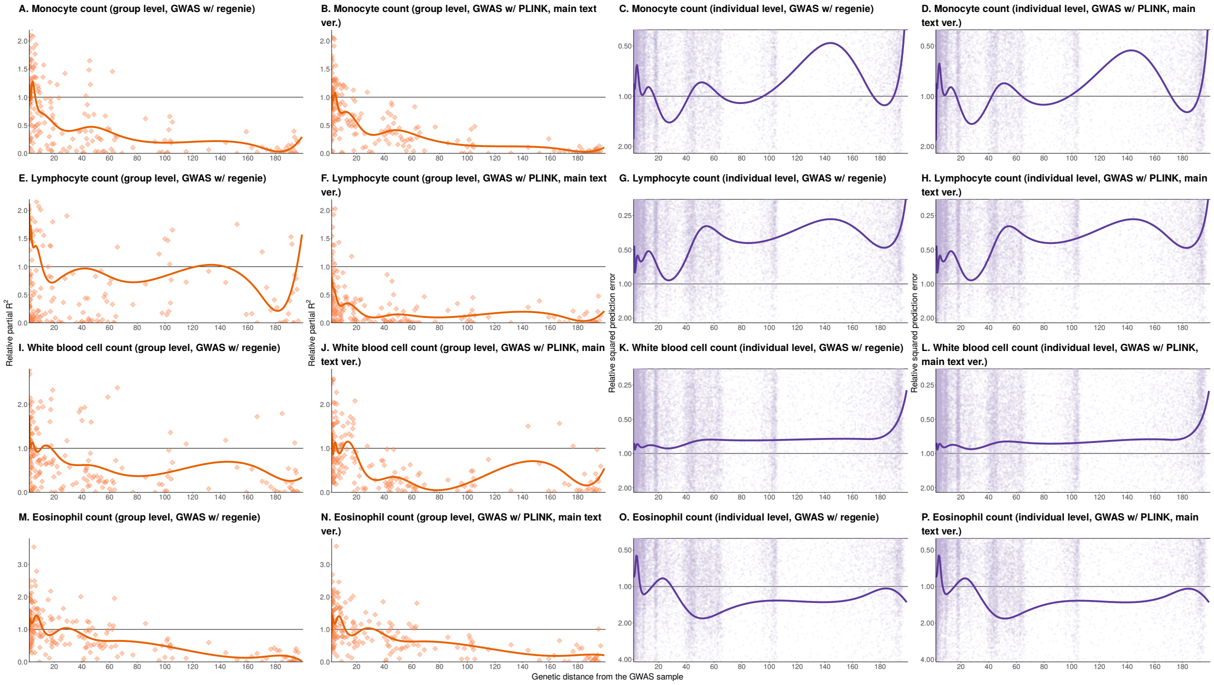

Figure S55: Trends of portability with GWAS run using *regenie*, for white blood-cell related traits. This figure presents the same analysis as **Fig. 2** and **Fig. S2**, but the first and third columns (**A, C, E, G, I, K, M, O**) are plots with GWAS run with *regenie*. The second and fourth columns (**B, D, F, H, J, L, N, P**) contain plots with GWAS run with *PLINK*, taken from the main text or supplementary text for comparison. At the group level (left two columns), we measured prediction accuracy with the squared partial correlation between the PGS and the trait value in 250 bins of 278 individuals each. At the individual level (right two columns), we measured prediction accuracy as the squared difference between the predicted phenotype and the true phenotype value. At both the group and individual levels, y-axis values show relative prediction accuracy, i.e. prediction accuracy divided by a baseline value. The baseline value is the mean prediction accuracy in 25 bins with median genetic distances that are closest to the mean genetic distance for GWAS individuals. **Table S1** details these baseline values for each trait. Curves show cubic spline fits, with 8 knots placed based on the density of data points.

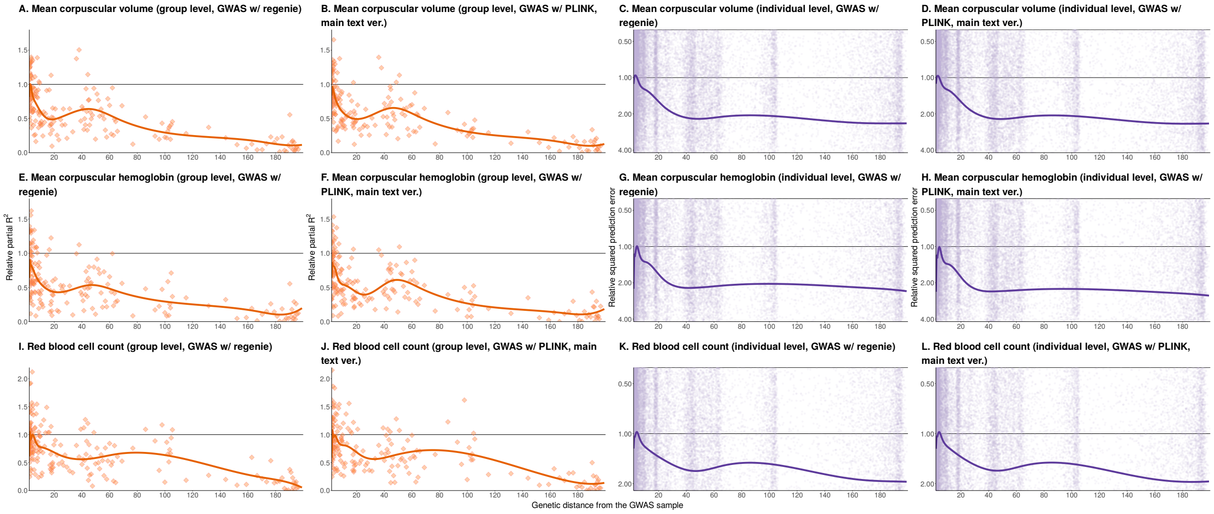

Figure S56: Trends of portability with GWAS run using *regenie*, for red blood cell-related traits. This figure presents the same analysis as **Fig. 2** and **Fig. S2**, but the first and third columns (**A, C, E, G, I, K, M, O**) are plots with GWAS run with *regenie*. The second and fourth columns (**B, D, F, H, J, L, N, P**) contain plots with GWAS run with *PLINK*, taken from the main text or supplementary text for comparison. At the group level (left two columns), we measured prediction accuracy with the squared partial correlation between the PGS and the trait value in 250 bins of 278 individuals each. At the individual level (right two columns), we measured prediction accuracy as the squared difference between the predicted phenotype and the true phenotype value. At both the group and individual levels, y-axis values show relative prediction accuracy, i.e. prediction accuracy divided by a baseline value. The baseline value is the mean prediction accuracy in 25 bins with median genetic distances that are closest to the mean genetic distance for GWAS individuals. **Table S1** details these baseline values for each trait. Curves show cubic spline fits, with 8 knots placed based on the density of data points.

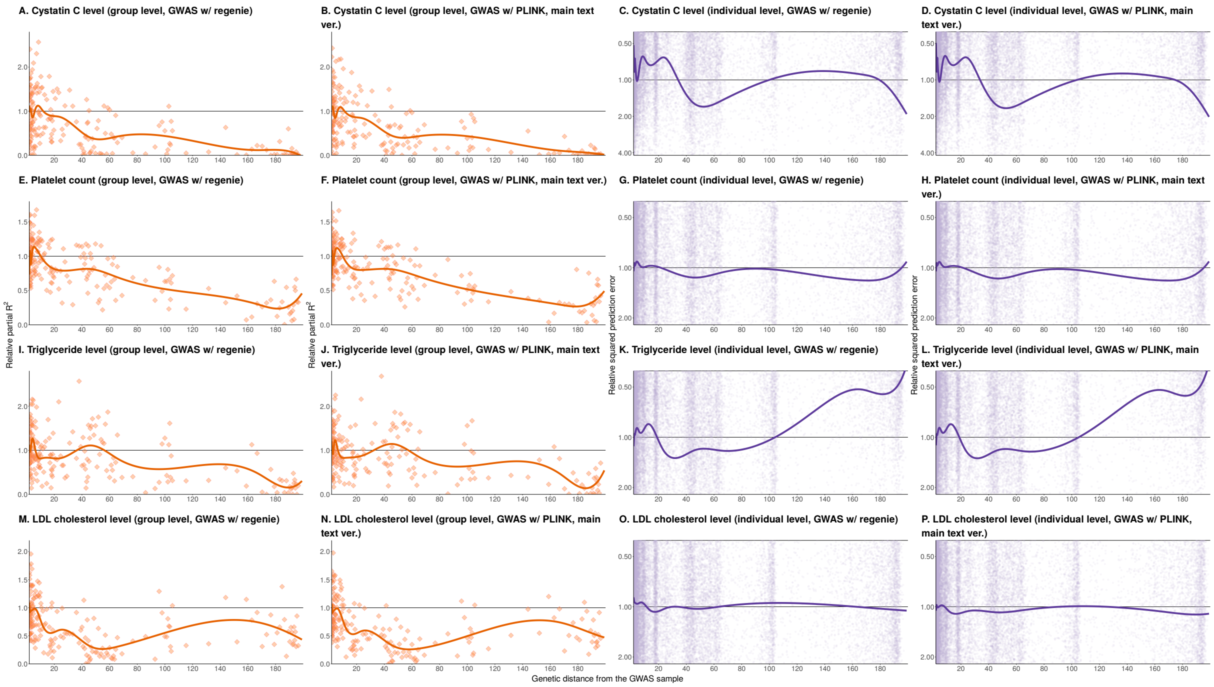

Figure S57: Trends of portability with GWAS run using *regenie*, for other biomarkers. This figure presents the same analysis as **Fig. 2** and **Fig. S2**, but the first and third columns (**A, C, E, G, I, K, M, O**) are plots with GWAS run with *regenie*. The second and fourth columns (**B, D, F, H, J, L, N, P**) contain plots with GWAS run with *PLINK*, taken from the main text or supplementary text for comparison. At the group level (left two columns), we measured prediction accuracy with the squared partial correlation between the PGS and the trait value in 250 bins of 278 individuals each. At the individual level (right two columns), we measured prediction accuracy as the squared difference between the predicted phenotype and the true phenotype value. At both the group and individual levels, y-axis values show relative prediction accuracy, i.e. prediction accuracy divided by a baseline value. The baseline value is the mean prediction accuracy in 25 bins with median genetic distances that are closest to the mean genetic distance for GWAS individuals. **Table S1** details these baseline values for each trait. Curves show cubic spline fits, with 8 knots placed based on the density of data points.

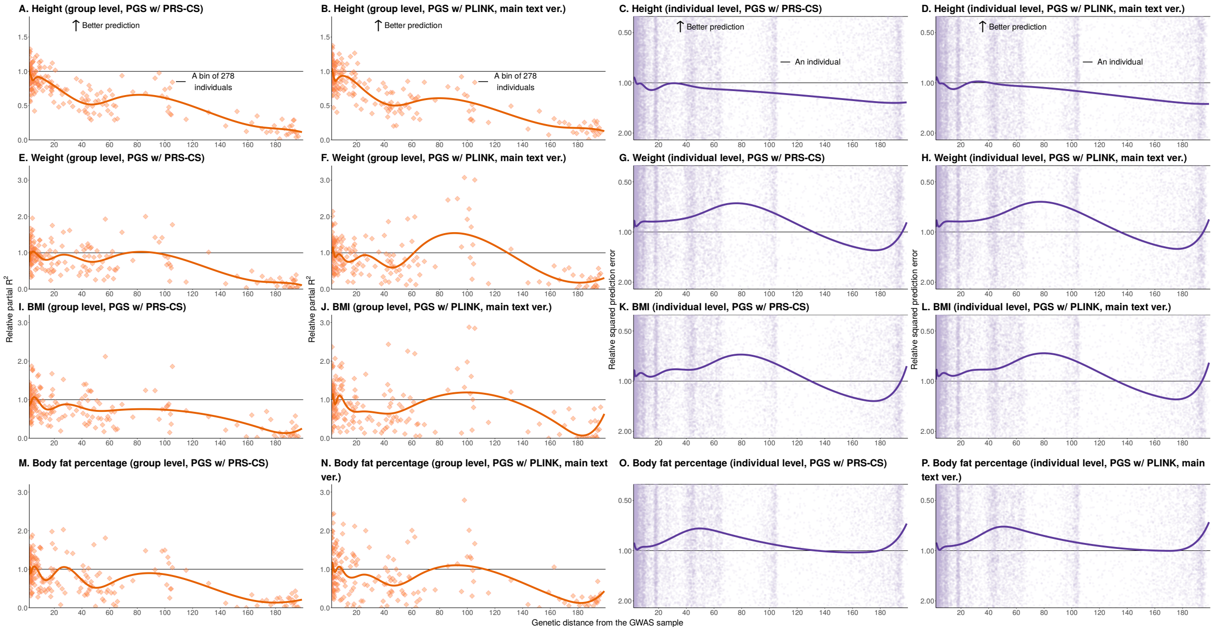

Figure S58: Trends of portability with PGS constructed using *PRS-CS*, for anthropometric measurements. This figure presents the same analysis as **Fig. 2** and **Fig. S2**, but the first and third columns (**A, C, E, G, I, K, M, O**) are plots with PRS reconstructed with *PRS-CS*. The second and fourth columns (**B, D, F, H, J, L, N, P**) contain plots with PGS constructed with *PLINK*, taken from the main text or supplementary text for comparison. At the group level (left two columns), we measured prediction accuracy with the squared partial correlation between the PGS and the trait value in 250 bins of 278 individuals each. At the individual level (right two columns), we measured prediction accuracy as the squared difference between the predicted phenotype and the true phenotype value. At both the group and individual levels, y-axis values show relative prediction accuracy, i.e. prediction accuracy divided by a baseline value. The baseline value is the mean prediction accuracy in 25 bins with median genetic distances that are closest to the mean genetic distance for GWAS individuals. **Table S1** details these baseline values for each trait. Curves show cubic spline fits, with 8 knots placed based on the density of data points.

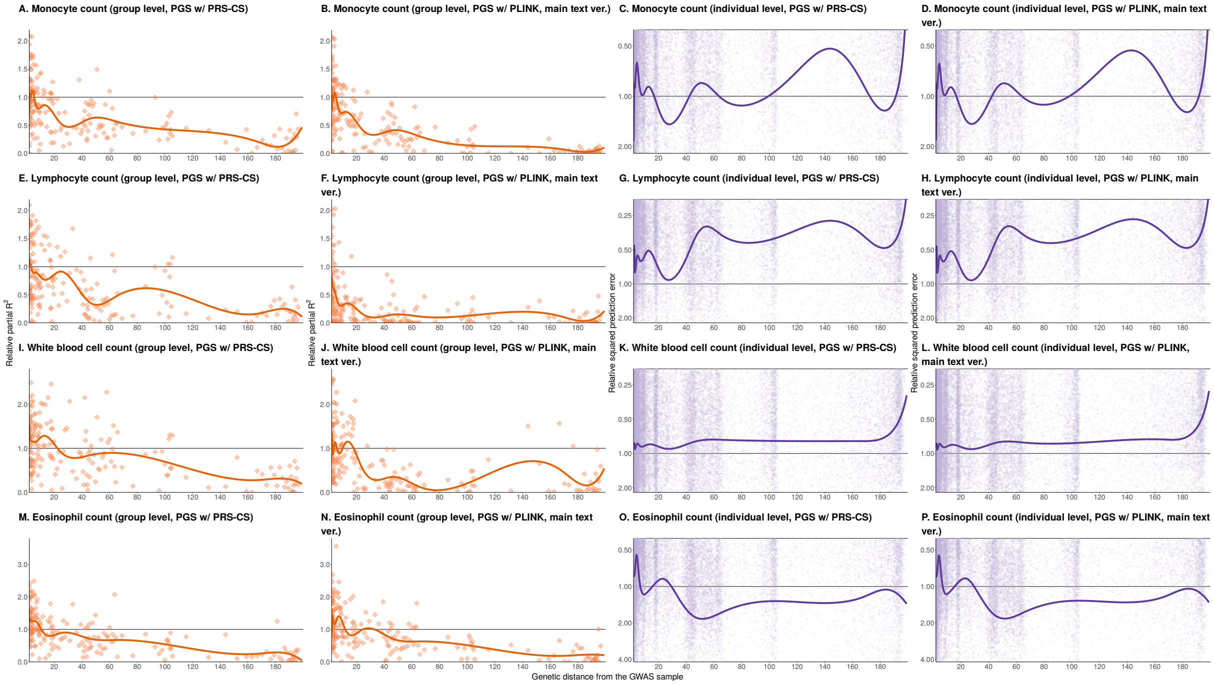

Figure S59: Trends of portability with PGS constructed using *PRS-CS*, for white blood-cell related traits. This figure presents the same analysis as **Fig. 2** and **Fig. S2**, but the first and third columns (**A, C, E, G, I, K, M, O**) are plots with PRS reconstructed with *PRS-CS*. The second and fourth columns (**B, D, F, H, J, L, N, P**) contain plots with PGS constructed with *PLINK*, taken from the main text or supplementary text for comparison. At the group level (left two columns), we measured prediction accuracy with the squared partial correlation between the PGS and the trait value in 250 bins of 278 individuals each. At the individual level (right two columns), we measured prediction accuracy as the squared difference between the predicted phenotype and the true phenotype value. At both the group and individual levels, y-axis values show relative prediction accuracy, i.e. prediction accuracy divided by a baseline value. The baseline value is the mean prediction accuracy in 25 bins with median genetic distances that are closest to the mean genetic distance for GWAS individuals. **Table S1** details these baseline values for each trait. Curves show cubic spline fits, with 8 knots placed based on the density of data points.

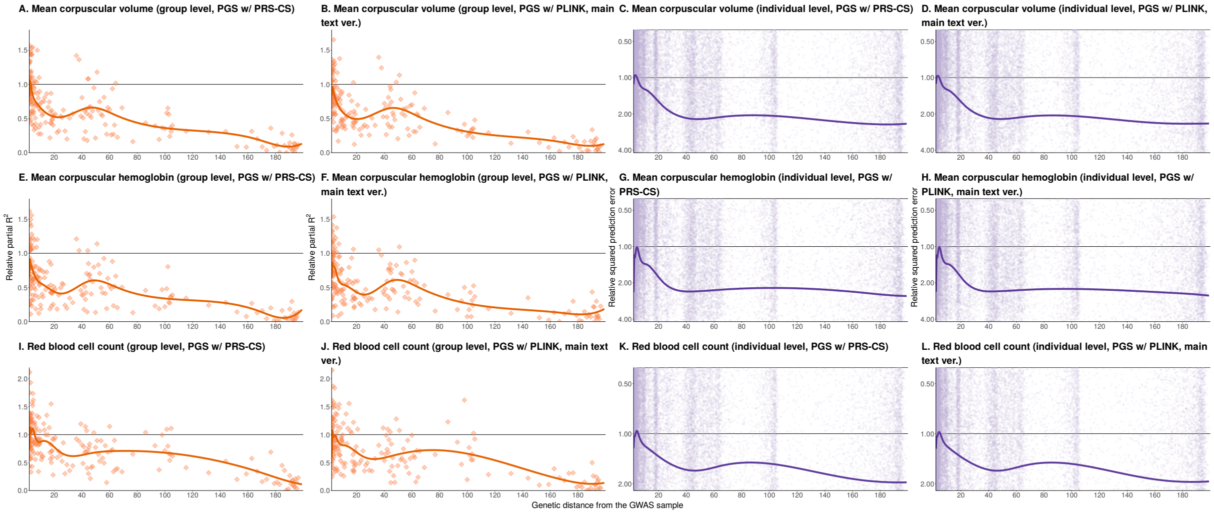

Figure S60: Trends of portability with PGS constructed using *PRS-CS*, for red blood cell-related traits. This figure presents the same analysis as **Fig. 2** and **Fig. S2**, but the first and third columns (**A, C, E, G, I, K, M, O**) are plots with PRS reconstructed with *PRS-CS*. The second and fourth columns (**B, D, F, H, J, L, N, P**) contain plots with PGS constructed with *PLINK*, taken from the main text or supplementary text for comparison. At the group level (left two columns), we measured prediction accuracy with the squared partial correlation between the PGS and the trait value in 250 bins of 278 individuals each. At the individual level (right two columns), we measured prediction accuracy as the squared difference between the predicted phenotype and the true phenotype value. At both the group and individual levels, y-axis values show relative prediction accuracy, i.e. prediction accuracy divided by a baseline value. The baseline value is the mean prediction accuracy in 25 bins with median genetic distances that are closest to the mean genetic distance for GWAS individuals. **Table S1** details these baseline values for each trait. Curves show cubic spline fits, with 8 knots placed based on the density of data points.

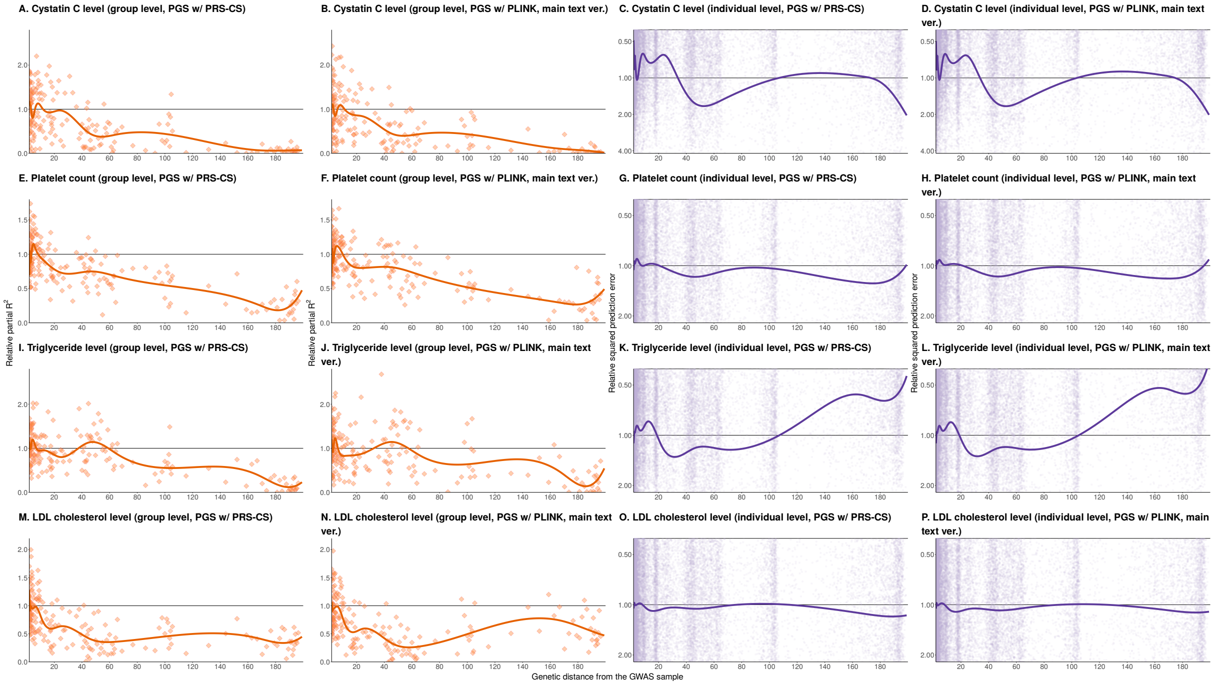

Figure S61: Trends of portability with PGS constructed using *PRS-CS*, for other biomarkers. This figure presents the same analysis as **Fig. 2** and **Fig. S2**, but the first and third columns (**A, C, E, G, I, K, M, O**) are plots with PRS reconstructed with *PRS-CS*. The second and fourth columns (**B, D, F, H, J, L, N, P**) contain plots with PGS constructed with *PLINK*, taken from the main text or supplementary text for comparison. At the group level (left two columns), we measured prediction accuracy with the squared partial correlation between the PGS and the trait value in 250 bins of 278 individuals each. At the individual level (right two columns), we measured prediction accuracy as the squared difference between the predicted phenotype and the true phenotype value. At both the group and individual levels, y-axis values show relative prediction accuracy, i.e. prediction accuracy divided by a baseline value. The baseline value is the mean prediction accuracy in 25 bins with median genetic distances that are closest to the mean genetic distance for GWAS individuals. **Table S1** details these baseline values for each trait. Curves show cubic spline fits, with 8 knots placed based on the density of data points.

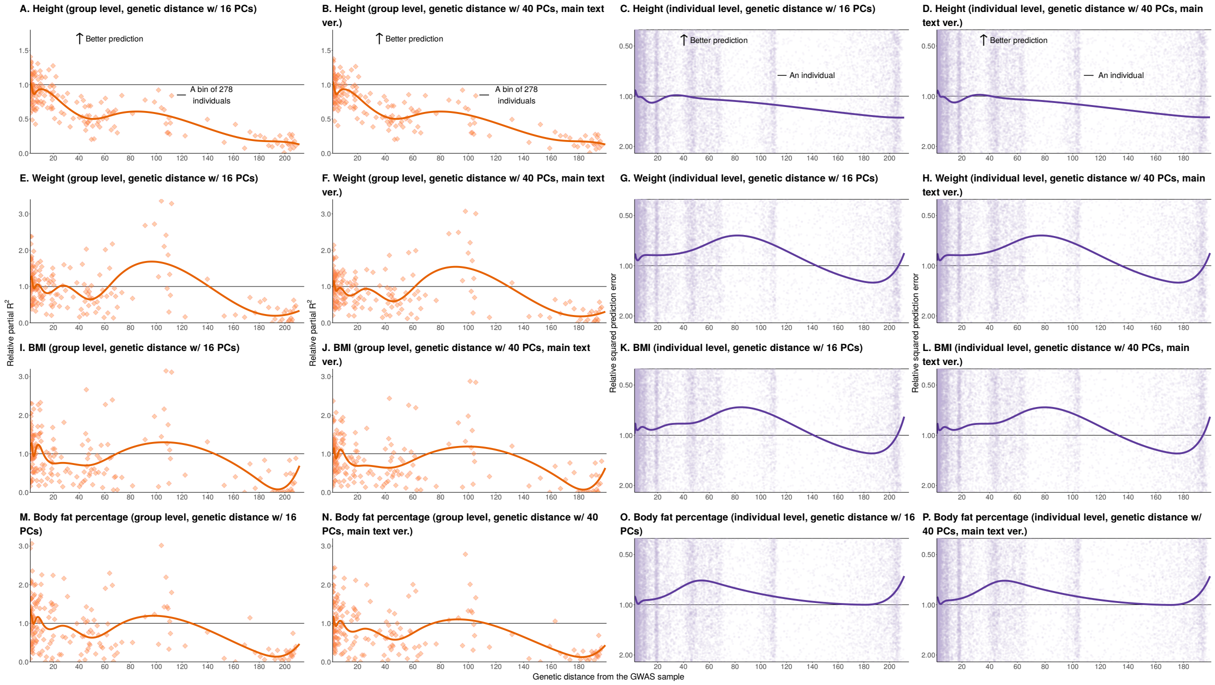

Figure S62: A comparison of portability trends, with genetic distance defined in two different ways, for anthropometric measurements. This figure presents the same analysis as **Fig. 2** and **Fig. S2**, but the first and third columns (**A, C, E, G, I, K, M, O**) show results obtained with a genetic distance metric defined using the first 16 PCs alone. The second and fourth columns (**B, D, F, H, J, L, N, P**) show results obtained with the measure of genetic distances that uses 40 PCs, as in the main text. At the group level (left two columns), we measured prediction accuracy with the squared partial correlation between the PGS and the trait value in 250 bins of 278 individuals each. At the individual level (right two columns), we measured prediction accuracy as the squared difference between the predicted phenotype and the true phenotype value. At both the group and individual levels, y-axis values show relative prediction accuracy, i.e. prediction accuracy divided by a baseline value. The baseline value is the mean prediction accuracy in 25 bins with median genetic distances that are closest to the mean genetic distance for GWAS individuals. **Table S1** details these baseline values for each trait. Curves show cubic spline fits, with 8 knots placed based on the density of data points.

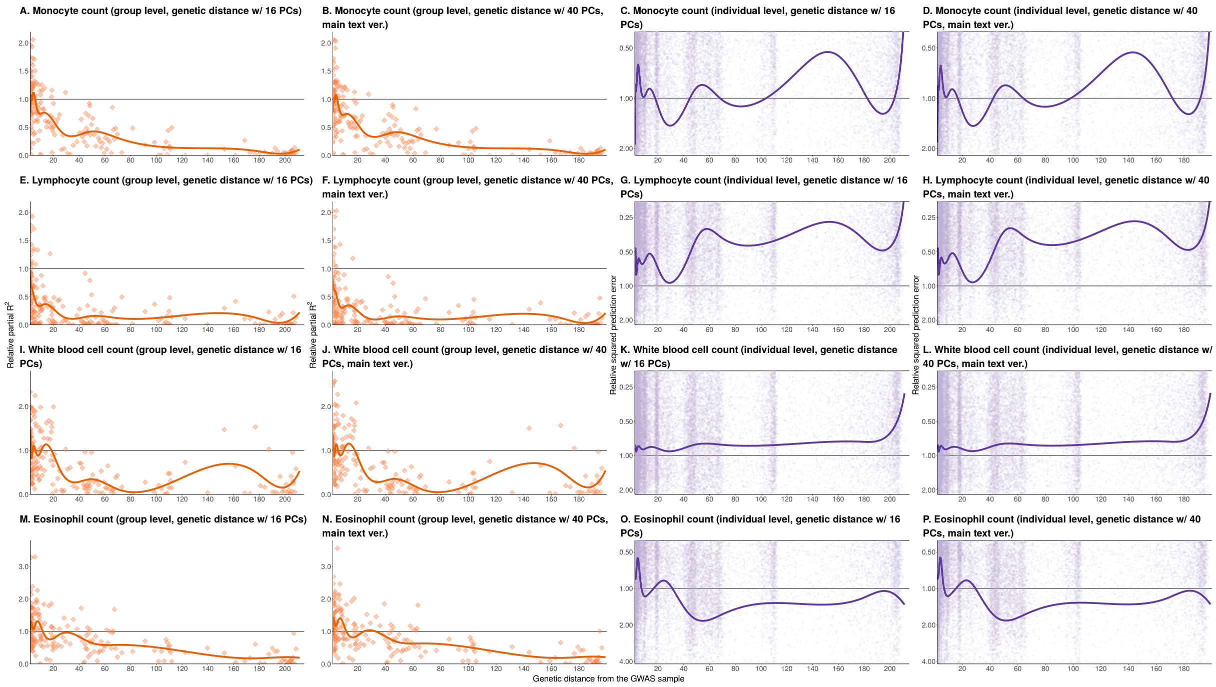

Figure S63: A comparison of portability trends, with genetic distance defined in two different ways, for white blood-cell related traits. This figure presents the same analysis as **Fig. 2** and **Fig. S3**, but the first and third columns (**A, C, E, G, I, K, M, O**) show results obtained with a genetic distance metric defined using the first 16 PCs alone. The second and fourth columns (**B, D, F, H, J, L, N, P**) show results obtained with the measure of genetic distances that uses 40 PCs, as in the main text. At the group level (left two columns), we measured prediction accuracy with the squared partial correlation between the PGS and the trait value in 250 bins of 278 individuals each. At the individual level (right two columns), we measured prediction accuracy as the squared difference between the predicted phenotype and the true phenotype value. At both the group and individual levels, y-axis values show relative prediction accuracy, i.e. prediction accuracy divided by a baseline value. The baseline value is the mean prediction accuracy in 25 bins with median genetic distances that are closest to the mean genetic distance for GWAS individuals. **Table S1** details these baseline values for each trait. Curves show cubic spline fits, with 8 knots placed based on the density of data points.

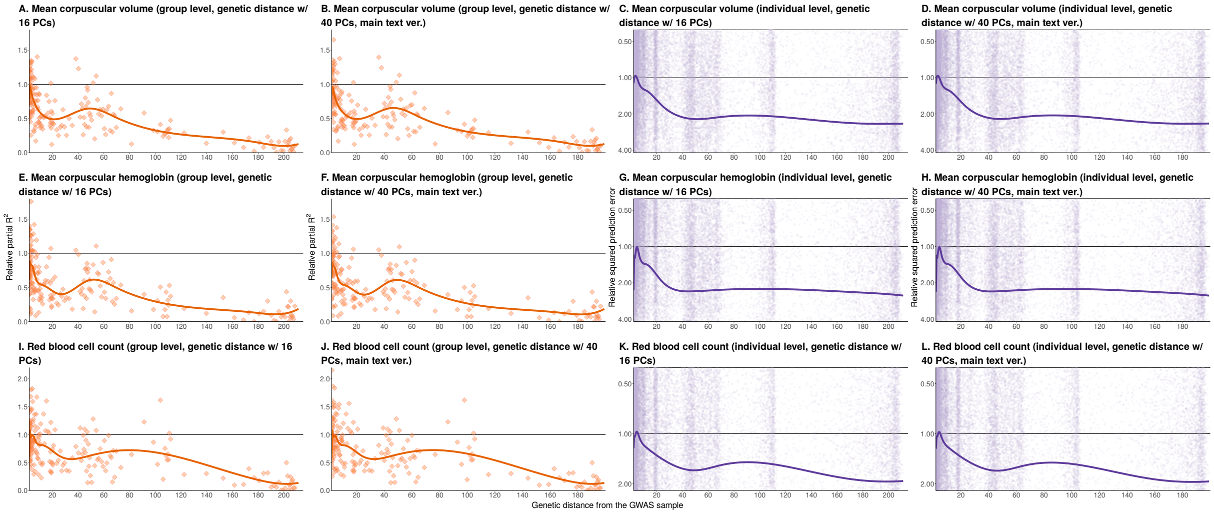

Figure S64: A comparison of portability trends, with genetic distance defined in two different ways, for red blood cell-related traits. This figure presents the same analysis as **Fig. 2** and **Fig. S4**, but the first and third columns (**A, C, E, G, I, K**) show results obtained with a genetic distance metric defined using the first 16 PCs alone. The second and fourth columns (**B, D, F, H, J, L**) show results obtained with the measure of genetic distances that uses 40 PCs, as in the main text. At the group level (left two columns), we measured prediction accuracy with the squared partial correlation between the PGS and the trait value in 250 bins of 278 individuals each. At the individual level (right two columns), we measured prediction accuracy as the squared difference between the predicted phenotype and the true phenotype value. At both the group and individual levels, y-axis values show relative prediction accuracy, i.e. prediction accuracy divided by a baseline value. The baseline value is the mean prediction accuracy in 25 bins with median genetic distances that are closest to the mean genetic distance for GWAS individuals. **Table S1** details these baseline values for each trait. Curves show cubic spline fits, with 8 knots placed based on the density of data points.

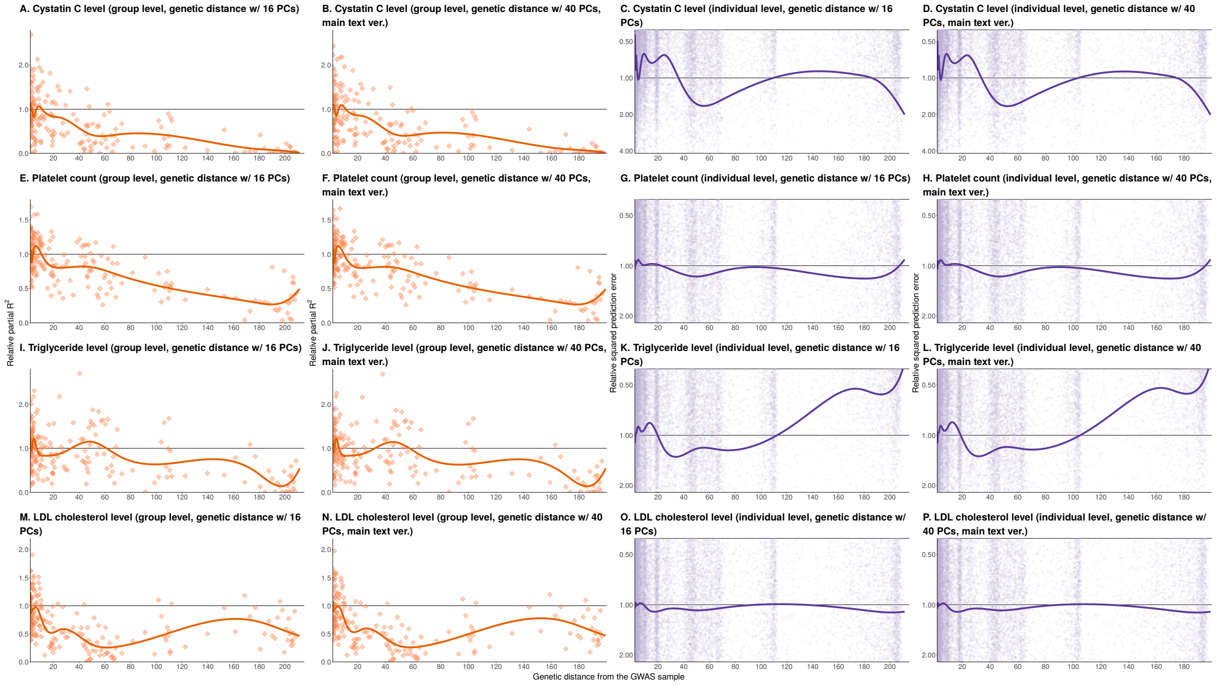

Figure S65: A comparison of portability trends, with genetic distance defined in two different ways, for other biomarkers. This figure presents the same analysis as **Fig. 2** and **Fig. S5**, but the first and third columns (**A, C, E, G, I, K, M, O**) show results obtained with a genetic distance metric defined using the first 16 PCs alone. The second and fourth columns (**B, D, F, H, J, L, N, P**) show results obtained with the measure of genetic distances that uses 40 PCs, as in the main text. At the group level (left two columns), we measured prediction accuracy with the squared partial correlation between the PGS and the trait value in 250 bins of 278 individuals each. At the individual level (right two columns), we measured prediction accuracy as the squared difference between the predicted phenotype and the true phenotype value. At both the group and individual levels, y-axis values show relative prediction accuracy, i.e. prediction accuracy divided by a baseline value. The baseline value is the mean prediction accuracy in 25 bins with median genetic distances that are closest to the mean genetic distance for GWAS individuals. **Table S1** details these baseline values for each trait. Curves show cubic spline fits, with 8 knots placed based on the density of data points.

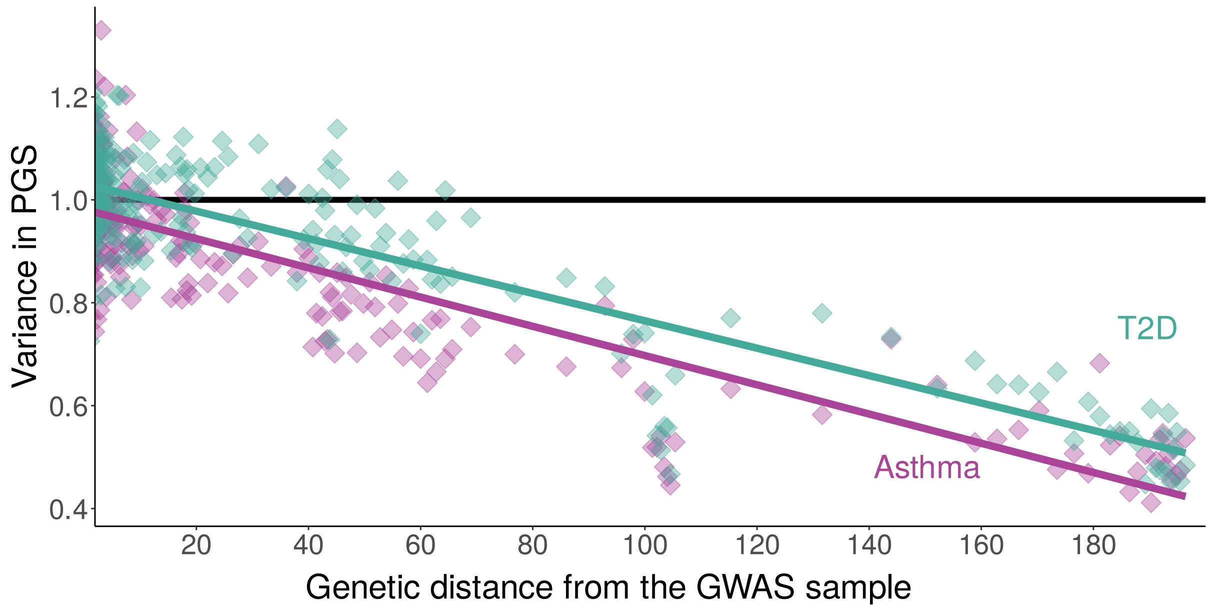

Figure S66: Variance in disease PGS as a function of genetic distance. Points show the PGS variance for each genetic distance bins. Lines show the ordinary least squares linear fit to the respective bin-level data.

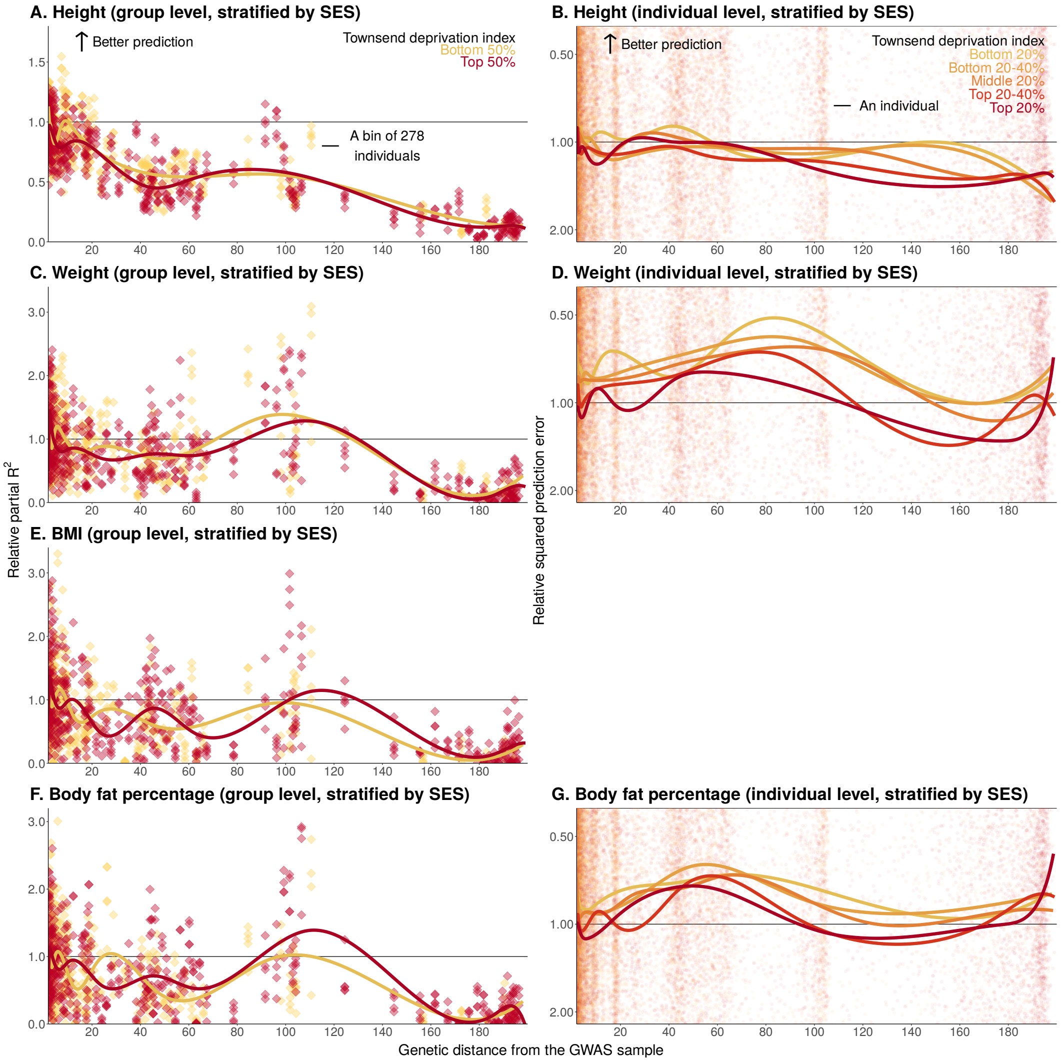

Figure S67: Portability as a function of both genetic distance and Townsend Deprivation Index, for anthropometric measurements. We first stratified the prediction sample individuals into equal-sized strata of Townsend Deprivation Index: 2 strata at the group level, shown on the left, and 5 equal-sized strata at the individual level, shown on the right panels. Within each stratum, we then performed the same analysis of prediction accuracy as a function of genetic distance as in in **Figs. 2, S2-S5**. At the group level, we measured prediction accuracy with the squared partial correlation between the PGS and the trait value in 250 bins of 278 individuals each. At the individual level, we measured prediction accuracy as the squared difference between the predicted phenotype and the true phenotype value. At both the group and individual levels, y-axis values show relative prediction accuracy, i.e. prediction accuracy divided by a baseline value. The baseline value is the mean prediction accuracy in 25 bins with median genetic distances that are closest to the mean genetic distance for GWAS individuals. Curves show cubic spline fits, with 8 knots placed based on the density of data points. "Top" represents those who are more deprived.

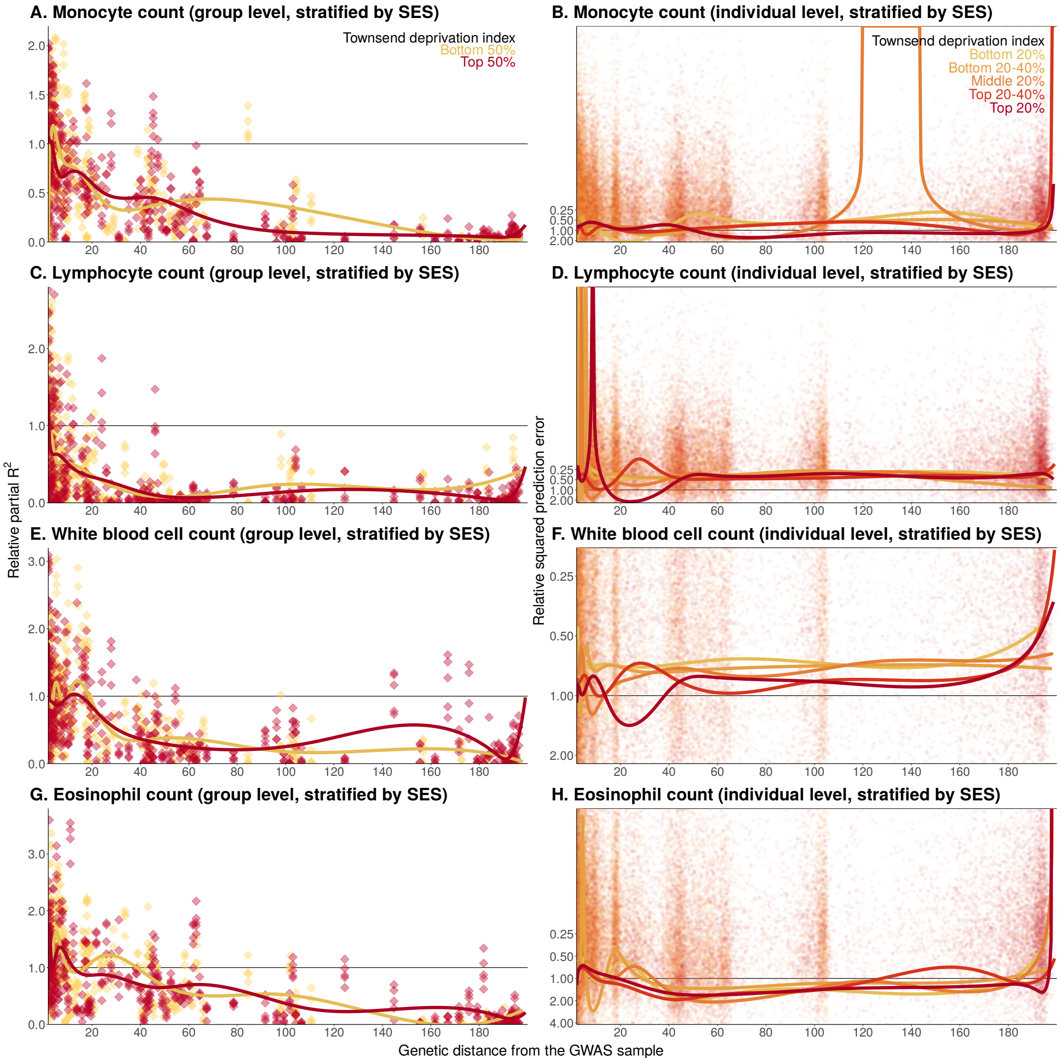

Figure S68: Portability as a function of both genetic distance and Townsend Deprivation Index, for white blood-cell related traits. We first stratified the prediction sample individuals into equal-sized strata of Townsend Deprivation Index: 2 strata at the group level, shown on the left, and 5 equal-sized strata at the individual level, shown on the right panels. Within each stratum, we then performed the same analysis of prediction accuracy as a function of genetic distance as in in **Figs. 2, S2-S5**. At the group level, we measured prediction accuracy with the squared partial correlation between the PGS and the trait value in 250 bins of 278 individuals each. At the individual level, we measured prediction accuracy as the squared difference between the predicted phenotype and the true phenotype value. At both the group and individual levels, y-axis values show relative prediction accuracy, i.e. prediction accuracy divided by a baseline value. The baseline value is the mean prediction accuracy in 25 bins with median genetic distances that are closest to the mean genetic distance for GWAS individuals. Curves show cubic spline fits, with 8 knots placed based on the density of data points. "Top" represents those who are more deprived. For **B, D**, when plotting the spline fit on a logarithmic scale, we replaced the non-positive values with $10^{-6}$.

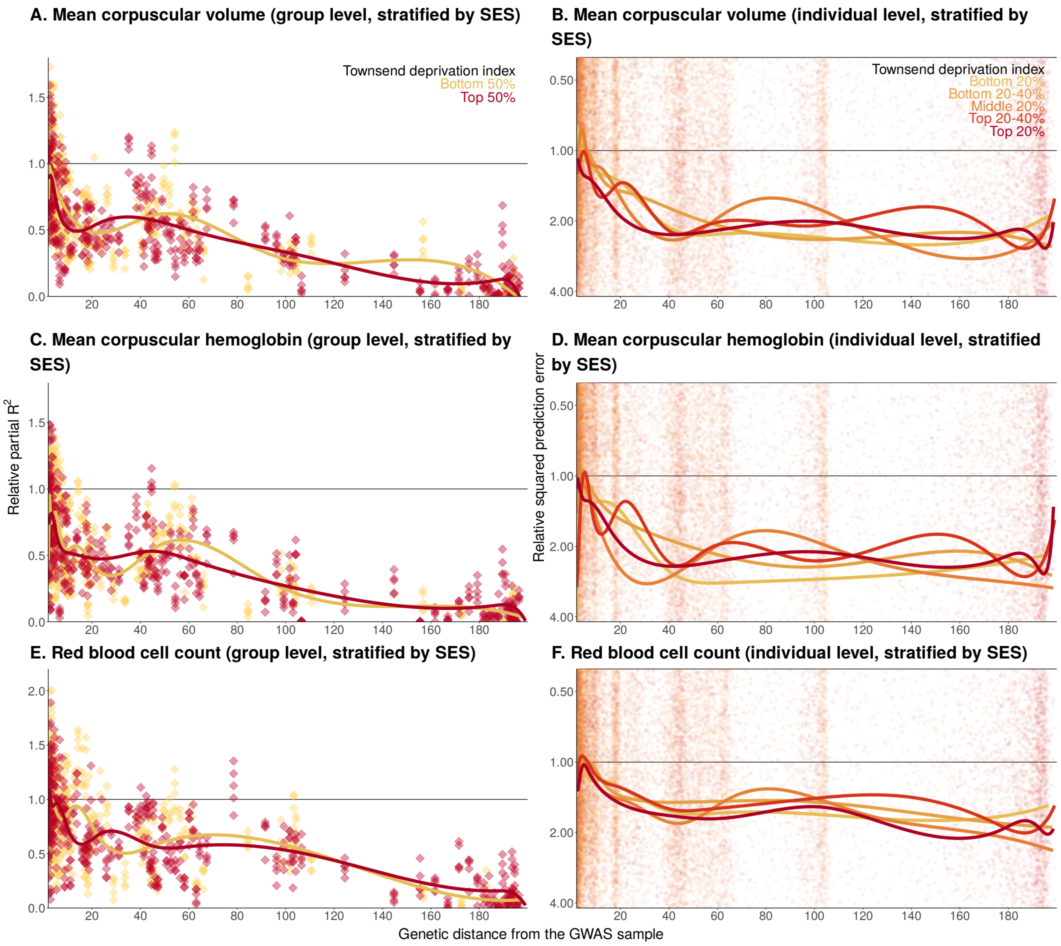

Figure S69: Portability as a function of both genetic distance and Townsend Deprivation Index, for red blood cell-related traits. We first stratified the prediction sample individuals into equal-sized strata of Townsend Deprivation Index: 2 strata at the group level, shown on the left, and 5 equal-sized strata at the individual level, shown on the right panels. Within each stratum, we then performed the same analysis of prediction accuracy as a function of genetic distance as in in **Figs. 2, S2-S5**. At the group level, we measured prediction accuracy with the squared partial correlation between the PGS and the trait value in 250 bins of 278 individuals each. At the individual level, we measured prediction accuracy as the squared difference between the predicted phenotype and the true phenotype value. At both the group and individual levels, y-axis values show relative prediction accuracy, i.e. prediction accuracy divided by a baseline value. The baseline value is the mean prediction accuracy in 25 bins with median genetic distances that are closest to the mean genetic distance for GWAS individuals. Curves show cubic spline fits, with 8 knots placed based on the density of data points. "Top" represents those who are more deprived.

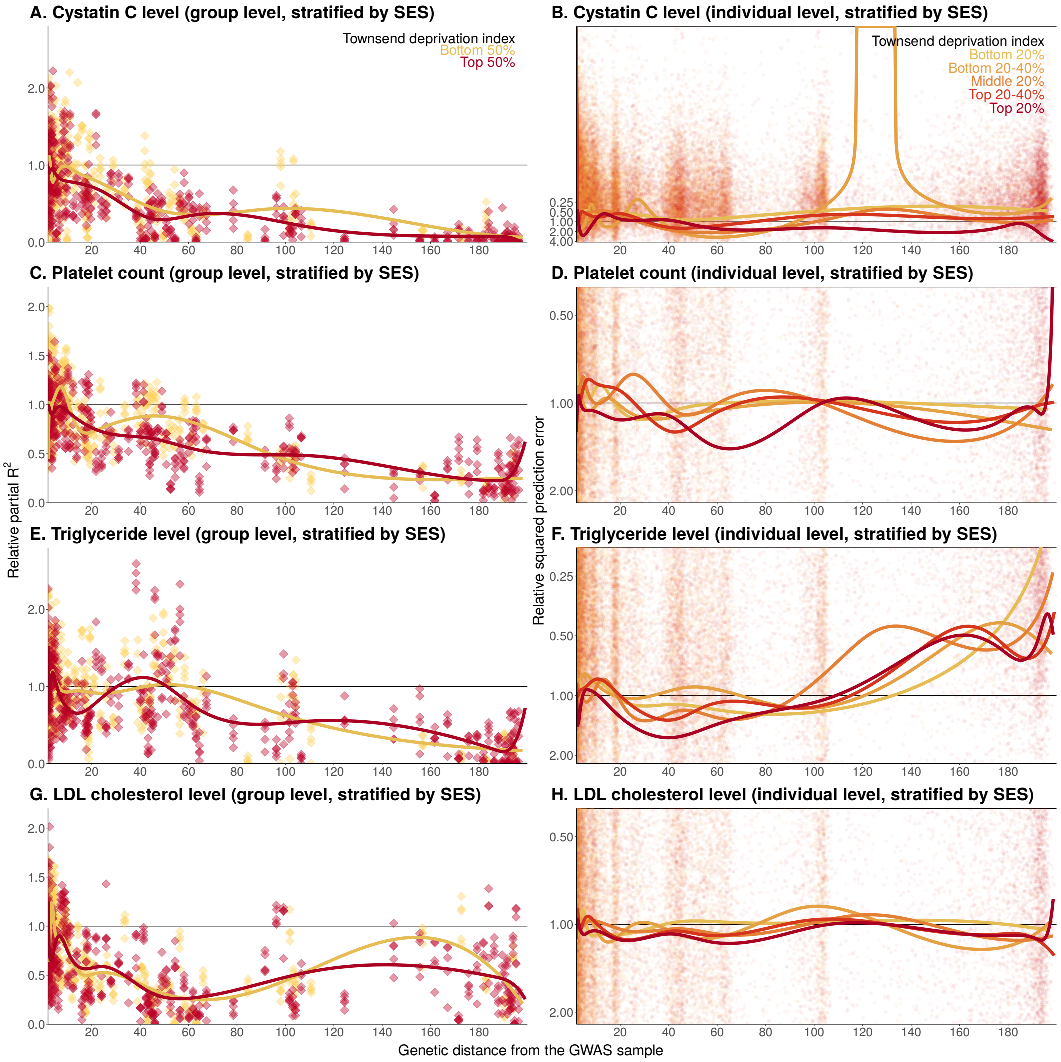

Figure S70: Portability as a function of both genetic distance and Townsend Deprivation Index, for other biomarkers. We first stratified the prediction sample individuals into equal-sized strata of Townsend Deprivation Index: 2 strata at the group level, shown on the left, and 5 equal-sized strata at the individual level, shown on the right panels. Within each stratum, we then performed the same analysis of prediction accuracy as a function of genetic distance as in in **Figs. 2, S2-S5**. At the group level, we measured prediction accuracy with the squared partial correlation between the PGS and the trait value in 250 bins of 278 individuals each. At the individual level, we measured prediction accuracy as the squared difference between the predicted phenotype and the true phenotype value. At both the group and individual levels, y-axis values show relative prediction accuracy, i.e. prediction accuracy divided by a baseline value. The baseline value is the mean prediction accuracy in 25 bins with median genetic distances that are closest to the mean genetic distance for GWAS individuals. Curves show cubic spline fits, with 8 knots placed based on the density of data points. “Top” represents those who are more deprived. For **B**, when plotting the spline fit on a logarithmic scale, we replaced the non-positive values with $10^{-6}$.

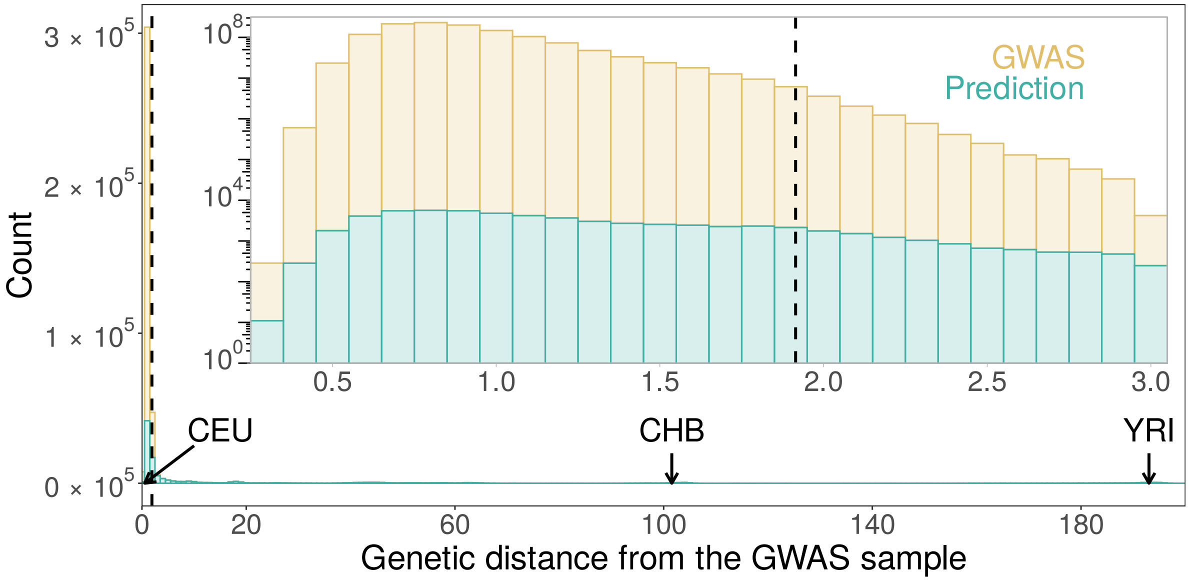

Figure S71: Measuring "genetic distance" from the GWAS sample, with White British individuals included in the prediction set. This figure mirrors main text **Fig. 1C**, but reflects different compositions of the GWAS sample and the prediction sample than the ones used for the main text analysis. Here, after filtering, from a total of 336,923 WB individuals, we randomly selected 300,000 as the GWAS set, and used the remaining 106,423 individuals as the prediction set (36,923 WB and 69,500 NWB). We followed the same procedures as described in **Methods**, and identified that across 10,000 individuals from the prediction set, genetic distance to the GWAS sample (calculated with 40 PCs) is highly correlated with $F_{st}$ between the GWAS sample and the individual. This figure is similar to **Figs. 1C**, showing the distribution of genetic distance, but to the new GWAS centroid. For reference, we show the mean genetic distances for subsets of the 1000 Genomes dataset[3]: CEU, Utah residents of primarily Northern and Western European descent; CHB, Han Chinese in Beijing, China; YRI, Yoruba in Ibadan, Nigeria. The dashed line represents the 97.5$^{\text{th}}$ percentile of genetic distance from among GWAS sample individuals. The inset is a zoomed-in view of genetic distances up to 3.0, on a log scale, to better visualize the distribution of the new GWAS sample.

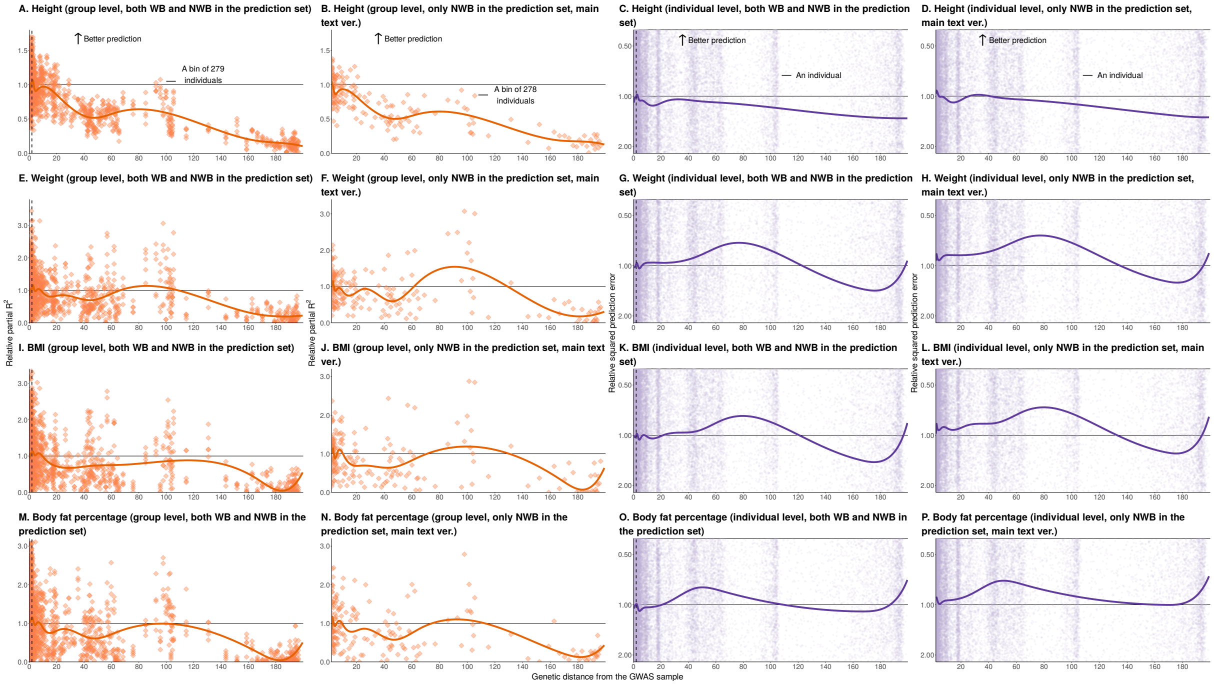

Figure S72: A comparison of portability trends with different choices of the GWAS and prediction samples, for anthropometric measurements. This figure presents the same analysis as **Fig. 2** and **Fig. S2**, but the first and third columns (**A, C, E, G, I, K, M, O**) show results obtained with a smaller GWAS set of 300,000 individuals labeled by the UKB as White British (WB) and a larger prediction set of 36,923 WB and 69,500 additional individuals not labeled as White British (NWB). The second and fourth columns (**B, D, F, H, J, L, N, P**) show results obtained with a GWAS set of 336,923 WB and a prediction set of 69,500 NWB only. At the group level (left two columns), we measured prediction accuracy with the squared partial correlation between the PGS and the trait value, in 382 bins of 279 individuals each in the first column and 250 bins of 278 individuals each in the second column. At the individual level (right two columns), we measured prediction accuracy as the squared difference between the predicted phenotype and the true phenotype value. At both the group and individual levels, y-axis values show relative prediction accuracy, i.e. prediction accuracy divided by a baseline value. The baseline value is the mean prediction accuracy in 38 bins with median genetic distances that are closest to the mean genetic distance for GWAS individuals. Curves show cubic spline fits, with 8 knots placed based on the density of data points.

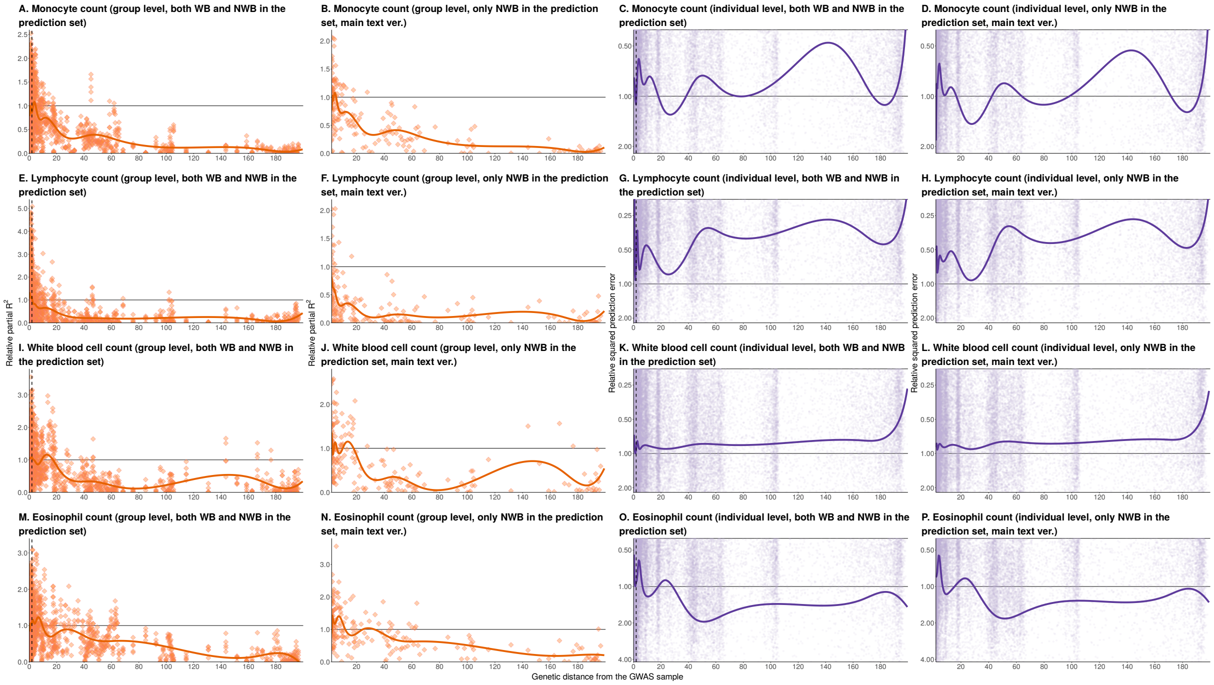

Figure S73: A comparison of portability trends with different choices of the GWAS and prediction samples, for white blood-cell related traits. This figure presents the same analysis as **Fig. 2** and **Fig. S2**, but the first and third columns (**A, C, E, G, I, K, M, O**) show results obtained with a smaller GWAS set of 300,000 individuals labeled by the UKB as White British (WB) and a larger prediction set of 36,923 WB and 69,500 additional individuals not labeled as White British (NWB). The second and fourth columns (**B, D, F, H, J, L, N, P**) show results obtained with a GWAS set of 336,923 WB and a prediction set of 69,500 NWB only. At the group level (left two columns), we measured prediction accuracy with the squared partial correlation between the PGS and the trait value, in 382 bins of 279 individuals each in the first column and 250 bins of 278 individuals each in the second column. At the individual level (right two columns), we measured prediction accuracy as the squared difference between the predicted phenotype and the true phenotype value. At both the group and individual levels, y-axis values show relative prediction accuracy, i.e. prediction accuracy divided by a baseline value. The baseline value is the mean prediction accuracy in 38 bins with median genetic distances that are closest to the mean genetic distance for GWAS individuals. Curves show cubic spline fits, with 8 knots placed based on the density of data points.

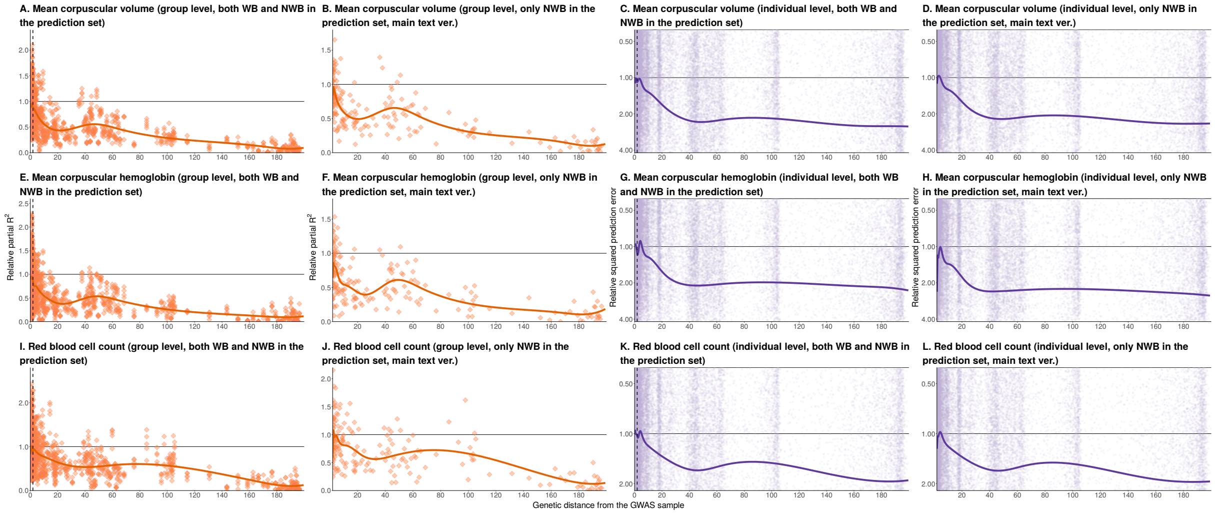

Figure S74: A comparison of portability trends with different choices of the GWAS and prediction samples, for red blood cell-related traits. This figure presents the same analysis as **Fig. 2** and **Fig. S2**, but the first and third columns (**A, C, E, G, I, K**) show results obtained with a smaller GWAS set of 300,000 individuals labeled by the UKB as White British (WB) and a larger prediction set of 36,923 WB and 69,500 additional individuals not labeled as White British (NWB). The second and fourth columns (**B, D, F, H, J, L**) show results obtained with a GWAS set of 336,923 WB and a prediction set of 69,500 NWB only. At the group level (left two columns), we measured prediction accuracy with the squared partial correlation between the PGS and the trait value, in 382 bins of 279 individuals each in the first column and 250 bins of 278 individuals each in the second column. At the individual level (right two columns), we measured prediction accuracy as the squared difference between the predicted phenotype and the true phenotype value. At both the group and individual levels, y-axis values show relative prediction accuracy, i.e. prediction accuracy divided by a baseline value. The baseline value is the mean prediction accuracy in 38 bins with median genetic distances that are closest to the mean genetic distance for GWAS individuals. Curves show cubic spline fits, with 8 knots placed based on the density of data points.

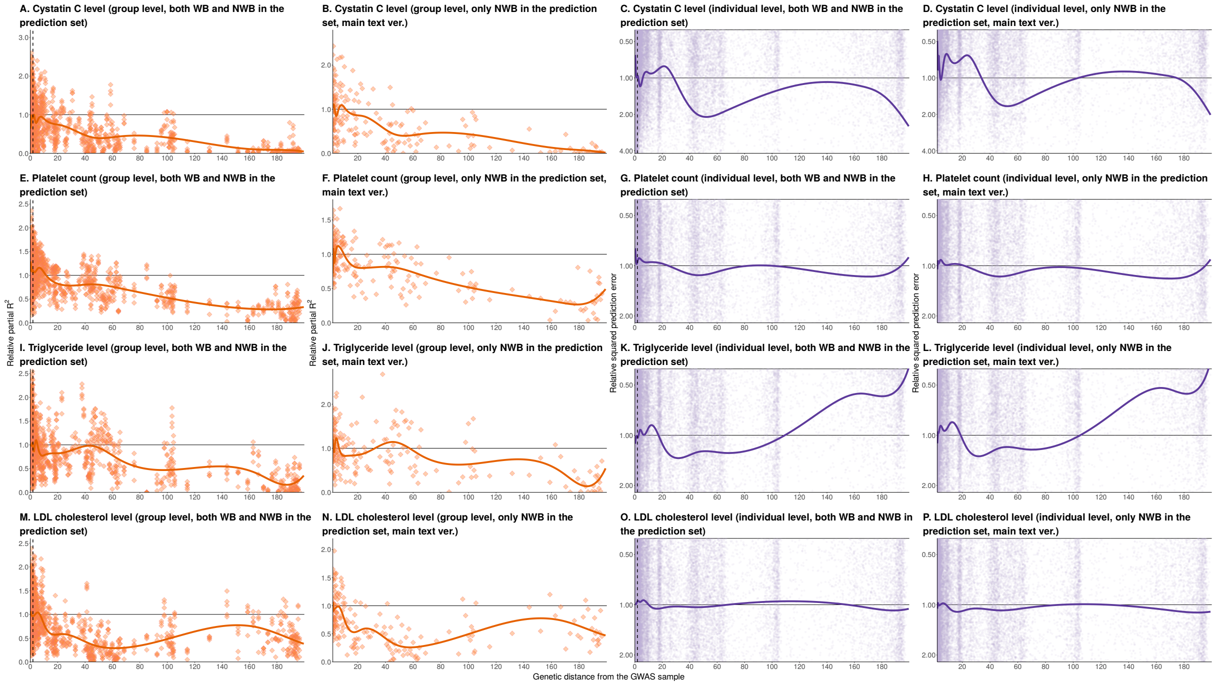

Figure S75: A comparison of portability trends with different choices of the GWAS and prediction samples, for other biomarkers. This figure presents the same analysis as **Fig. 2** and **Fig. S2**, but the first and third columns (**A, C, E, G, I, K, M, O**) show results obtained with a smaller GWAS set of 300,000 individuals labeled by the UKB as White British (WB) and a larger prediction set of 36,923 WB and 69,500 additional individuals not labeled as White British (NWB). The second and fourth columns (**B, D, F, H, J, L, N, P**) show results obtained with a GWAS set of 336,923 WB and a prediction set of 69,500 NWB only. At the group level (left two columns), we measured prediction accuracy with the squared partial correlation between the PGS and the trait value, in 382 bins of 279 individuals each in the first column and 250 bins of 278 individuals each in the second column. At the individual level (right two columns), we measured prediction accuracy as the squared difference between the predicted phenotype and the true phenotype value. At both the group and individual levels, y-axis values show relative prediction accuracy, i.e. prediction accuracy divided by a baseline value. The baseline value is the mean prediction accuracy in 38 bins with median genetic distances that are closest to the mean genetic distance for GWAS individuals. Curves show cubic spline fits, with 8 knots placed based on the density of data points.

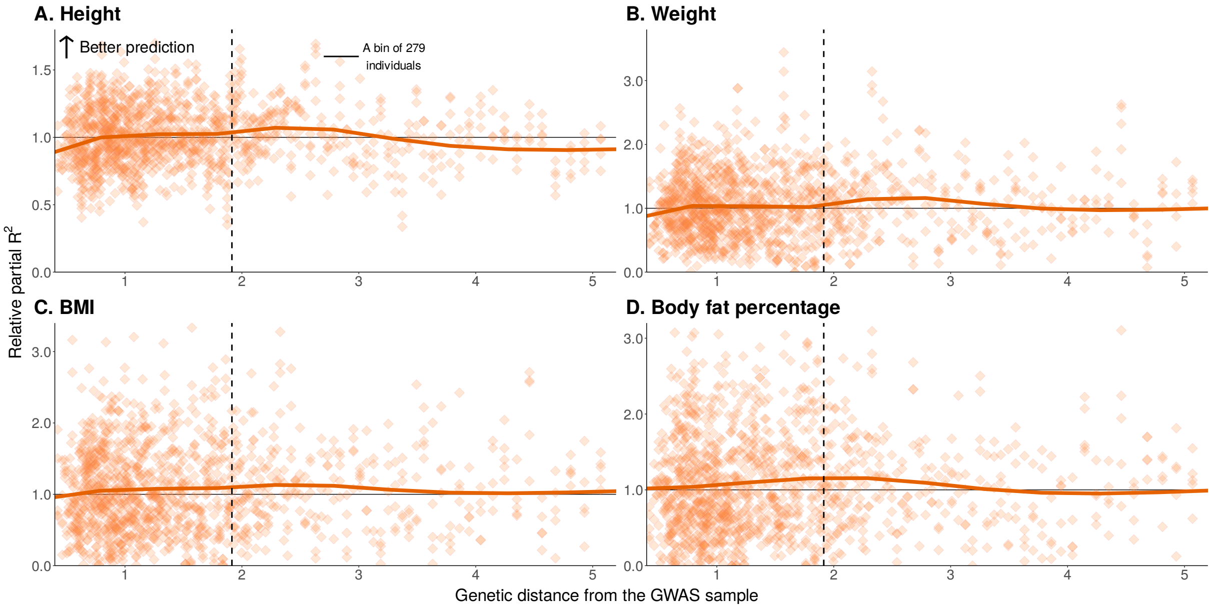

Figure S76: Zoomed in portability trends, with WB in the prediction set, for anthropometric measurements. This figure is a zoomed-in version of **Fig. S72A,E,I,M**, to better visualize the bins at small genetic distances. The results are obtained from a GWAS set of 300,000 WB and a prediction set of both WB and NWB (36,923 WB and 69,500 NWB). We measured prediction accuracy with the squared partial correlation between the PGS and the trait value, in 382 bins of 279 individuals each. Y-axis values show relative prediction accuracy, i.e. prediction accuracy divided by a baseline value. The baseline value is the mean prediction accuracy in 38 bins with median genetic distances that are closest to the mean genetic distance for GWAS individuals. Curves show cubic spline fits, with 8 knots placed based on the density of data points. The x-axis is zoomed in to up to 5.0.

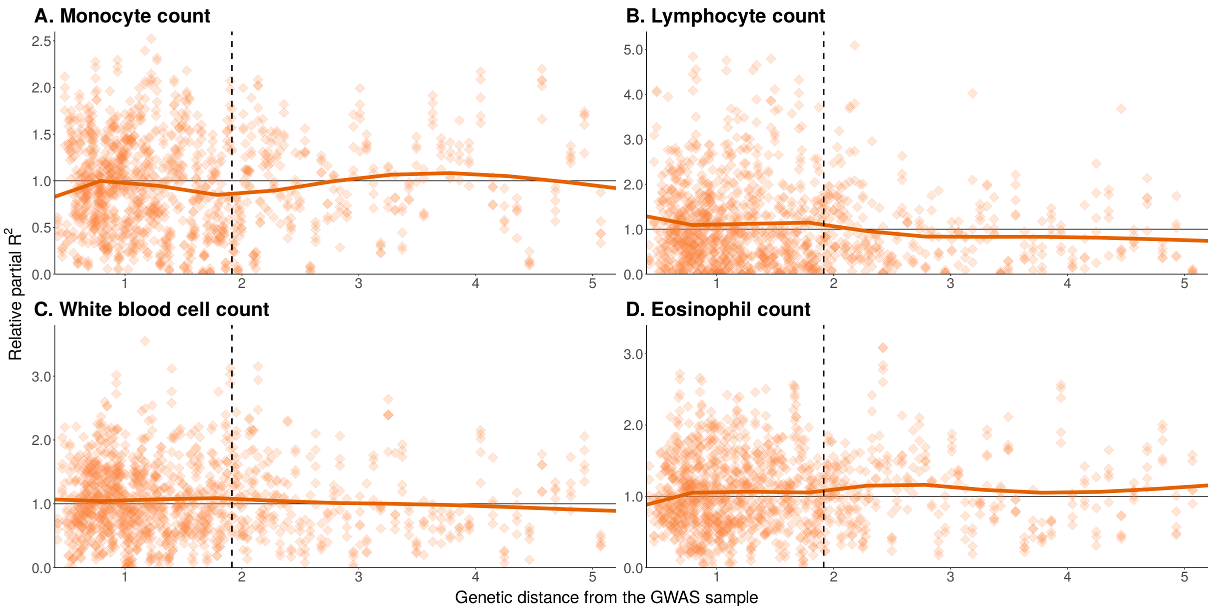

Figure S77: Zoomed in portability trends, with WB in the prediction set, for white blood-cell related traits. This figure is a zoomed-in version of **Fig. S73A,E,I,M**, to better visualize the bins at small genetic distances. The results are obtained from a GWAS set of 300,000 WB and a prediction set of both WB and NWB (36,923 WB and 69,500 NWB). We measured prediction accuracy with the squared partial correlation between the PGS and the trait value, in 382 bins of 279 individuals each. Y-axis values show relative prediction accuracy, i.e. prediction accuracy divided by a baseline value. The baseline value is the mean prediction accuracy in 38 bins with median genetic distances that are closest to the mean genetic distance for GWAS individuals. Curves show cubic spline fits, with 8 knots placed based on the density of data points. The x-axis is zoomed in to up to 5.0.

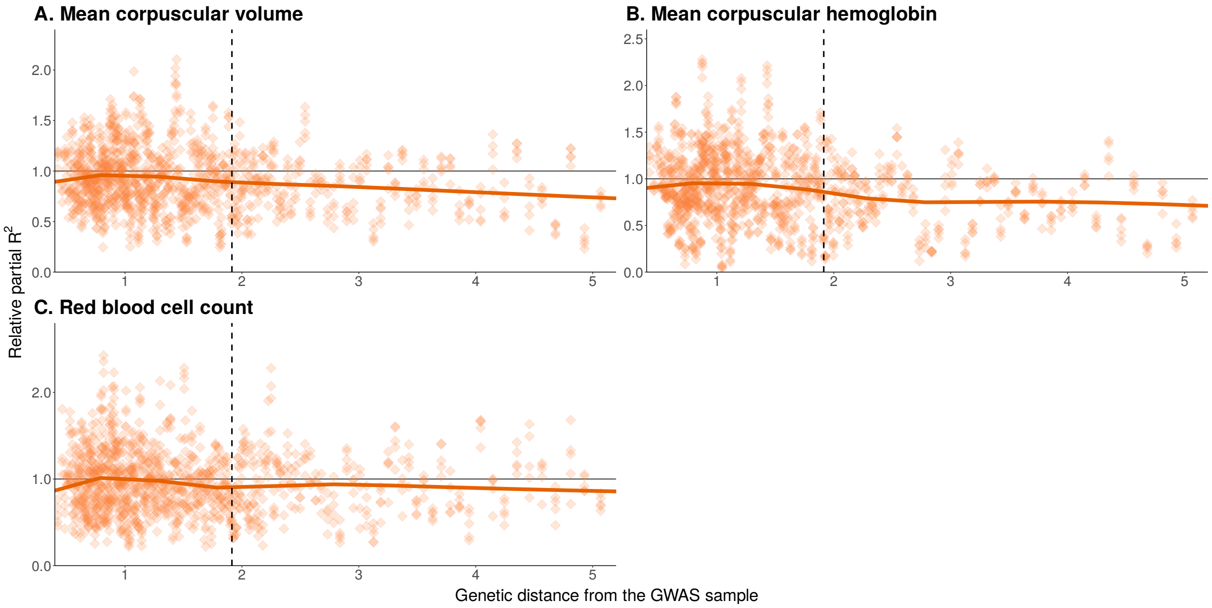

Figure S78: Zoomed in portability trends, with WB in the prediction set, for red blood cell-related traits. This figure is a zoomed-in version of **Fig. S74A,E,I**, to better visualize the bins at small genetic distances. The results are obtained from a GWAS set of 300,000 WB and a prediction set of both WB and NWB (36,923 WB and 69,500 NWB). We measured prediction accuracy with the squared partial correlation between the PGS and the trait value, in 382 bins of 279 individuals each. Y-axis values show relative prediction accuracy, i.e. prediction accuracy divided by a baseline value. The baseline value is the mean prediction accuracy in 38 bins with median genetic distances that are closest to the mean genetic distance for GWAS individuals. Curves show cubic spline fits, with 8 knots placed based on the density of data points. The x-axis is zoomed in to up to 5.0.

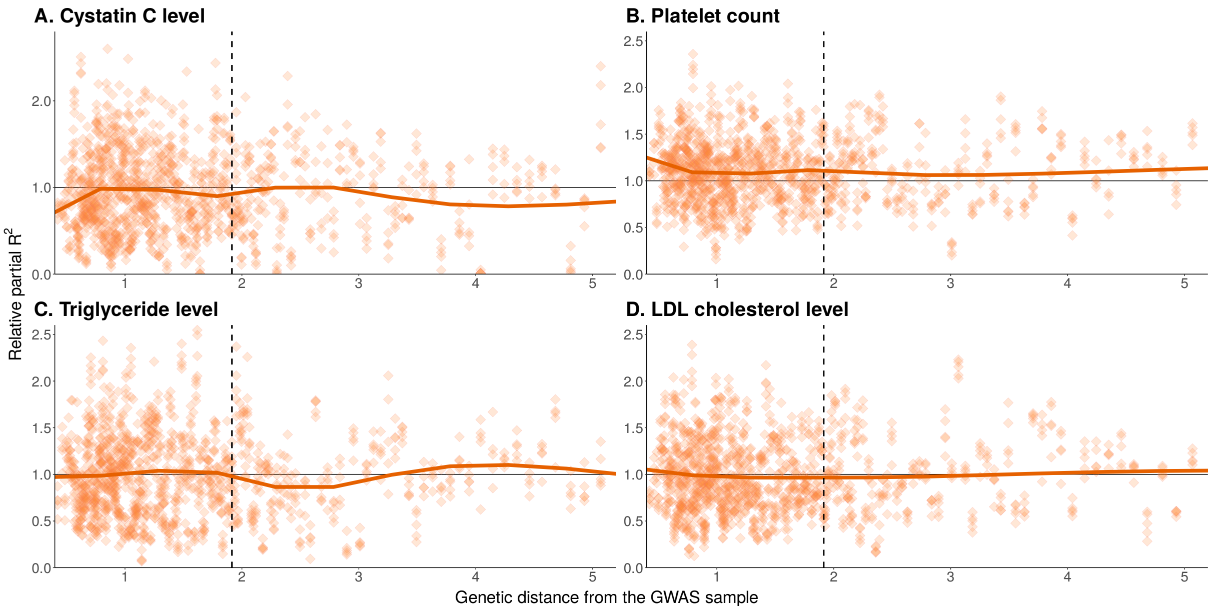

Figure S79: Zoomed in portability trends, with WB in the prediction set, for other biomarkers. This figure is a zoomed-in version of **Fig. S75A,E,I,M**, to better visualize the bins at small genetic distances. The results are obtained from a GWAS set of 300,000 WB and a prediction set of both WB and NWB (36,923 WB and 69,500 NWB). We measured prediction accuracy with the squared partial correlation between the PGS and the trait value, in 382 bins of 279 individuals each. Y-axis values show relative prediction accuracy, i.e. prediction accuracy divided by a baseline value. The baseline value is the mean prediction accuracy in 38 bins with median genetic distances that are closest to the mean genetic distance for GWAS individuals. Curves show cubic spline fits, with 8 knots placed based on the density of data points. The x-axis is zoomed in to up to 5.0.
